# *Drosophila* SPG12 ortholog, reticulon-like 1, governs presynaptic ER organization and Ca^2+^ dynamics

**DOI:** 10.1101/2021.12.17.473229

**Authors:** Juan José Pérez-Moreno, Rebecca C Smith, Megan K Oliva, Filomena Gallo, Shainy Ojha, Karin H. Müller, Cahir J O’Kane

## Abstract

Neuronal endoplasmic reticulum (ER) appears continuous throughout the cell. Its shape and continuity are influenced by ER-shaping proteins, mutations in which can cause distal axon degeneration in Hereditary Spastic Paraplegia (HSP). We therefore asked how loss of *Rtnl1*, a *Drosophila* ortholog of the human HSP gene *RTN2* (*SPG12*), which encodes an ER-shaping protein, affects ER organization and the function of presynaptic terminals. Loss of Rtnl1 depleted ER membrane markers at *Drosophila* presynaptic motor terminals, and appeared to deplete narrow tubular ER while leaving cisternae largely unaffected, thus suggesting little change in resting Ca^2+^ storage capacity. Nevertheless, these changes were accompanied by major reductions in activity-evoked Ca^2+^ fluxes in the cytosol, ER lumen, and mitochondria, as well as reduced evoked and spontaneous neurotransmission. We found that reduced STIM-mediated ER-plasma membrane contacts underlie presynaptic Ca^2+^ defects in *Rtnl1* mutants. Our results show the importance of ER architecture in presynaptic physiology and function which are therefore potential factors in the pathology of HSP.

## INTRODUCTION

Synapses, the computational units of nervous systems, are immensely adaptable. Endoplasmic reticulum (ER) is a site of some of the processes that tune synaptic strength. It is a major Ca^2+^ store of the cell, a site of lipid biosynthesis, and can regulate the physiology of other organelles through contact sites that support exchange of Ca^2+^ and lipids. Axonal ER comprises an extensive continuous network formed mainly of tubules (Wu et al., 2017) that are shaped by proteins of the reticulon and REEP families, among others. These contain intramembrane hairpin domains that can curve ER membrane (Shibata et al., 2009; Voeltz et al., 2006); loss of these proteins can reduce tubule number and curvature of axonal ER, and disrupt network continuity (O’Sullivan et al., 2012; Yalçın et al., 2017).

The ER network appears to influence axon survival and maintenance; mutations affecting ER-shaping proteins can lead to the axon degeneration disease, Hereditary Spastic Paraplegia (HSP) (Öztürk et al., 2020). HSPs show degeneration of the distal regions of longer upper motor axons, consistent with “dying back” degeneration from axon terminals, preferentially in the longest upper motor axons. The links between HSP and ER-shaping proteins suggest a critical relationship between ER architecture, and axonal and presynaptic maintenance. However, we lack a good model for how ER spatial organization affects axonal or presynaptic function and degeneration. One way in which ER might do this is as an intracellular Ca^2+^ store. ER Ca^2+^ stores can either contribute to presynaptic cytosolic Ca^2+^ responses, or sequester excess Ca^2+^ from the cytosol in response to high frequency stimulation (Chanaday et al., 2021; de Juan-Sanz et al., 2017; Kuromi & Kidokoro, 2002; Lindhout et al., 2019; Sanyal et al., 2005; Singh et al., 2021; Tran & Stricker, 2021). ER might potentially also influence presynaptic physiology via its interactions with mitochondria. Like ER, mitochondria can also take up Ca^2+^ when cytosolic Ca^2+^ is elevated, although this process appears unable to buffer Ca^2+^ sufficiently to affect synaptic transmission (Chouhan et al., 2010; Verstreken et al., 2005); rather its role appears to be to stimulate ATP synthesis in response to the Ca^2+^ influx that occurs on synaptic activity (Chouhan et al., 2012; Datta & Jaiswal, 2021; Jouaville et al., 1999). The cytosol, rather than ER, seems to be the main source of Ca^2+^ for mitochondria in presynaptic terminals (Ashrafi et al., 2020; Chouhan et al., 2012).

As well as being a bulk Ca^2+^ source or sink, the ER dynamically responds to lumenal Ca^2+^ levels. STIM (stromal interacting molecule) proteins in the ER membrane, when activated by depletion of ER Ca^2+^ stores, interact with effectors in the plasma membrane (PM), such as the Orai1 Ca^2+^ channel, leading to influx of extracellular Ca^2+^ into the cytosol (store-operated Ca^2+^ entry, SOCE) and refilling of ER stores (Serwach & Gruszczynska-Biegala, 2020). In primary hippocampal neurons, evoked neurotransmitter release correlates with ER Ca^2+^ content, due to a feedback loop dependent on inhibition of voltage-gated Ca^2+^ channels in the plasma membrane by activated STIM1 (de Juan-Sanz et al., 2017). Also in primary hippocampal neurons, SOCE, in a STIM2-dependent manner, can transiently increase presynaptic Ca^2+^ levels, and robustly augment spontaneous glutamate vesicular release (Chanaday et al., 2021).

Since mutations in several ER-shaping proteins can cause axon degeneration, altered ER architecture may elicit degenerative changes in axonal or presynaptic function. One mechanism could be local dysregulation of Ca^2+^ - for example through reduced availability of Ca^2+^ for synaptic signaling or capacity to remove cytosolic Ca^2+^, or interference with regulatory effects of the ER STIM Ca^2+^ sensors (Chanaday et al., 2021; de Juan-Sanz et al., 2017). Presynaptic ER comprises a network of interconnected tubules with occasional cisternae (Wu et al., 2017). While the presence of cisternae might indicate regions specialized in storing Ca^2+^, the predominance of narrow ER tubules suggests a local regulatory role for this organelle. To test how spatial organization of ER could impair synaptic function, we used *Drosophila* to study the effects of removing the tubular ER-shaping protein Rtnl1 on presynaptic ER organization, and on the Ca^2+^ fluxes between compartments during synaptic activity. *Rtnl1* is one of two documented reticulons in *Drosophila*, orthologous to the four RTNs found in mammals (RTN1-4), including the HSP causative gene RTN2 (Montenegro et al., 2012). It is distributed continuously in motor axons all the way to presynaptic termini (O’Sullivan et al., 2012), and its loss leads to partial depletion of axonal ER (O’Sullivan et al., 2012; Yalçın et al., 2017). Loss of Rtnl1 greatly decreases Excitatory Junction Potential (EJP) amplitude at the larval neuromuscular junction (NMJ) (Summerville et al., 2016), implying major effects on presynaptic physiology, although its effects on presynaptic ER and its role in presynaptic physiology has not been well defined.

Here we reveal how loss of Rtnl1 depletes tubular ER but not cisternae or ER volume at NMJs, and how this loss significantly decreases ER, mitochondrial, and cytosolic evoked Ca^2+^ fluxes. We find that this impact of Rtnl1 loss on presynaptic Ca^2+^ handling is caused by a decrease of STIM-mediated ER-PM contacts. Our results suggest a feedback loop, whereby tubular ER and inactivated STIM control evoked Ca^2+^ entry into the presynaptic compartment and neurotransmitter release.

While Rtnl1 loss reduces axonal ER levels mainly in longer motor axons (O’Sullivan et al., 2012; Yalçın et al., 2017), we observe that it reduces presynaptic tubular ER independently of axonal length. This phenotype provides a unique model to study the contribution of the presynaptic tubular ER network and ER-shaping HSP proteins to presynaptic Ca^2+^ handling. Our findings display a local role for presynaptic ER tubules, and how synaptic dysfunction might be a consequence of impairing functions of proteins that are mutated in HSP.

## RESULTS

### *Rtnl1*^*18*^ is a null mutant allele

To understand how Rtnl1 contributes to presynaptic ER organization and physiology, we generated new mutant alleles of *Rtnl1*. We had previously analyzed a loss-of-function allele, *Rtnl1*^*1*^ (Wakefield & Tear, 2006), with an internal deletion of around 5 kb (Yalçın et al., 2017) and obtained ER phenotypes that were broadly consistent with those of RNAi knockdown, but obtained only partial rescue using a genomic *Rtnl1* construct (O’Sullivan et al., 2012; Yalçın et al., 2017). Since additional alleles could allow us to exclude phenotypes due to second-site mutations, we generated new CRISPR/Cas9 mutant *Rtnl1* alleles. We targeted these using two *gRNAs*, each just upstream of the sequence encoding each intramembrane domain, present in all splice variants (Fig. 1; Supplementary Fig. S1). We characterized three new alleles; one of these, *Rtnl1*^*18*^, had multiple changes: an inversion of the region between both *gRNA* sequences, a 6-bp deletion at the site of the upstream *gRNA*, and a 19-bp deletion at the site of the downstream *gRNA* (Fig. 1; Supplementary Fig. S1A). This lesion inverts the sequence encoding the first intramembrane domain; and it deletes some of the sequence encoding the second intramembrane domain, leaving only the C-terminal half of this domain potentially able to be translated from an in-frame *AUG* codon (Supplementary Fig. S1B). While *Rtnl1*^*18*^ does not abolish *Rtnl1* transcription (Supplementary Fig. S2), it is likely to be a null allele, since the rearrangements downstream of the first *gRNA* site make it impossible to express more than a fragment of the protein, which lacks an intact intramembrane domain.

**Figure 1.**
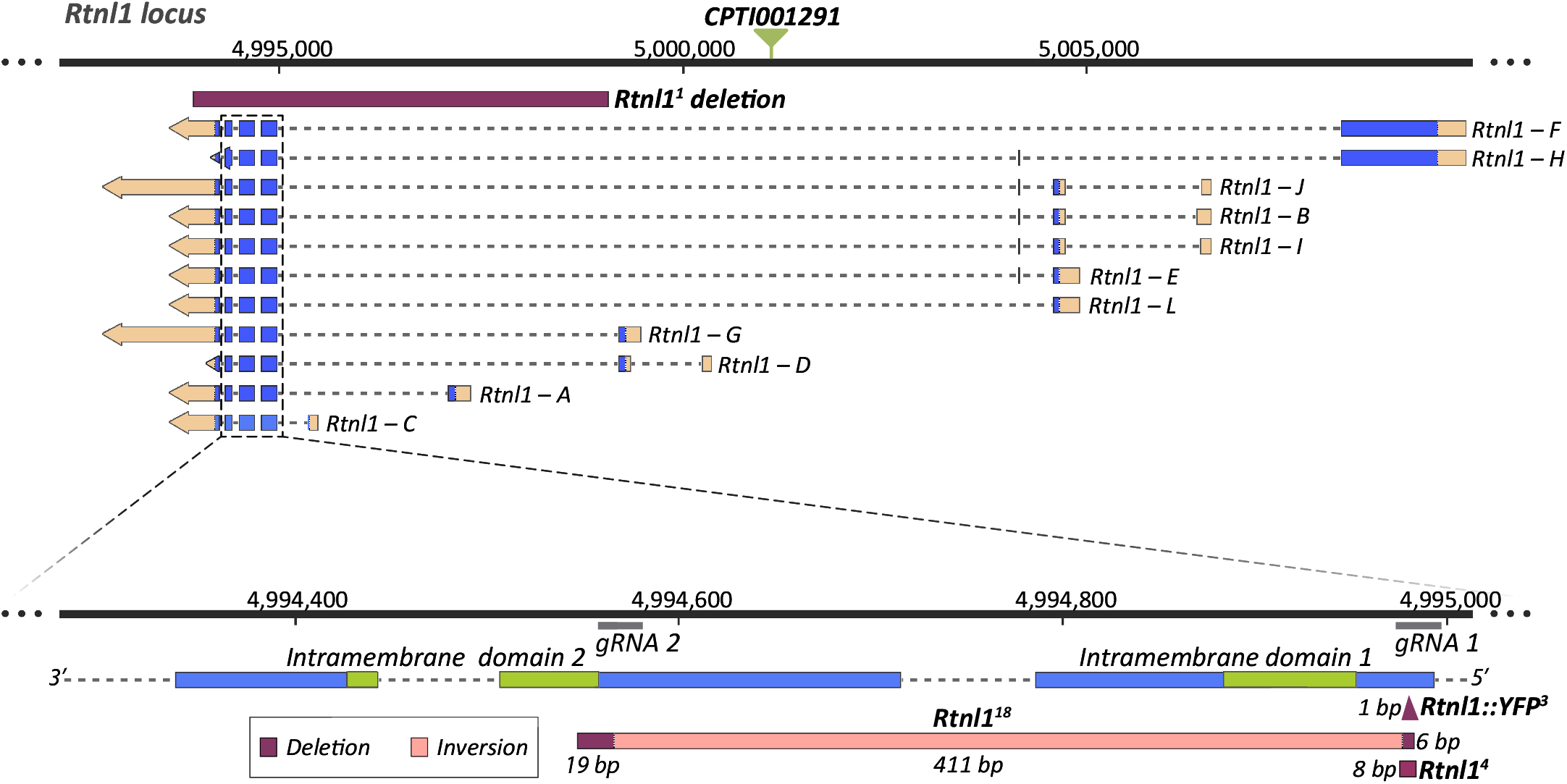
Generation of new *Rtnl1* null mutant alleles. For the different *Rtnl1* splice variants, blue boxes indicate coding sequence, broken lines introns, light orange boxes UTR sequences, and arrows the direction of transcription. Green triangle indicates the position of the *Rtnl1::YFP* exon trap insertion (*CPTI001291*). Genomic coordinates are from Release 6.26 of the *Drosophila melanogaster* genome (http://flybase.org). Magnified detail shows exons encoding Rtnl1 intramembrane domains (green; bottom), locations recognized by *gRNAs*, the *Rtnl1*^*18*^ lesion, and the 8-bp (*Rtnl1*^*4*^) and the 1-bp (*Rtnl1::YFP*^*3*^) frameshift deletions, generated at the site of the upstream *gRNA* on *Rtnl1*^*+*^ and *Rtnl1*^*CPTI001291*^ backgrounds respectively, but not used further in this work. The length of each mutated region is shown. In *Rtnl1*^*18*^, the region encoding the first intramembrane domain is completely disrupted by an inversion, while the initial part of the sequence encoding the second intramembrane domain is absent due to a deletion and frameshift (see **Supp. Fig. 1** for more molecular details of alleles).

### Rtnl1 loss-of-function decreases presynaptic ER membrane labeling

We next tested *Rtnl1* mutant NMJs for altered ER distribution. We expressed tdTom::Sec61β, which labels presynaptic ER (Summerville et al., 2016), using *GAL4* drivers specific for each main type of larval excitatory motor neuron (MN), Ib and Is (Pérez-Moreno & O’Kane, 2019), which differ structurally (Menon et al., 2013) and physiologically (Aponte-Santiago et al., 2020; Aponte-Santiago & Littleton, 2020; Chouhan et al., 2010). Compared with *wildtype* (*WT*) NMJs of the same genetic background, *Rtnl1*^*18*^ showed decreased tdTom::Sec61β levels in both Type Ib (Fig. 2A) and Is (Fig. 2B) NMJs. This decrease was not due to lowered *GAL4*-dependent expression; the plasma membrane marker, CD4::tdGFP (Han et al., 2011), expressed at NMJs using the same *GAL4* lines, was not affected by *Rtnl1*^*18*^. We found a similar reduction in tdTom::Sec61β using a general MN driver, *D42-GAL4* (Sanyal, 2009) in *Rtnl1*^*18*^ NMJs (Fig. 2C).

**Figure 2.**
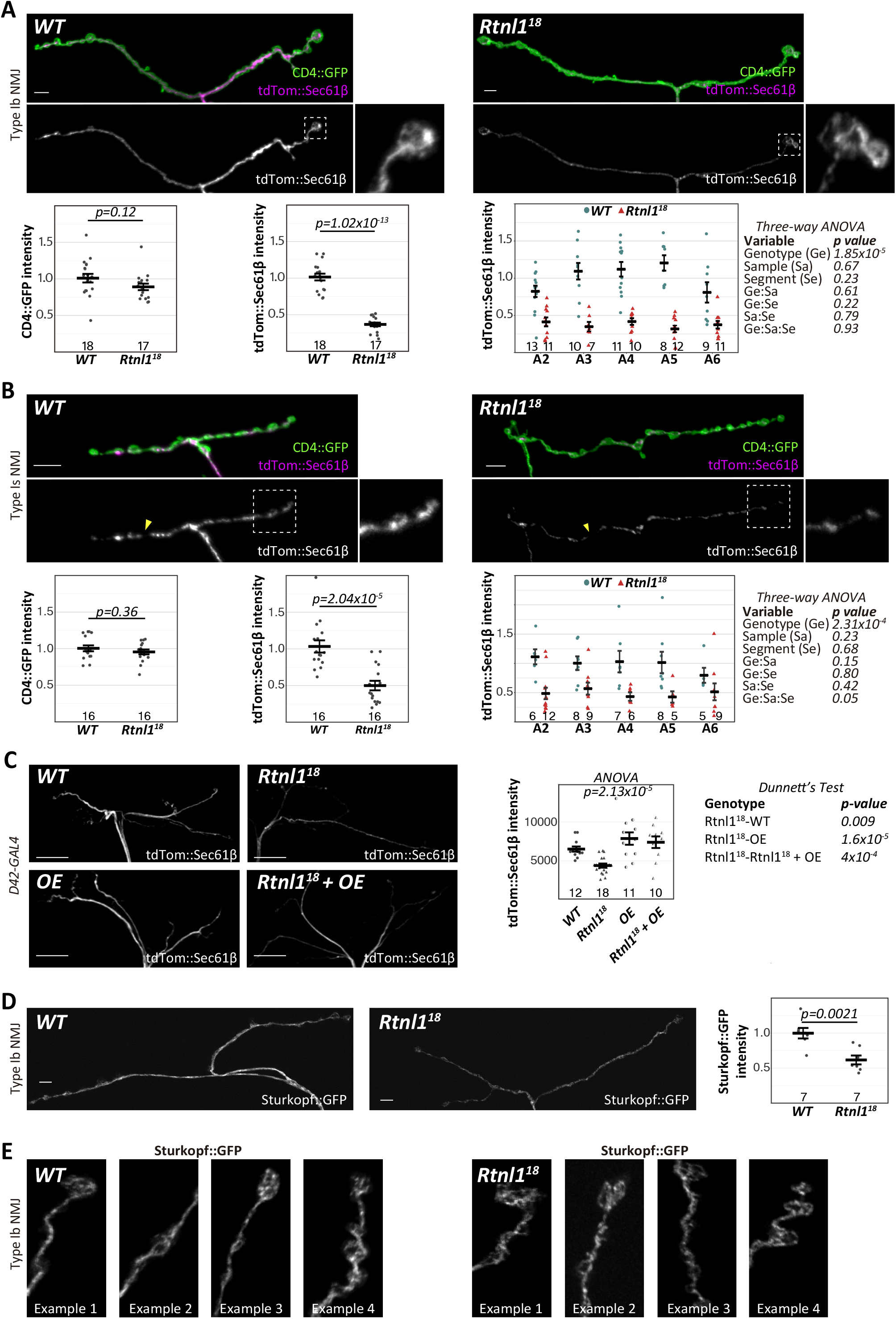
Rtnl1 loss depletes presynaptic ER membrane markers. (**A, B**) Representative confocal projections and quantifications of ER marker tdTom::Sec61β in (**A**) Type Ib terminals on muscle 1 and (**B**) Type Is terminals on muscles 1-9, of *WT* and *Rtnl1*^*18*^ larvae. Insets show magnified views of the areas inside broken lines. Arrowheads in **B** indicate gaps in tdTom::Sec61β signal. (**C**) Rtnl1::HA overexpression (OE) rescues the depletion of tdTom::Sec61β in *Rtnl1*^*18*^ mutant NMJs at muscle 1/9. Since Type Ib and Is *GAL4* constructs, *UAS-Rtnl1::HA*, and *UAS-tdTom::Sec61*β, are all inserted at *attP2*, we used *D42-GAL4* which could be recombined with the *UAS* insertions. (**D**) Sturkopf::GFP in Type Ib muscle 1 NMJs, with intensity analyzed as above, and (**E**) high magnification examples. All plots show individual larval datapoints and mean ± SEM; y-axes indicate arbitrary units (au) after normalization to *WT*; sample size is within the plots for each genotype. Where tdTom::Sec61β signal is compared between segments (**A** and **B**), sample size is the number of hemisegments (NMJs); for other plots, sample size is the number of larvae. For each larva, several NMJs across segments A2-A6 were randomly analysed, and the mean value used as a larval datapoint. For each larva in **C**, we analyzed (but did not distinguish) Type Ib and Is branches on muscles 1 and 9 in a single segment chosen randomly between A4-A6. Student’s t-tests were used for pairwise comparisons; ANOVA was used as shown for comparisons of more than two categories, or of multiple factors, with Dunnett’s post-hoc testing where appropriate. Scale bars for **A, B** and **D** are 5 μm, for **C** 20μm. Insets in **A** and **B** are 8×8 μm. Panel width in **E** is 10 μm. Genotypes: **A-B**, *GAL4, UAS-CD4::tdGFP / UAS-tdTom::Sec61*β; **C**, *D42-GAL4, UAS-tdTom::Sec61*β and *D42-GAL4, UAS-tdTom::Sec61*β */ UAS-Rtnl1::HA* in either *Rtnl1*^*+*^ (*WT*) or *Rtnl1*^*18*^ background (OE and *Rtnl1*^*18*^ + OE respectively); **D-E**, *Ib-GAL4, UAS-Sturkopf::GFP / +* ; all in either a *WT* or *Rtnl1*^*18*^ background.

Loss of *Rtnl1* reduces ER levels specifically in distal long motor axons (O’Sullivan et al., 2012; Yalçın et al., 2017); however, lowered presynaptic tdTom::Sec61β levels in *Rtnl1*^*18*^ NMJs were independent of motor axon length (Fig. 2A-B). *Rtnl1*^*18*^ presynaptic terminals still showed an interconnected and mostly continuous ER network (insets in Fig. 2A; Supplementary Fig. S3A), with a distribution resembling that of *WT*. These findings suggest that loss of Rtnl1 leads to less presynaptic ER membrane, but that its general organizational features are unaffected. The phenotype appears to affect ER membrane in general rather than tdTom::Sec61β in particular; a second ER membrane marker, Sturkopf::GFP (Thiel et al., 2013; Yalçın et al., 2017), also showed decreased levels in *Rtnl1*^*18*^ NMJs (Fig. 2D), and an apparently normal ER network organization (Fig. 2E; Supplementary Fig. S3B). Type Is terminals, but not Type Ib, showed occasional apparent gaps in the ER network (Fig. 2B) in both *WT* and *Rtnl1*^*18*^ NMJs, perhaps due to having a smaller ER network that is more sensitive to partial loss of tubules.

The partial depletion of ER membrane was specifically due to loss of Rtnl1. First, we also observed it at Type Ib and Type Is NMJs in a second mutant allele, *Rtnl1*^*1*^ (Supplementary Fig. S4A). Second, expression of *UAS-Rtnl1::HA* (Summerville et al., 2016) under GAL4 control fully rescued tdTom::Sec61β levels in *Rtnl1*^*18*^ NMJs (Fig. 2C). Surprisingly, a *UAS-Rtnl1::GFP* construct (Rao et al., 2016), appeared to be dominant negative; it produced a similar tdTom::Sec61β decrease as *Rtnl1* loss-of-function, and even made a *Rtnl1* loss-of-function phenotype slightly more severe (Supplementary Fig. S4B). Taken together, our results in a variety of genotypes and with two different markers support the conclusion that Rtnl1 controls presynaptic ER levels.

### *Rtnl1* loss-of-function specifically decreases presynaptic tubular ER network

The reduced levels of ER membrane markers in *Rtnl1* mutants contrasted with the unchanged resting fluorescence of the Ca^2+^ sensor GCaMP6-210 in the ER lumen, on loss of multiple ER-shaping proteins including Rtnl1 (Oliva et al., 2020). While the fluorescence of GCaMP6-210 is also Ca^2+^-dependent, we hypothesized that the role of reticulons in generating highly curved ER tubules (Voeltz et al., 2006) might explain the apparent discrepancy, by preferentially removing ER tubules compared to cisternae, due to their greater curvature, in *Rtnl1* mutant NMJs. Similar to mammalian neurons (Wu et al., 2017), we observed both ER tubules with a lumen too small to be visible (as reported previously in axons (Terasaki, 2018; Yalçın et al., 2017)) and cisternae with a larger lumen, in presynaptic motor terminals, using electron microscopy (Fig. 3A). Tubules were observed most clearly in regions of the bouton where synaptic vesicles (SVs) were sparse, but harder to distinguish from SVs in regions of the bouton that were rich in SVs due to the similar diameters of ER tubules and SVs, thus preventing reconstruction of the whole ER network. Using the ER lumen marker BiP::sfGFP::HDEL (Summerville et al., 2016), hereafter referred to as GFP::HDEL, we observed larger and brighter puncta that we interpret as cisternae with a large lumen, joined by stretches of fainter signal that we interpret as ER tubules with a narrow lumen, that we observed using electron microscopy (Fig. 3B-D). These bright puncta often, but not always, overlap with puncta of tdTom::Sec61β (Supplementary Fig. S5A), suggesting that enrichment of these two markers highlights different features of ER that are sometimes found together - indeed, electron microscopy of axonal ER shows both cisternae (with a noticeable lumen) and small sheets (without a noticeable lumen) that are sometimes, but not always, apposed to each other (Yalçın et al., 2017).

**Figure 3.**
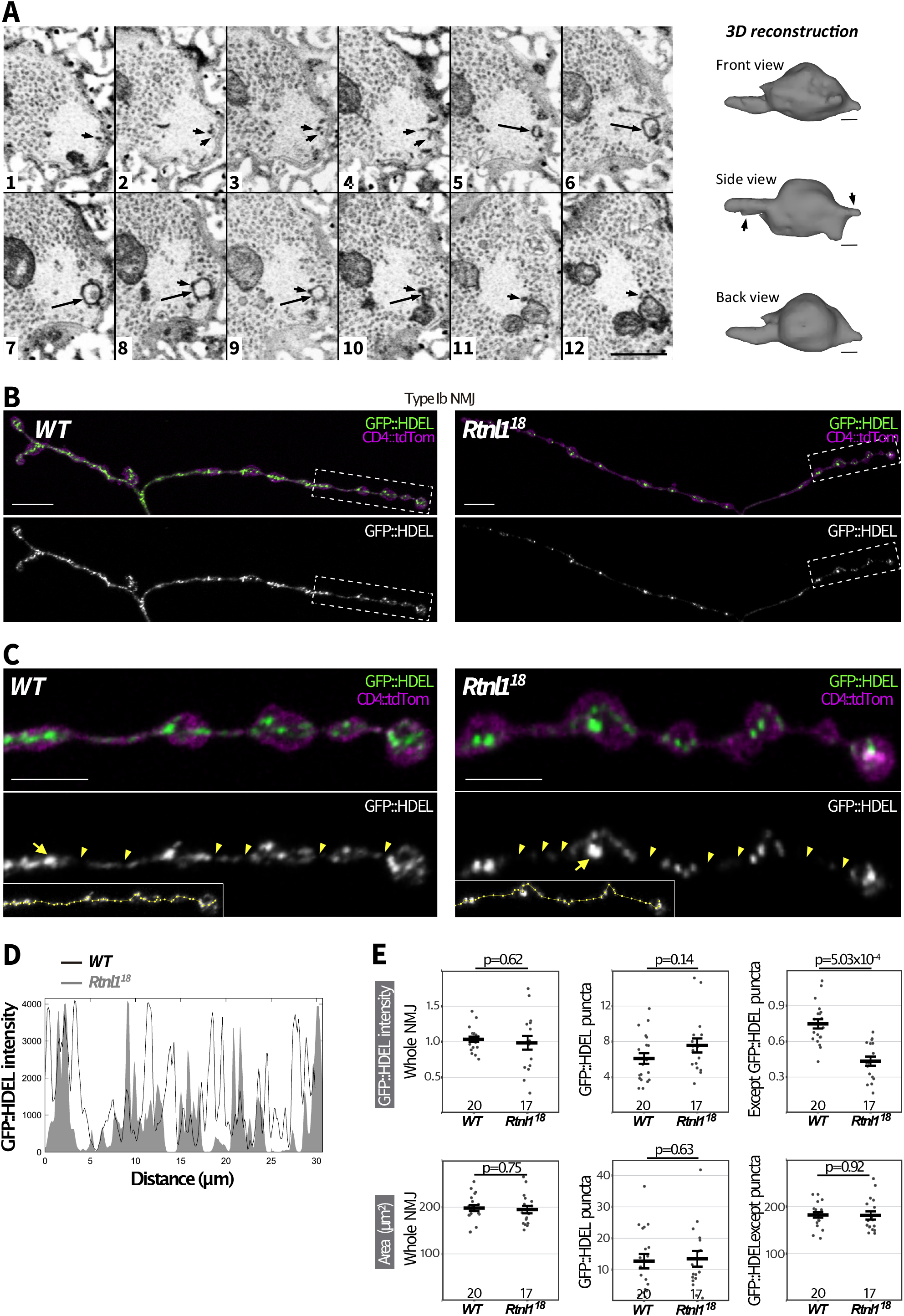
Rtnl1 loss causes localized depletion of presynaptic ER lumenal marker. (**A**) A series of twelve serial ATUMtome 50-nm sections through a ROTO-stained larval bouton, visualized by scanning EM and inverted grayscale images. An example is shown of a cisterna (large arrows in sections 5-10) that appears to be continuous with tubules in the same and neighboring sections (smaller arrows), in a region with few synaptic vesicles. Tubules are identified as darkly staining puncta in successive sections; their lumen is too small to detect using this method (Terasaki, 2018; Yalçın et al., 2017). Scale bar in panel 12, 500 nm. The cisterna highlighted by the large arrow and its attached tubules (arrows in “Side View”) were used to generate a reconstruction, shown on the right and viewed from three different angles. EM section 1 is on the left of the reconstruction, section 12 on the right. Scale bars, 70 nm. Larval genotype is *Rtnl1*^*18*^ */ WT*. (**B**) Representative confocal projections showing GFP::HDEL in Type Ib muscle 1 NMJs of *WT* and *Rtnl1*^*18*^ larvae. Scale bars, 10 μm. (**C**) Magnified views of the areas inside broken lines in **B**. Scale bars, 5 μm. Arrowheads indicate regions with low levels of GFP::HDEL which we interpret as tubules, and arrows indicate examples of puncta which we interpret as cisternae. Insets show the lines drawn along GFP::HDEL signal to plot intensity profiles (**D**). (**E**) GFP::HDEL intensity across the whole NMJ, within a region of interest (ROI) containing all puncta, and in the NMJ excluding the puncta ROI. Plots show individual larval datapoints and mean ± SEM; intensity levels indicate arbitrary units (au) after normalization to control (*WT*); intensity values are relative to CD4::tdTom (Han et al., 2011) signal. Larval datapoints were calculated and are shown and compared using Student’s t-tests as in **Fig. 2**. Genotypes are *Ib-GAL4, UAS-CD4::tdTom / UAS-BiP::sfGFP::HDEL*, in either a *Rtnl1*^*+*^ (*WT*) *or Rtnl1*^*18*^ background. Transheterozygous *Rtnl1*^*18*^ */ Rtnl1*^*1*^ mutant larvae also showed a similar phenotype to *Rtnl1*^*18*^ (**Supp. Fig. S5B**).

In *Rtnl1*^*18*^ NMJs, labeling of bright GFP::HDEL puncta was not affected, but labeling between puncta was significantly decreased compared to *WT* (Fig. 3B-E; Supplementary Fig. S5B). We conclude that the tubular ER network, but not the amount or volume of ER cisternae, is reduced in *Rtnl1* mutant NMJs; and that the stronger depletion of ER membrane markers in *Rtnl1* mutant genotypes reflects the higher proportion of these markers in tubules compared to cisternae, due to the high surface area/volume ratio of presynaptic ER tubules.

### Rtnl1 loss reduces STIM-mediated ER-PM contact sites

The decreased levels of ER membrane and tubules, but not cisternae, in *Rtnl1* mutant presynaptic termini provided a unique model to explore the specific functional and physiological roles of presynaptic ER tubules. Since contact sites must depend on availability of ER membrane, we first tested two critical roles of ER-PM contact sites, in lipid and Ca^2+^ exchange between both compartments (Öztürk et al., 2020).

The PM phospholipid phosphatidylinositol 4,5-bisphosphate (PI(4,5)P2) is derived from phosphatidylinositol (PI) formed at the ER membrane. PI(4,5)P2 binds to ER membrane proteins to mediate ER-PM tethering (Muallem et al., 2017), and it is also required for presynaptic function (Lauwers et al., 2016). However, using the sensor PLCL::PH::GFP (Verstreken et al., 2009), we did not detect any difference in PI(4,5)P2 levels between *WT* and *Rtnl1* NMJs (Supplementary Fig. S6). Although we cannot exclude that other ER-PM contact-site-dependent lipids are affected in *Rtnl1* mutants, a decrease in presynaptic tubular ER does not necessarily impact lipid homeostasis.

ER-PM contact sites can also mediate Ca^2+^ exchange. STIM proteins are ER Ca^2+^ sensors that can also bind to the PM at ER-PM contact sites, promoting SOCE upon emptying of ER Ca^2+^ stores (Serwach & Gruszczynska-Biegala, 2020). STIM proteins are distributed throughout the ER membrane, but enriched at ER-PM contacts, which increase in number when ER is depleted of Ca^2+^ (Liou et al., 2005; Shen et al., 2021). *Drosophila* has one STIM gene, orthologous to mammalian STIM1 and STIM2 (Williams et al., 2001). In *WT* motor neuron terminals, STIM::mCherry (Bi et al., 2014) showed, as expected, a mostly continuous distribution, with some foci characteristic of STIM accumulation at ER-PM contact sites. STIM:mCherry levels were generally reduced in *Rtnl1* NMJs (Fig. 4), like other ER membrane markers (Fig. 2), consistent with less presynaptic ER surface. STIM::mCherry foci were slightly less intense and took up less area in *Rtnl1* mutants than in *WT* larvae, suggesting a reduction of these ER-PM contact sites (Fig. 4), and that Ca^2+^ stores are not depleted in *Rtnl1* mutants. STIM::mCherry foci area was highly variable among wildtype NMJs, but consistently low in *Rtnl1* NMJs, suggesting that *WT* presynaptic ER could be responding to physiological demands (e.g. variable emptying of ER Ca^2+^) by adjusting the number of STIM-containing ER-PM contact sites.

**Figure 4.**
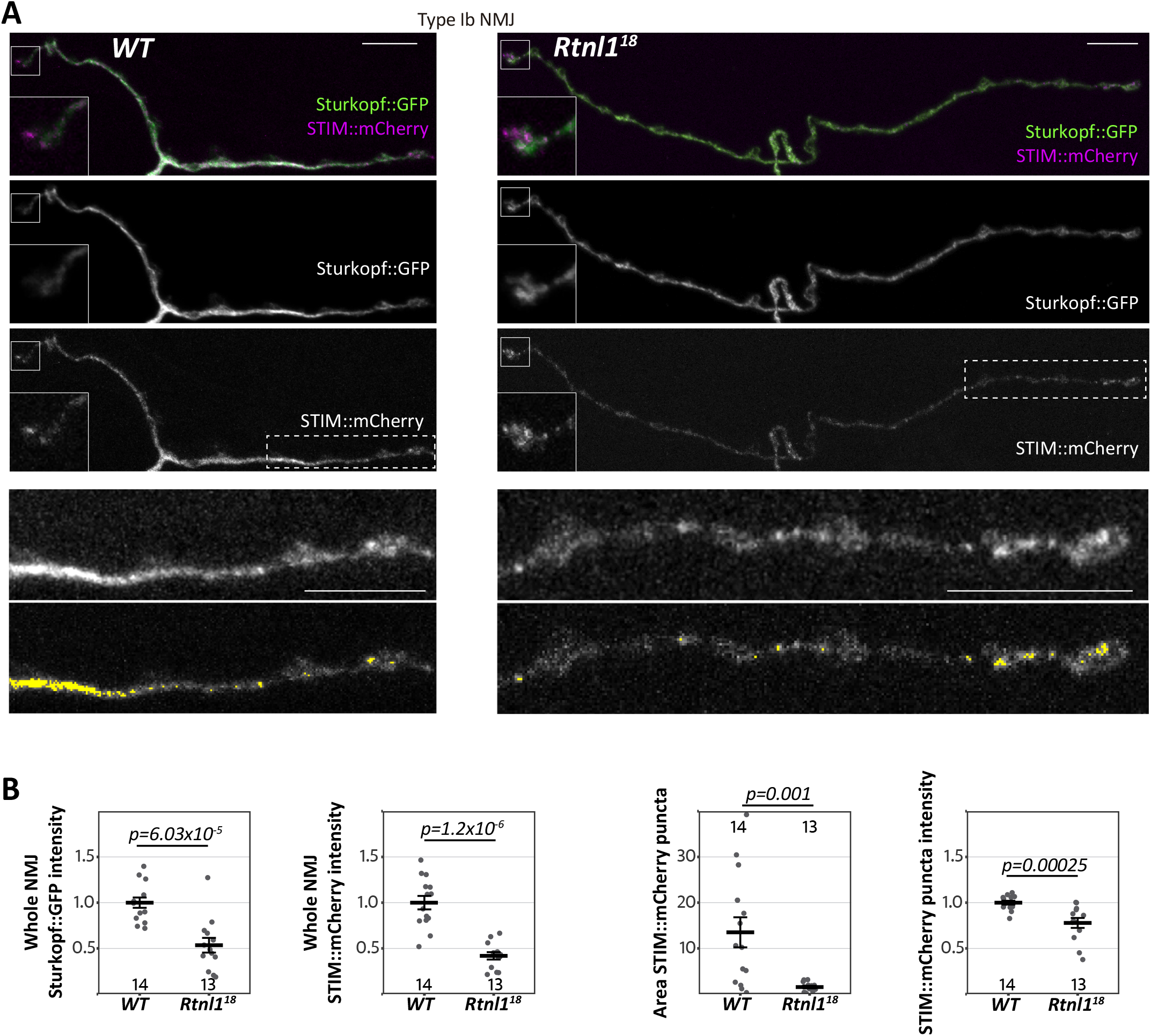
Rtnl1 loss depletes presynaptic STIM and reduces STIM foci. (**A**) Representative confocal projections showing the ER markers STIM::mCherry and Sturkopf::GFP in Type Ib muscle 1 NMJs of *WT* and *Rtnl1*^*18*^ larvae. Insets in the top three rows show a magnified view of terminal boutons. Bottom panels show magnified views of the areas indicated with a broken line. Areas of elevated STIM (“foci”) are sometimes punctate and sometimes extended, and are highlighted in yellow in the bottom panels. Scale bars, 10 μm. Insets, 6×6 μm. (**B**) Plots show individual larval datapoints and mean ± SEM; intensity levels indicate arbitrary units (au) after normalization to control (*WT*). Larval datapoints were calculated and are shown as in **Fig. 2**; Except for Sturkopf::GFP intensity (Student’s t-test), pairwise comparisons were performed with Mann-Whitney U tests. Genotypes are *Ib-GAL4, UAS-Sturkopf::GFP / UAS-STIM::mCherry*, in either a *WT or Rtnl1*^*18*^ background.

### *Rtnl1* loss reduces neurotransmitter release and evoked presynaptic Ca^2+^ influx by impairing STIM-dependent ER-PM contacts

Synaptic strength is regulated by Ca^2+^, both extracellular and in the ER. Alterations to ER as the largest intracellular Ca^2+^ store could therefore potentially alter presynaptic Ca^2+^ physiology, with consequences for processes such as synaptic strength and ATP generation, that could contribute to neurodegeneration in conditions including the HSPs. However, since the levels of the lumenal marker GFP::HDEL (Fig. 3) and the lumenal Ca^2+^ sensor GCaMP-210 (Oliva et al., 2020) are relatively unaffected by loss of Rtnl1, processes that depend simply on the amount or capacity of presynaptic ER Ca^2+^ storage might not necessarily be affected. We therefore tested how far synaptic function and presynaptic Ca^2+^ fluxes were affected by the loss of Rtnl1, which reduced presynaptic ER membrane and ER tubules, but left ER volume and Ca^2+^ storage capacity largely intact.

We first tested a readout of synaptic transmission using a Ca^2+^ indicator inserted in the postsynaptic muscle membrane, Mhc-SynapGCaMP6f (Newman et al., 2017). Rtnl1 loss caused a large decrease in postsynaptic Ca^2+^ responses to low-frequency stimulation, suggesting decreased presynaptic neurotransmitter release (Supplementary Fig. S7A-C; Videos 1, 2). This result is consistent with strongly reduced EJP amplitudes in *Rtnl1*^*1*^ NMJs (Summerville et al., 2016). Loss of Rtnl1 also strongly reduced the frequency of miniature transmission (spontaneous vesicle release, seen as transient highly localized elevations in SynapGCaMP6f fluorescence) (Supplementary Fig. S7D-E; Supplementary Fig S8; Videos 3, 4). Loss of Rtnl1 had no effect on the amplitude of miniature transmission (Supplementary Fig. S7F) indicating that loss of Rtnl1 does not change neurotransmitter load per vesicle.

We next tested whether reduced synaptic transmission at *Rtnl1* mutant NMJs could be due to low presynaptic Ca^2+^ responses to neuronal activity. We tested Type Ib and Is presynaptic terminals separately, due to their different physiological properties that could potentially be linked to Ca^2+^ storage and fluxes, and their different contributions to postsynaptic responses (Aponte-Santiago et al., 2020; Aponte-Santiago & Littleton, 2020; Chouhan et al., 2010). A Ca^2+^ sensor targeted to the cytosolic face of the presynaptic plasma membrane, myrGCaMP6s, showed no significant change in resting fluorescence in both Type Is and Type Ib *Rtnl1*^*18*^ boutons compared to *WT* (Fig. 5A; Supplementary Fig. S9A), indicating that Rtnl1 loss does not change resting cytosolic [Ca^2+^]. We corroborated this conclusion using a ratiometric cytosolic Ca^2+^ sensor *UAS-tdTom-p2a-GCaMP56* (Daniels et al., 2014), which also detected no difference in resting cytosolic Ca^2+^ in Type Ib *Rtnl1*^*18*^ boutons compared to *WT* (Fig. 5B). Evoked myrGCaMP6s responses (Videos 5-6) were reduced in Type Ib and Is *Rtnl1*^*18*^ boutons at all stimulation frequencies tested (Fig. 5C-D; Supplementary Fig. S9B, S10A). This reduction appeared to be due to loss of Rtnl1; it could be rescued by a *UAS-Rtnl1::HA* transgene (Fig. 5E); furthermore, transheterozygous *Rtnl1*^*1*^/*Rtnl1*^*18*^ mutant larvae displayed the same phenotype as homozygous *Rtnl1*^*18*^ mutants (Supplementary Fig. S9B-C). Interestingly, synaptic size was not reduced in *Rtnl1*^*18*^ NMJs; instead they showed a small but significant increase in the number of synaptic boutons (muscle 1 Type Ib NMJ; *WT:* 24.6 ± 0.5 (mean ± SEM), n=18 larvae; *Rtnl1*^*18*^: 27.8 ± 1.0 (mean ± SEM), n=17 larvae; p=0.0075**, Student’s T-test), consistent with previous data from *Rtnl1*^*1*^ (Summerville et al., 2016). Loss of Rtnl1 did not affect the dynamics of the response, as neither the time to peak response, nor the recovery half-time of Type Is or Ib boutons were affected (Supplementary Fig. S9D-F, S10B-D). Taken together, our data suggest reduced neurotransmission in *Rtnl1*^*18*^ NMJs. This reduced neurotransmission could potentially be accounted for by reduced evoked presynaptic cytosolic Ca^2+^ responses.

**Fig 5.**
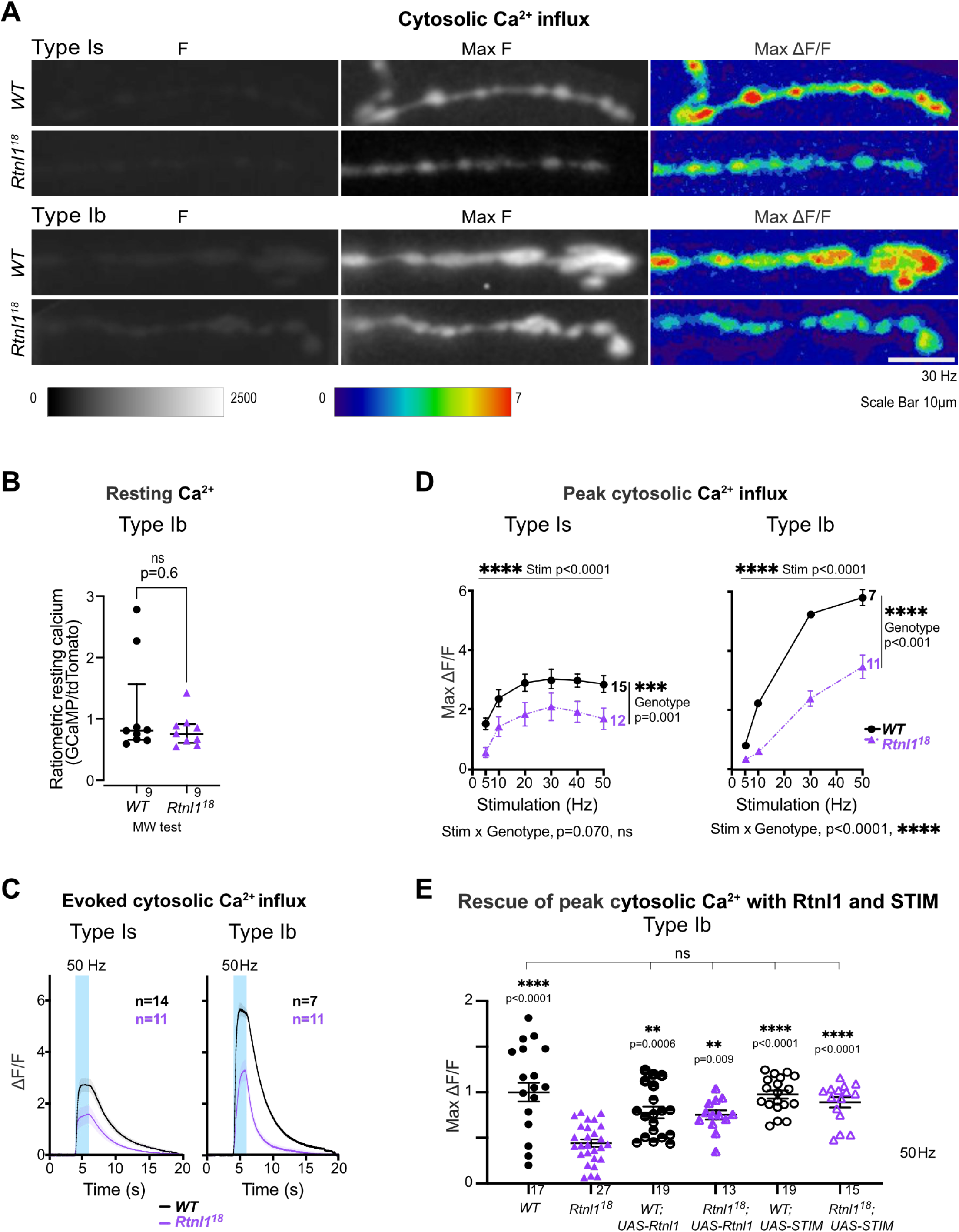
Loss of Rtnl1 decreases evoked cytosolic Ca^2+^ responses in presynaptic NMJs. Cytosolic Ca^2+^ responses to 2 s of 30-Hz stimulation were measured at Type Is and Type Ib termini at muscle 1 in segment A4-A6, using myr::GCaMP. (**A**) Panels show GCaMP fluorescence at rest (F), maximum fluorescence (Max F), and maximum relative change in fluorescence (Max ΔF/F) in representative examples of *WT* and *Rtnl1*^*18*^ presynaptic terminals. (**B**) The ratiometric cytosolic Ca^2+^ sensor tdTom-p2a-GCaMP56 detected no difference in resting cytosolic Ca^2+^ in Type Ib *Rtnl1*^*18*^ boutons compared to *WT*. Graphs show individual larval datapoints for the ratio of GCaMP to Tomato fluorescence, with one Type Ib muscle 1 NMJ per larva, averaged across the responding area. Genotypes were compared using a Mann-Whitney U test, with median ± interquartile range shown. (**C**) Timecourses of evoked cytosolic GCaMP responses to 50 Hz stimulation (mean ± SEM of larval datapoints). (**D**) Loss of Rtnl1 significantly decreases cytosolic Ca^2+^ responses across a range of stimulation frequencies. Graph shows mean ± SEM of maximum evoked ΔF/F values (larval datapoints), with comparisons using mixed-effects model repeated-measures ANOVA. (**A**-**D**) Genotypes: *Is-GAL4* or *Ib*-*GAL4, UAS-myr::GCaMP6s / UAS-tdTom::Sec61*β in either a *WT* or *Rtnl1*^*18*^ mutant background. (**E**) Expression of *UAS-Rtnl1::HA* or *UAS-STIM::mCherry* using *Ib-GAL4* rescues evoked cytosolic Ca^2+^ response in mutant larvae. Data were normalized to the *WT* mean of maximum cytosolic GCaMP responses (ΔF/F) evoked by 50 Hz stimulation, from two different microscope setups. Graph shows individual larval datapoints, with one NMJ per larva, averaged across the responding area. Genotypes: *Ib*-*GAL4, UAS-myr::GCaMP6s / attP2*^*+*^, or *Ib*-*GAL4, UAS-myr::GCaMP6s / UAS-Rtnl1::HA*, or *Ib*-*GAL4, UAS-myr::GCaMP6s / UAS-STIM::mCherry*, in either a *WT* or *Rtnl1*^*18*^ mutant background; the *attP2* landing site acts as a *W*T control for insertion of *UAS-Rtnl1::HA* at *attP2*.

Since STIM-mediated ER-PM contact sites are known to mediate Ca^2+^ exchange, and both STIM levels and foci were reduced in *Rtnl1* mutants (Fig. 4), we tested whether STIM levels were associated with the observed defects in presynaptic Ca^2+^ influx. We found that overexpression of STIM::mCherry in motor neurons rescued cytosolic Ca^2+^ influx in Rtnl1 mutants (Fig. 5E), suggesting that depletion of the presynaptic ER surface may lower presynaptic Ca^2+^ influx by downregulating STIM signaling.

### Rtnl1 loss decreases evoked ER Ca^2+^ uptake at Type Is and Type Ib NMJs

Further, we examined whether depletion of ER tubules differentially affected ER Ca^2+^ fluxes in NMJs with phasic or sustained firing profiles, Type Is and Type Ib respectively.

At *WT* Type Is terminals, an ER lumenal GCaMP6-210 reporter showed rapid transient decreases in ER lumenal fluorescence immediately after stimulation, suggesting release of Ca^2+^ to the cytosol evoked by repetitive stimulation, followed by a slow increase in fluorescence, which was more pronounced at higher stimulation frequencies (Fig. 6; Videos 7-8), and lagged and outlasted the evoked elevation of cytosolic Ca^2+^ (Fig. 5; Supplementary Figs. S9, S10). Rtnl1 loss did not significantly change resting fluorescence (Fig. 6A-B), nor the rapid release (Fig. 6C) across most stimulation frequencies, except that release was slightly higher at the lowest frequency, 5 Hz (Fig. 6D). However, *Rtnl1*^*18*^ mutants showed less Ca^2+^ uptake by ER than *WT*, after the initial rapid release (Fig. 6C) across all stimulation frequencies tested (Fig. 6E; Supplementary Fig. S11A), suggesting that the ER tubule depletion in *Rtnl1* mutants primarily diminishes the uptake of ER Ca^2+^ in Type Is boutons.

**Fig 6.**
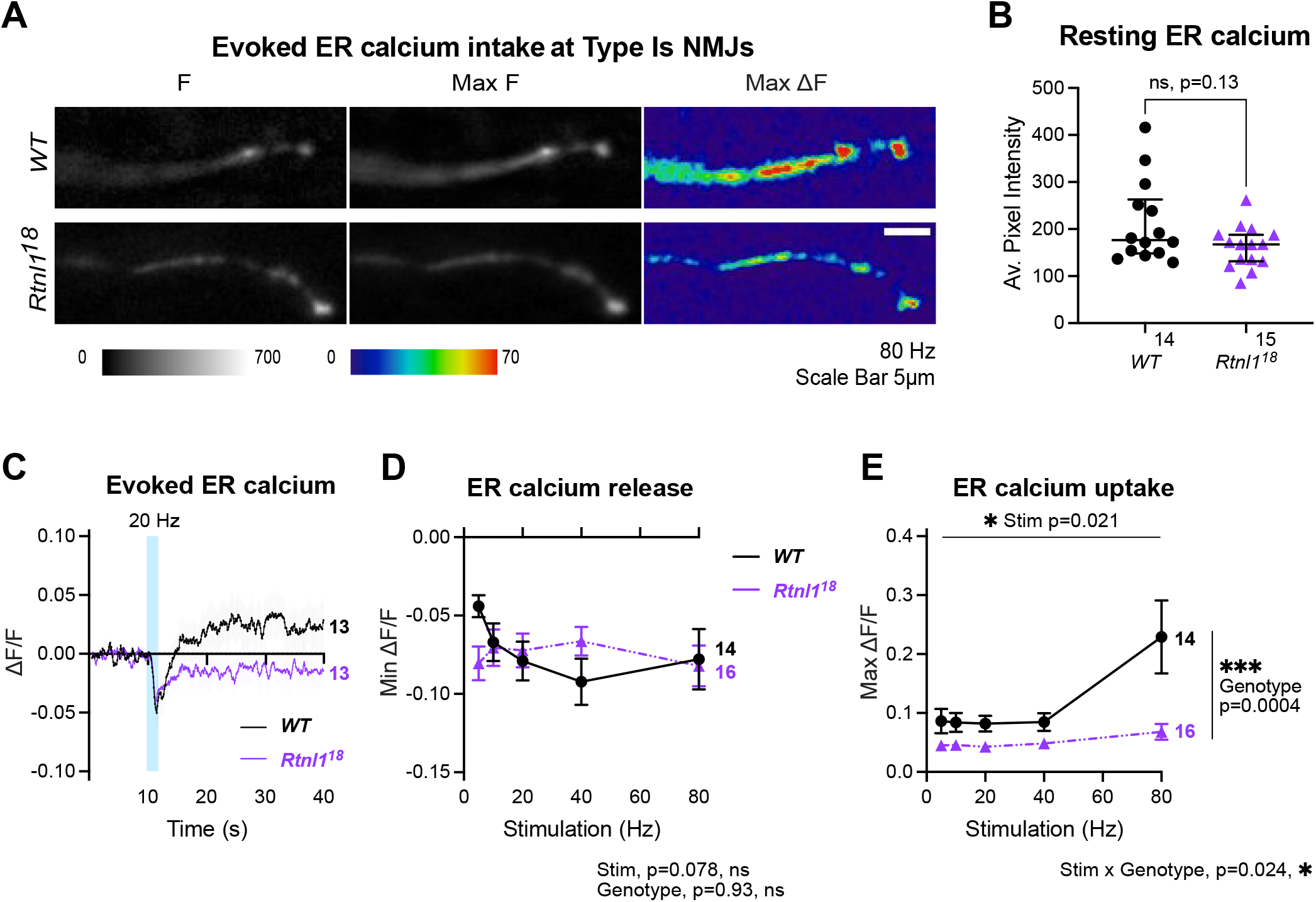
Rtnl1 loss decreases evoked ER Ca^2+^ uptake at Type Is termini. Evoked ER Ca^2+^ responses were measured at Type Is termini at muscle 1 in segment A4-A6. (**A**) Lumenal GCaMP fluorescence at Type Is NMJs, presented at rest (F), maximum fluorescence (Max F), and maximum change in fluorescence (Max ΔF) in representative examples of *WT* and *Rtnl1*^*18*^ presynaptic terminals. (**B**) Loss of Rtnl1 does not affect resting ER Ca^2+^. The graph shows larval datapoints as in **Fig. 5B**, with median ± interquartile ranges, compared using a Mann-Whitney U-test. (**C**) ΔF/F timecourse of evoked lumenal GCaMP responses to 20 Hz stimulation (mean ± SEM). (**D**) Loss of Rtnl1 does not affect ER Ca^2+^ release immediately following stimulation. The graph shows the minimum ΔF/F value after stimulation. (**E**) Loss of Rtnl1 significantly decreases evoked ER Ca^2+^ uptake over a range of stimulation frequencies. The graph shows the maximum ΔF/F value reached after stimulation. In **D**-**E**, graphs and analyses are as for **Fig. 5D**. Genotypes: *Is-GAL4, UAS-ER-GCaMP6-210 / UAS-tdTom::Sec61*β in either a *WT* or *Rtnl1*^*18*^ mutant background.

*WT* Type Ib terminals only occasionally showed the initial rapid evoked decrease in ER lumenal fluorescence seen in Type Is termini, and showed a strong and sustained increase in lumenal fluorescence that lagged and outlasted the cytosolic Ca^2+^ response, similar to Type Is boutons, but stronger (Fig. 7; Videos 9-11). Rtnl1 loss again did not alter resting ER lumenal GCaMP fluorescence relative to *WT* (Fig. 7A-B), but often caused the initial rapid Ca^2+^ release to appear (Fig. 7C-D) at every stimulation frequency tested (Fig. 7E; Supplementary Fig. S11B), and led to lower evoked ER Ca^2+^ uptake (Fig. 7F), similar to Type Is termini.

**Fig 7.**
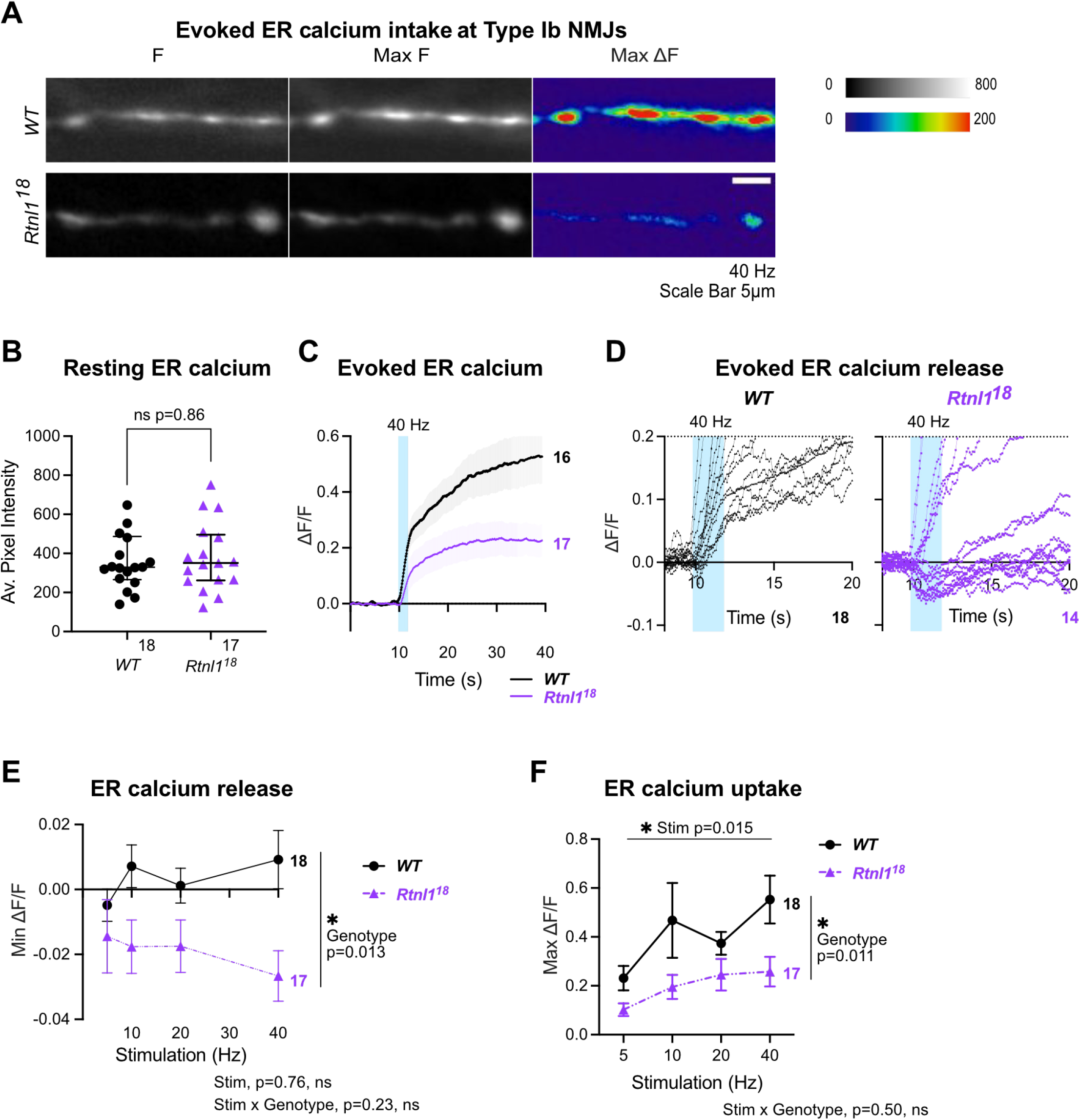
Rtnl1 loss decreases evoked ER Ca^2+^ uptake in Type Ib termini. Evoked ER Ca^2+^ responses were measured at Type Ib termini at muscle 1 in segment A4-A6. (**A**) Lumenal GCaMP fluorescence at Type Ib NMJs presented as in **Fig. 6A**. (**B**) Loss of Rtnl1 does not affect resting ER resting lumen GCaMP. Graphing and analysis are as for **Fig. 6B**. Some outlier datapoints are excluded from the graph, but included in statistics. (**C**) ΔF/F time course of evoked lumenal GCaMP responses to 40 Hz stimulation (mean ± SEM). (**D**) ΔF/F time courses from individual larvae show that transient evoked Ca^2+^ release from ER, common in Type Is termini (**Fig. 6C**) is mostly undetectable in *WT* Type Ib termini, but found in over half of *Rtnl1*^*18*^ Type Ib termini tested. (**E**) Loss of Rtnl1 increases ER Ca^2+^ release across a range of stimulation frequencies. (**F**) Loss of Rtnl1 significantly decreases evoked ER Ca^2+^ uptake over a range of stimulation frequencies. Graphs in **E** and **F** are presented and analyzed as in **Fig. 5D**. Genotypes: *Ib*-*GAL4, UAS-ER-GCaMP6-210 / UAS-tdTom::Sec61*β in either a *WT* or *Rtnl1*^*18*^ mutant background.

### *Rtnl1* loss decreases mitochondrial Ca^2+^ uptake at Type Is and Type Ib NMJs

Mitochondrial Ca^2+^ stimulates ATP production (Datta & Jaiswal, 2021), and since Ca^2+^ released from ER and/or present in the cytosol can potentially be taken up by mitochondria, we hypothesized that mitochondrial Ca^2+^ could be impaired by the changes in ER architecture elicited by Rtnl1 loss. We tested this hypothesis using a Ca^2+^ sensor adapted for mitochondrial conditions, CEPIA3mt (Suzuki et al., 2014).

In *WT* Type Is and Ib NMJs, mitochondria showed a rapid increase in [Ca^2+^] in response to repetitive stimulation, followed by a rapid partial decline, and a slow return to baseline over a period greater than 40 seconds (usually 1-2 minutes) (Fig. 8; Videos 12-13). As with ER lumenal Ca^2+^, this response followed and outlasted the evoked cytosolic Ca^2+^ response. Loss of Rtnl1 did not affect resting CEPIA3mt fluorescence in either Type Is or Ib boutons (Fig. 8A-B, Supplementary Figs. S12A, S13A-B). Immunostaining for the myc epitope tag of the CEPIA3mt sensor showed that CEPIA3mt sensor levels were not altered between *WT* and *Rtnl1*^*18*^ larvae (Supplementary Fig. S13A, C), implying that the levels of expression of the sensor were comparable and the CEPIA3mt fluorescence is reporting relative [Ca^2+^] in these genotypes. Although *Rtnl1*^*1*^ NMJ mitochondria have been reported as larger than *WT* (O’Sullivan et al., 2012), we detected only a small and non-significant increase in mitochondrial size in *Rtnl1*^*18*^ muscle 1 NMJs compared to *WT* (Supplementary Fig. S13D).

**Fig 8.**
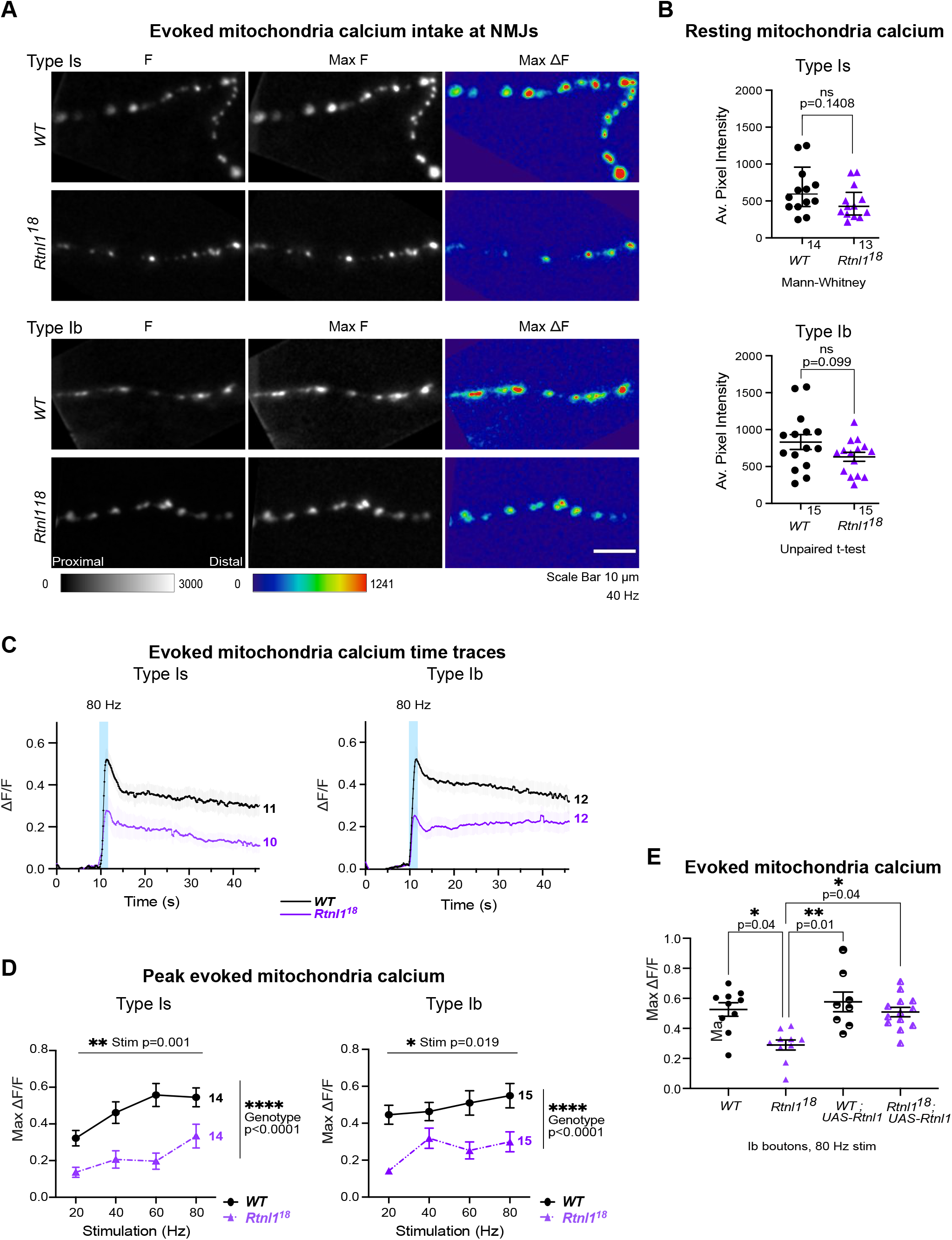
*Rtnl1* loss decreases evoked mitochondrial Ca^2+^ uptake in Ib and Is presynaptic termini. Evoked mitochondrial Ca^2+^ responses were measured at Type Ib and Type Is termini at muscle 1 in segment A4-A6. (**A**) Mitochondrial CEPIA3mt fluorescence presented as in **Fig. 6A**. (**B**) Loss of Rtnl1 does not alter resting mitochondrial Ca^2+^ levels in Type Is or Type Ib terminals. Some outlier datapoints are excluded from the graph, but included in statistics. Graphs and analyses are as in **Fig. 6B**, except that Type Ib are shown as mean ± SEM, and compared using Student’s t-test. (**C**) Time courses of evoked mitochondrial CEPIA3mt responses to 80 Hz stimulation (mean ± SEM) show reduced Ca^2+^ uptake by mitochondria on loss of Rtnl1. (**D**) Loss of Rtnl1 significantly decreases mitochondrial Ca^2+^ uptake across a range of stimulation frequencies. Graphs and analyses are as for **Fig. 5D**. (**A**-**D**) Genotypes: *Is-GAL4* or *Ib*-*GAL4, UAS-CEPIA3mt / UAS-tdTom::Sec61*β in either a *WT* or *Rtnl1*^*18*^ mutant background. (**E**) Expression of *UAS-Rtnl1::HA* under *Ib-GAL4* control rescues evoked Ca^2+^ uptake by mitochondria in mutant larvae. Data shows maximum evoked mitochondrial CEPIA3mt responses (ΔF/F) after 80 Hz stimulation. Graph shows individual larval datapoints, with one NMJ datapoint per larvae, averaged across the responding area. Genotypes: *Ib*-*GAL4, UAS-CEPIA3mt / UAS-Rtnl1::HA* or *Ib*-*GAL4, UAS-CEPIA3mt / attP2*^*+*^ in either a *WT* or *Rtnl1*^*18*^ mutant background; the *attP2* landing site acts as a *WT* control for insertion of *UAS-Rtnl1::HA* at *attP2*.

However, Rtnl1 loss significantly lowered evoked NMJ mitochondrial Ca^2+^ uptake compared to *WT* at every stimulation frequency tested (Fig. 8C-D; Supplementary Figs. S12B, S13E). Transheterozygous mutant controls had statistically similar responses to homozygous *Rtnl1*^*18*^ larvae (Supplementary Fig. S12B-C), corroborating that the phenotype is due to loss of Rtnl1. Further, expression of *UAS-Rtnl1::HA* under GAL4 control fully rescued evoked mitochondria Ca^2+^ uptake in Ib presynaptic boutons (Fig. 8E).

Although Type Is and Ib NMJs have different mitochondrial densities, responses to nerve stimulation, and Ca^2+^ sequestration capacities (Chouhan et al., 2010), Rtnl1 loss reduced evoked mitochondria Ca^2+^ responses in both bouton classes. Heterozygous *Rtnl1*^*18*^*/WT* NMJ mitochondria Ca^2+^ responses were similar to *WT* larvae at the higher stimulation frequencies, but were reduced at lower frequencies, suggesting a mild dominant effect of *Rtnl1*^*18*^ (Supplementary Fig. S12B-C).

Genotype did not affect time to peak evoked Ca^2+^ in either Is or Ib boutons (Supplementary Figs. S12D, S13F), nor time to 50% recovery in Ib boutons (Supplementary Fig. S13G). In mitochondria of Type Is boutons, loss of Rtnl1 elicited faster 50% recovery times, in both *Rtnl1*^*18*^ mutants and transheterozygous controls, compared to *WT* controls (Supplementary Fig. S12E).

### Comparison of Ca^2+^ responses and their Rtnl1 dependence across compartments

Comparison of the above Ca^2+^ responses in different organelles (Fig. 9) revealed a rapid rise and fall in cytosolic Ca^2+^, followed by strong and longer lasting Ca^2+^ responses in both mitochondria and ER. These time courses reveal that loss of Rtnl1 affects the evoked Ca^2+^ responses of the three presynaptic compartments with similar severity, with Ca^2+^ responses being reduced to just over a half of those of *WT* in Ib boutons and Is cytosolic responses, and just under half in Is mitochondria and ER responses.

**Fig 9.**
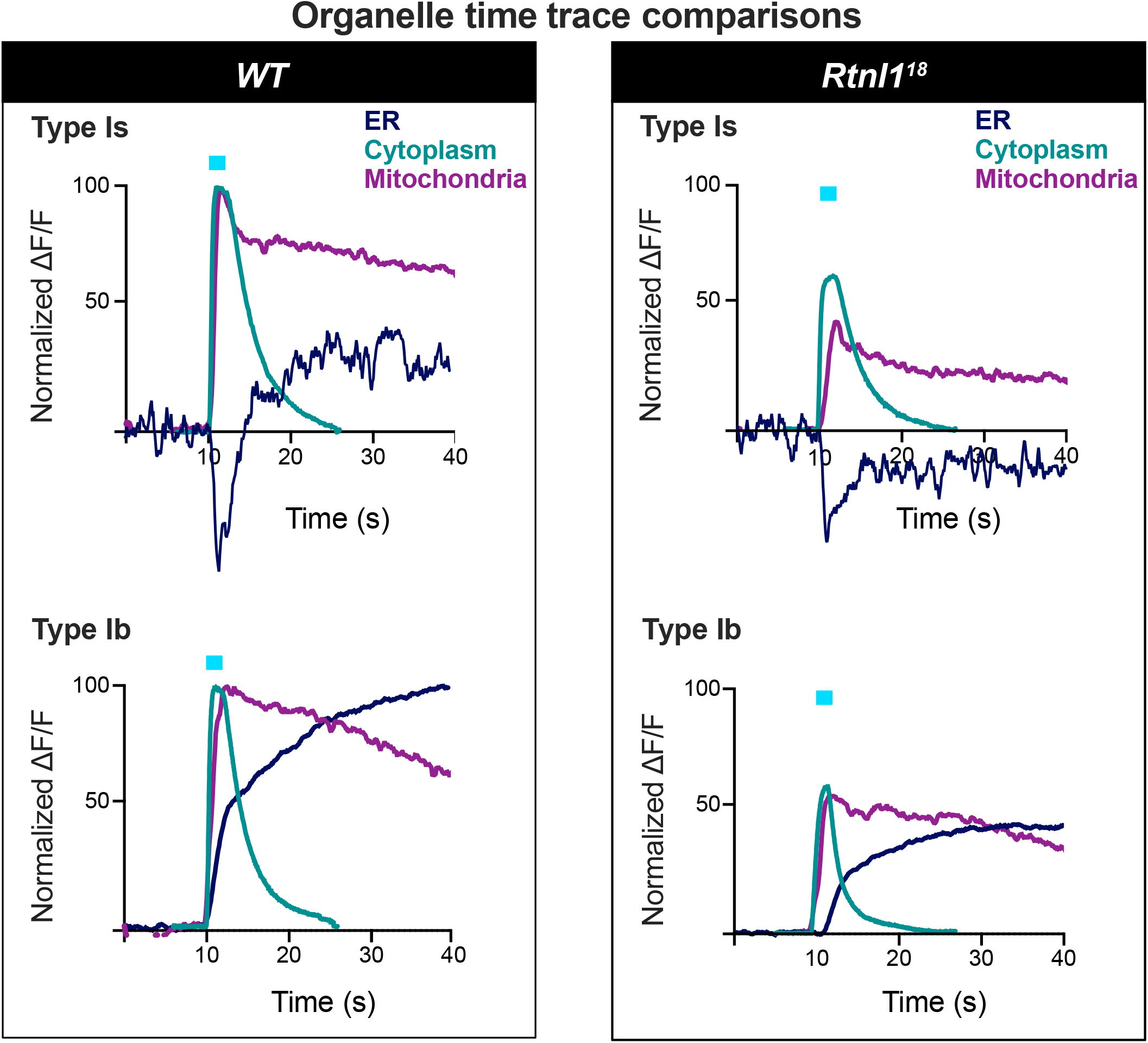
A comparison of superimposed time courses. Mean time traces (data from **Fig. 5** - **Fig. 8**) from ER, mitochondria, and cytosolic sensors in *WT* and a *Rtnl1* mutant (*Rtnl1*^*-*^), and in Type Is and Ib excitatory synapses. Plots are normalized so that the difference of minimum to maximum ΔF/F values are set to 100 for the *WT* recordings in each compartment. Most time courses are from 40 Hz stimulation trials. Two exceptions are: the Ib cytosolic trace was with 50 Hz stimulation, as 40 Hz was not tested, Is ER responses were with 20 Hz, as this frequency was most representative of the lack of statistical difference between *WT* and *Rtnl1* mutant responses **(Fig. 6D)**.

## DISCUSSION

From a body of work mostly in cultured non-neuronal cells, we know that ER controls Ca^2+^ handling (Öztürk et al., 2020). This role appears conserved in presynaptic ER, where both ER-PM contacts (de Juan-Sanz et al., 2017) and the amount of ER (Kuijpers et al., 2021) are critical for neurotransmission. Given the relatively larger Ca^2+^ storage capacity of ER cisternae, are presynaptic ER tubules contributing to Ca^2+^ dynamics? In this work, we have disrupted the tubular ER-shaping protein Rtnl1, and found a decrease of presynaptic ER tubules (schematized in Fig. 10). Rtnl1 loss results in a unique model to study the role of ER tubules at the presynaptic region. Here, we use this model to analyze Ca^2+^ handling, which relates with neurotransmission and the biological basis of neurological disorders associated with tubular ER.

**Fig 10.**
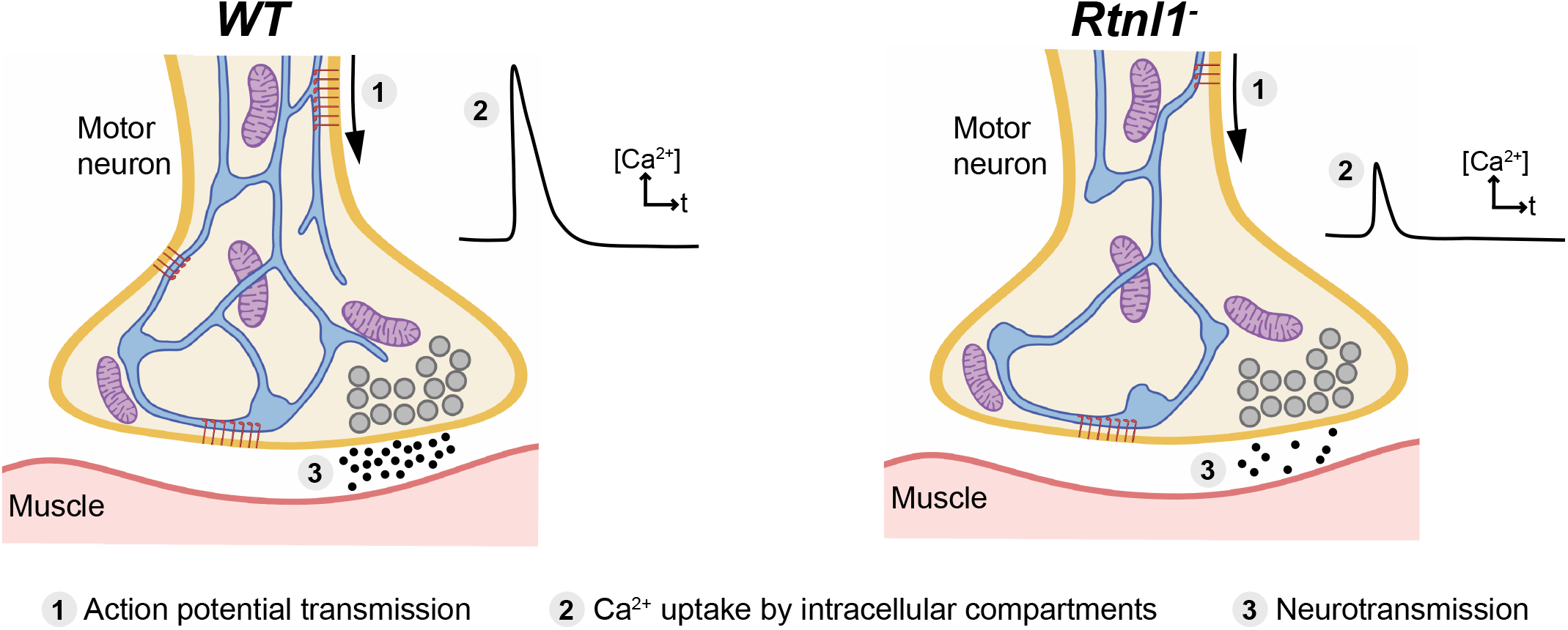
Model of effect of Rtnl1 loss on presynaptic ER network and synaptic function. Schematic diagrams of a *WT* and a *Rtnl1* mutant (*Rtnl1*^*-*^) presynaptic terminal. The latter shows fewer ER tubules, resulting in less ER network surface, and hence in less ER contact surface with other cell compartments. Reduced presynaptic ER surface is accompanied by generally lower Ca^2+^ fluxes in presynaptic compartments (cytosol, ER lumen and mitochondria), and by lower neurotransmission. These reductions could be a potential route by which the loss of ER-shaping HSP protein function might lead to motor neuron axonopathy. Key: ER network, blue; mitochondria, purple; synaptic vesicles, grey; MN PM, dark orange; muscle PM, dark red; ER-PM contacts, accumulation of STIM1 (red rods); released neurotransmitter, black dots.

Loss of either Rtnl1 or REEPs causes a loss of axonal ER preferentially in longer motor neurons of *Drosophila* larvae (Yalçın et al., 2017). In contrast, the decrease of presynaptic ER network in *Rtnl1* mutants is independent of axon length (Fig. 2A-B). The tubular ER network in presynaptic terminals (mouse nucleus accumbens) is more extensive than in axons (Wu et al., 2017). It is therefore possible that presynaptic ER is more dependent on the function of tubular ER-shaping proteins than axonal ER. Regarding the continuity of the network, it is possible that neurons can better tolerate loss of ER tubules in presynaptic terminals than in axons, where there are relatively few tubules to begin with. This might explain why loss of Rtnl1 causes ER network discontinuity in axons (Yalçın et al., 2017), but not at the presynaptic region.

Increasing the levels of axonal and presynaptic ER (by blocking neuronal autophagy) correlates with elevated Ca^2+^ release from ER, which in turn increases neurotransmission (Kuijpers et al., 2021). In agreement with this, we show that lower ER levels correlate with decreased neurotransmission (Supplementary Fig. S7), indicating that the relationship works conversely. Interestingly, we also found a severe decrease in the frequency of miniature neurotransmission in *Rtnl1* mutants (Supplementary Fig. S7D-F; Supplementary Fig. S8). Since presynaptic ER tubules, but not cisternae, appear to be reduced in *Rtnl1* mutants (Fig. 3), we propose that tubules help regulate Ca^2+^ handling across presynaptic compartments. Tubule loss abates Ca^2+^ handling in the ER (Fig. 6, 7), cytosol (Fig. 5), and mitochondria (Fig. 8), in both types of excitatory NMJs in *Drosophila* larvae. As neurotransmitter release is triggered by cytosolic Ca^2+^ influx through PM voltage-gated Ca^2+^ channels (Olivera et al., 1994; Dunlap et al, 1995; Catterall et al., 2013), the decrease of postsynaptic Ca^2+^ responses (Supplementary Fig. S7) and presynaptic cytosolic influx (Fig. 5) corroborate each other.

ER-PM contacts have an inhibitory role in cytosolic Ca^2+^ influx in primary hippocampal neurons via activated STIM1 on emptying of ER stores (de Juan-Sanz et al., 2017); other studies in neurons have demonstrated that emptying of Ca^2+^ from ER stores (thereby activating STIM) promotes spontaneous (miniature), but not evoked vesicle release at excitatory terminals by the action of STIM2 (Chanaday et al., 2021). Here, we do not expect STIM to be playing an inhibitory role to presynaptic Ca^2+^, as we observed a decrease in STIM levels and foci in *Rtnl1* mutants (Fig. 4), as well as a decrease in evoked Ca^2+^ fluxes, which were rescued by overexpressing STIM (Fig. 5). Work in non-neuronal cells indicate that STIM1/STIM2 promotes Ca^2+^ entry via coupling with Ca^2+^ channels on the PM (Berna-Erro et al., 2009; Chanaday et al., 2021; Gudlur et al., 2018; Jennette et al., 2022), and our *in vivo* data support a similar relationship in motor neurons for the *Drosophila* STIM1/STIM2 ortholog STIM. Accordingly, we propose a model for presynaptic compartments, whereby narrow tubular ER modulates Ca^2+^ entry via STIM (Fig. 10). Other components of the network with higher volume, such as cisternae, would work as a Ca^2+^ store, since both ER volume (Fig. 3) and resting ER Ca^2+^ levels (Fig. 6, 7) are unaffected in *Rtnl1* mutants. Evoked Ca^2+^ uptake by the ER was also decreased by loss of *Rtnl1* (Fig. 6, 7). While most lumenal GFP::HDEL or GCaMP, and hence Ca^2+^, appears to be in cisternae that would act as Ca^2+^ reservoirs, evoked ER Ca^2+^ uptake is reduced upon depletion of the tubular ER network in both phasic and tonic firing boutons. Moreover, we see the tubular network influencing the physiological profile of ER Ca^2+^, with ER Ca^2+^ release occurring on stimulation of *WT* Is boutons, and rarely on stimulation of *WT* Ib boutons. However, with loss of *Rtnl1* and depletion of tubular ER, ER in Ib boutons released Ca^2+^ during neuronal activity in over half of NMJs tested, and thus caused it to act like smaller Is boutons. Our results demonstrate a complex relationship between ER structure, and uptake and release of Ca^2+^ by the ER, suggesting that intact ER cisternae are not sufficient to maintain full levels of synaptic ER Ca^2+^ release and uptake, but that ER tubules are necessary as well.

Rtnl1 loss also decreased evoked presynaptic Ca^2+^ uptake by mitochondria (Fig. 8), similar to the ER and cytosolic compartments. As mitochondrial Ca^2+^ and ATP production are positively correlated (Datta & Jaiswal, 2021), energy generation in both Type Is and Ib boutons might potentially be impaired on depletion of tubular ER. Mitochondria normally receive Ca^2+^ directly from ER (de Brito and Scorrano, 2010; Hayashi et al., 2009; Rowland and Voeltz, 2012; Vance, 2014; Lee and Min, 2018), although synaptic mitochondria can take up Ca^2+^ from the cytosol (Ashrafi et al., 2020; Chouhan et al., 2010, 2012). However the comparative kinetics of the Ca^2+^ responses of *WT* cytosol, ER and mitochondria (Fig. 9), do not allow us to distinguish between mitochondrial Ca^2+^ uptake from the cytosol or from ER in this situation.

Beyond the Ca^2+^ handling defects characterized in this work, we propose *Rtnl1* mutants as a useful model to explore other biological roles of presynaptic ER, and to understand which presynaptic ER-related processes might be relevant to the mechanisms of HSP. In addition to Ca^2+^ handling, presynaptic ER depletion might affect other cellular processes such as autophagy, vesicle trafficking, glucose transport or lipid metabolism (Öztürk et al., 2020). Although we do not observe any PI(4,5)P_2_ alterations in *Rtnl1* mutants (Supplementary Fig. S6), it is still possible that other ER-dependent lipids might be affected, since at least some ER-PM contacts seem to be reduced (according to the decreased levels observed for STIM1 foci). Failed lipid transfer at ER-PM contacts has been previously related with reduced expansion of the growth cone and with neuronal PM growth in general (Petkovic et al., 2014), but we did not observe any defect in morphology of mature motor neurons in *Rtnl1* mutants, apart from a slight increase in the number of boutons.

Our results reveal at least three aspects of the *Rtnl1* mutant phenotype that could be relevant to the disease mechanisms of HSPs caused by mutations in ER-shaping proteins. First, *Rtnl1* mutants show a reduced frequency of miniature neurotransmission. This induces bouton fragmentation in young adult *Drosophila* so as to resemble that of aged *Drosophila*, whereas elevated minis can delay age-associated fragmentation and prolong motor ability in adult *Drosophila* (Banerjee et al., 2021). Second, *Rtnl1* mutants show decreased evoked cytosolic Ca^2+^ influx and neurotransmission at NMJs, which controls muscle contraction. Third, decreased evoked mitochondrial Ca^2+^ fluxes predict lower ATP generation capacity and potential energy deficits, although these may be mitigated by the reduced levels of synaptic transmission. Our results suggest mechanisms whereby altered organization of ER tubules could be a potential mechanism for diseases including HSP by affecting synaptic Ca^2+^ handling, and thus synapse function.

## METHODS

### *Drosophila* genetics

*Drosophila* stocks (Supplementary Table 1) were maintained on standard culture media at 18°C (Fly Facility from Department of Genetics, University of Cambridge). Crosses used to generate data were performed at 25°C and are listed in Supplementary Table 2.

#### Generation and analysis of *Rtnl1* and *Rtnl1::YFP* mutant alleles

disruption of *Rtnl1* was performed by expressing Cas9 together with two guide RNAs (*gRNAs*) that target the genomic sequences encoding Rtnl1 intramembrane domains (Fig. 1; Supplementary Fig. 1A). We used the online tool FIND CRISPRs (2017 version) (https://www.flyrnai.org/crispr/) from DRSC/TRiP Functional Genomics Resources (Harvard Medical School) for *gRNA* designing, targeting *Rtnl1* and filtering for at least 4 mismatches to any off-target sequences. BLAST sequence searches at www.ncbi.nlm.nih.gov were used to validate the absence of potential off-target sequences for the selected *gRNAs. gRNA* sequences were cloned into the tRNA::gRNA plasmid *pCFD5* by following the protocol on http://www.crisprflydesign.org/. In brief, *pCFD5* was digested with Bbs1-HF; DNA fragments containing *gRNAs* were generated using Q5 High-Fidelity DNA Polymerase with the guided sequences *CAGTGGAATCCCTTATCTAC* (1) and *CAAGTTCGGCGTCATTCTGT* (2); Gibson Assembly Master Mix was used to ligate the generated DNA fragments with digested *pCDF5*; High efficiency Transformation Protocol was used to establish bacterial colonies containing the assembled plasmids; colonies containing inserts with the correct size were sent for sequencing (Source BioSciences) to validate insert orientation. The resulting *pCFD5-2xgRNA* expression plasmid was purified using a QIAprep Spin Miniprep kit, and injected into *Drosophila* embryos (microinjection service at Department of Genetics, University of Cambridge). *Rtnl1*^*18*^ *and Rtnl1*^*4*^ were recovered from microinjection into embryos containing *nos-Cas9* on the X chromosome; we used the 2^nd^ chromosome of this stock as *Rtnl1*^*+*^ *WT*, as the parental chromosome of the *Rtnl1*^*18*^ *and Rtnl1*^*4*^ alleles. Surviving F0 male adults were individually crossed with *If/CyO* virgin females, and the male F1 adults from this cross were crossed with *Sco/CyO* virgin females for a few days, to establish stocks containing any new *Rtnl1* mutations. These males were then sacrificed for genomic DNA extraction and mutations in *Rtnl1* locus were analysed by PCR using Surveyor Mutation Detection Kit with Phusion High-Fidelity Polymerase and *GGCAAGGTAAACAGCGAGAC* (*Rtnl1-1A*) and *TGTGTATGGTGGACAAAAGCA* (*Rtnl1-1B*) primers (*WT* amplicon, 629 bp; Thermo Fisher Scientific) (Supplementary Fig. S1A). The same DNA polymerase and primers were used to generate PCR products from the mutated lines for sequencing (Source BioScience). To generate *Rtnl1*^*+*^ line, a single male from the *nos-Cas9* stock was crossed with *If/CyO* virgin females, and the *Rtnl1*^*+*^/*CyO, Tb* males from the progeny were crossed again with *If/CyO* virgin females to establish the stock.

To generate *Rtnl1::YFP*^*3*^, we first established a stable line expressing the *Rtnl1 gRNAs*. The *pCDF5-2xgRNA* plasmid was injected into *Drosophila* embryos of genotype *nos-phiC31 v*^*-*^ on the X chromosome, and the *attP2* landing site on the 3^*rd*^ chromosome. Successful phiC31 integrase-mediated integration of *pCDF5*, which contains an *attB* site, was verified by *v*^*+*^ in the resulting F1 adult flies. The *Rtnl1* double *gRNA* line was then used with *nos-Cas9* to generate *Rtnl1* mutations in the germline of *Rtnl1*^*CPTI001291*^ flies.

For RT-PCR, we used a One-Step RT-PCR kit, on total RNA template from a TRIzol reagent extraction (15 males per genotype), and *GGCAAGGTAAACAGCGAGAC* (forward) and *GTTCAAACCCACTGTCCAGG* (reverse) primers (Supplementary Fig. S2A), which do not amplify *Rtnl1*^*1*^ (negative control), and amplify only *WT* mRNA (636 bp) but not genomic DNA (13955 bp).

Unless otherwise specified, PCRs were performed with DreamTaq Green PCR Master Mix. Gel electrophoresis was performed in 2% agarose in TBE. DNA fragment sizes were estimated using a GeneRuler 1 kb Plus DNA Ladder. Primers were designed with Primer3web 4.1.0 (https://primer3.ut.ee). Cloning design and sequence analysis were done with SnapGene 4.2.11. Bacterial cultures were grown using BD Difco LB Broth, Miller and BD Bacto Agar.

#### Genotyping *Rtnl1* alleles

We used PCR primer pairs to verify the orientation of the region inverted in *Rtnl1*^*18*^ (Fig. 1A; Supplementary Fig. S1A). Primers *GGCAAGGTAAACAGCGAGAC* (*Rtnl1-1A*) and *CTGTCGCACGAAAAGGTACA* (*Rtnl1-2B*) gave a product of 272 bp in *WT*, but none in *Rtnl1*^*18*^. Primers *GGCAAGGTAAACAGCGAGAC* (*Rtnl1-1A*) and *GAGGGTTAGGAGCGACAAGT* (*Rtnl1-2A*) primers (both forward in *WT*) gave a PCR product of 242 bp in *Rtnl1*^*18*^, but none in *WT. GGCAAGGTAAACAGCGAGAC* (*Rtnl1-1A*) and *CTGTCGCACGAAAAGGTACA* (*Rtnl1-2B*), were also used to detect *UAS-Rtnl1::HA*, which gave a 207 bp band (lacking a 65 bp intron) compared to the 272 bp band of *WT* genomic DNA. To genotype *Rtnl1*^*1*^, *GGAAATTGCGTGGAACTCAT* and *TATTCGCATTTCCTCGATCC* primers gave a PCR product of 5623 bp in *WT*, and 496 bp in *Rtnl1*^*1*^ (Yalçın et al., 2017). New stocks were genotyped for *Rtnl1* alleles and *UAS-Rtnl1::HA* after they were constructed to verify their genotypes, and at intervals throughout the work.

#### Generation of mitochondrial Ca^2+^ sensor

*17xUASTattB-CEPIA3mt::myc* was constructed as for the lumenal ER sensors (Oliva et al., 2020), and integrated by injection at phiC31 landing site *attP86Fb* (Bischof et al., 2007).

### Histology and confocal microscopy

Third instar larvae were dissected in chilled Ca^2+^-free HL3 solution (prefixation HL3) (Stewart et al., 1994), and fixed for 10 min in PBS with 4% formaldehyde. Unless otherwise specified, visualization of fluorescent tags was performed via direct imaging, without immunostaining. For immunostaining, the dissected preparations were permeabilized in PBS containing 0.1% Triton X-100 (PBT) at room temperature for 1 h. After permeabilization, samples were blocked in PBT with 4% bovine serum albumin for 30 min at room temperature, incubated with primary antibodies (Supplementary Table 3) overnight at 4ºC, and finally incubated with secondary antibodies (Supplementary Table 3) for 2 h at room temperature. For myc and HA immunostaining, samples were permeabilized for 30 min with PBT, blocked for 1 h with 5% Normal Goat Serum in PBT, and incubated with anti-myc or anti-HA overnight at 4ºC. Processed preparations were mounted in Vectashield, and images were collected using EZ-C1 acquisition software (Nikon) on a Nikon Eclipse C1si confocal microscope (Nikon Instruments, UK) with 488 nm (for GFP and Alexa-488 signals), 561 nm (for tdTom, mCherry and Alexa-594 signals) and 638 nm (for Alexa-647 signal) lasers. Images were captured using a 40x/1.3NA oil Nikon Plan Fluor DIC H/N2 infinity-0.17 WD 0.2 objective, or a 60x/1.4NA oil Plan Apo VC infinity-0.17 DIC N2.

### Live Imaging

Wandering third instar larvae were fillet dissected in ice-cold Schneider’s *Drosophila* medium and imaged in ice-cold HL3 of the appropriate kind. Presynaptic responses were imaged in low-Mg^2+^ HL3 containing 1 mM Ca^2+^ (Oliva et al., 2020). Postsynaptic responses were imaged in low-Mg^2+^ HL3, with 1 mM L-glutamic acid to partially saturate postsynaptic receptors; higher concentrations of L-glutamic acid blocked postsynaptic responses, and lower concentrations did not inhibit muscle contractions enough. Muscle contractions often obscured postsynaptic responses even with 1 mM L-glutamic acid. New HL3 was kept no longer than 3 days at 4ºC, as Ca^2+^ responses became unreliable with HL3 older than this.

Nerves were cut at the base of the VNC with dissection scissors, suctioned into a heat-polished glass pipette (Macleod et al., 2002), attached to a stimulator and train generator, and imaged on an upright widefield Olympus BX50 microscope as described previously (Oliva et al., 2020) using LED illumination (Cairn Instruments): a 470nm LED with an ET470/40x T495LPXR excitation cube, a cool white LED with an ET572/X35 holder and ET500/20xT485/68dcrb excitation cube, a 59022bs infinity cube, and an Cairn Optosplit II beamsplitter with an ET520/40m T565LPXR-UF2 ET632/60m emission cube. ER lumen and mitochondrial responses were imaged at 10 frames per second, and EM gain level 100. Presynaptic cytosolic and postsynaptic responses were imaged at 50 frames per second and EM gain level 100. Live images were acquired using a 40x/1.0NA long working distance, water-immersion objective (W Plan-Apochromat 40x/1.0 DIC M27), a 2x C-mount fixed focal length lens extender (Cairn, Faversham, UK), and an Andor EMCCD camera (iXon Ultra model 897_BV, 512×512 pixels, Andor Technology, Belfast, UK). Imaging data were acquired using Micro-Manager (Edelstein et al., 2014) and saved as multilayer TIFF files.

### Image analysis and figure preparation

Confocal stack images (.nd2) were processed with Fiji ImageJ (Schindelin et al., 2012). Selection of specific z-stack ranges, generation of maximum intensity projections, brightness adjustment (identical for all images within the same experiment), and/or image cropping were performed as required. Thresholding of the resulting images was used to select regions of interest (ROIs) for quantification. For overall axon and NMJ structures, thresholding was manually adjusted on the PM marker channel (when present), and the resulting ROIs were applied to the remaining channels. For punctate signals, to reduce variability in ROI selection between samples, one of the predefined Fiji threshold algorithms was chosen, based on its ability to filter puncta previously identified by eye in multiple random images. The selected ROIs were used to measure mean pixel intensity and/or area values, which were recorded in a Microsoft Excel 2019 file (.xlsx). Files generated during successive steps in image processing, the threshold algorithms used in each experiment (when required), the threshold values used for all ROIs, and the quantification datasets, can be found in the underpinning dataset for this paper.

Wide-field multilayer tif time course files were opened in Fiji ImageJ (Schindelin et al., 2012), and channels were background subtracted (Rolling Ball Radius: 50 pixels). NMJs were stabilized using Register ROI from the cookbook menu with an ROI containing the entire NMJ. Some NMJs needed to be registered multiple times. Fluorescent intensity time traces were obtained for ROIs traced around the entire NMJ, or around all individual mitochondria in the NMJ, in each tif frame in a given data set using the Time Series Analyzer V3 plugin and ROI manager, recording the average fluorescence in each ROI (or across all mitochondria ROIs). Mitochondria or NMJs that moved and could not be corrected with registration software, or drifted out of focus during recording, were discarded from analysis. If only one or two mitochondria/boutons in the NMJ were available for analysis, due to focus or drift, data were discarded from analysis. Miniature neurotransmission frequencies were manually counted (Supplementary Fig. S8) at the distal bouton over a 20 second interval. Miniature neurotransmission amplitude was calculated via a maximum ΔF/F value reached at the distal bouton over a 20 second interval in R. Time traces were saved as CSV files and NMJ averages were fed into R scripts (R Core Team, 2020) to perform bleach correction and rolling averages (5 frames). Bleach correction was performed by fitting a bleach correction power curve (PC = a*x^b^) to pre-stimulation fluorescence (a and b coefficients calculated in R), and dividing raw fluorescence over the entire time course by this fitted curve. R scripts were also used to collect resting fluorescence (immediate pre-stimulation value of the fitted power curve), maximum fluorescence (maximum raw fluorescent value), ΔF, ΔF/F, time to peak, time to 50% recovery, and, with the cytosolic sensor, time to 100% recovery. All files generated during successive steps in image processing, ROIs, and the quantification analyses, are available in the dataset for this paper.

### Electron microscopy

#### Fixation and embedding

Fixation of dissected third instar larvae was done by replacing HL3 solution with fix solution (0.05 M sodium cacodylate at pH 7.4 containing 4% formaldehyde, 2% vacuum distilled glutaraldehyde, and 2 mM CaCl_2_). The larvae were incubated at 4°C overnight. After fixation the preparations were washed 3 times for 10 minutes each at 4°C using 0.05 M cold cacodylate buffer. Samples were osmicated in 1% osmium tetroxide/1.5 % potassium ferricyanide/0.05 M sodium cacodylate buffer for 3d at 4ºC. The preparations were washed four times with deionized water at room temperature. Thiocarbohydrazide solution was prepared at a concentration of 0.1% in deionized water, incubated in a 60°C oven for 1 hour while agitated by swirling every 10 minutes to facilitate dissolution, and filtered through two 9 cm filter papers just before use. The preparations were then incubated in thiocarbohydrazide solution for 20-30 minutes at room temperature and covered with aluminum foil to protect from light. They were then rinsed with deionized water at room temperature 5 times for 3 minutes, incubated in 2% aqueous osmium tetroxide for 30-60 minutes at room temperature, and rinsed with deionized water at room temperature 5 times for 3 minutes. Afterwards the preparations were incubated in 2% uranyl acetate (maleate-buffered to pH 5.5) at 4°C for 3 days, and rinsed with deionized water at room temperature 5 times for 3 minutes. They were then incubated in lead aspartate solution (0.66 g of lead nitrate was dissolved in 100 ml 0.03 M aspartic acid solution and pH adjusted to 5.5 with 1 M KOH, in a 60°C oven for 30 minutes) at 60°C for 30 minutes, and rinsed with deionized water at room temperature 5 times for 3 minutes. They were then dehydrated twice with each of 50%, 70%, 90% and 100% ethanol and twice with dried ethanol, twice with dried acetone, and twice with dry acetonitrile. The preparations were incubated in 50/50 acetonitrile/Quetol 651 overnight at room temperature. They were then incubated 72 hours in Quetol epoxy resin 651 (Agar Scientific, Stansted, UK) without BDMA and five times for 24 hours in Quetolepoxy resin 651 with BDMA (dimethylbenzylamine). Afterwards they were incubated at 60°C for a minimum of 48 hours.

#### Sectioning

The resin blocks were sectioned using a Leica Ultracut E ultramicrotome. Sections were mounted on aluminum SEM stubs using carbon sticky pads and coated with 30 nm carbon for conductivity. The sections were imaged in a Verios 460 SEM at 4keV and 0.2 nM probe current using the concentric backscatter detector at low magnification, and stitched image maps were acquired using MAPS automated acquisition software to give an overview through the central region of the larva. Once NMJs were identified, blocks were trimmed down to the region containing the structures of interest. The samples were then sectioned with a 4 mm UltraMaxi Diatome 35° knife using an ATUMtome (RMC/ Boeckeler Instruments) at a thickness of 50 nm and were collected onto rolling Kapton tape. Kapton tape strips were cut and mounted on 4-inch silicon wafers using double-sided carbon sticky tape. Wafers were then sputter coated with 30 nm carbon using a Quorum Q150 T E carbon coater. Wafers were imaged in a Verios 460 scanning EM (FEI/Thermofisher) at 4 keV accelerating voltage and 0.2 nA probe current in backscatter mode using the concentric backscatter detector (CBS) in immersion mode at a working distance of 3.5-4 mm; 1536 × 1024 pixel resolution, 3 µs dwell time, four line integrations. Stitched maps were acquired using FEI MAPS software using the default stitching profile and 5% tile overlap.

#### Imaging

Kapton tape strips were mounted on 4-inch silicon wafers using double-sided carbon sticky tape. Wafers were then sputter coated with 30 nm carbon using a Quorum Q150 T E carbon coater. Wafers were imaged in a Verios 460 scanning electron microscope (FEI/Thermofisher) at 4 keV accelerating voltage and 0.2 nA probe current in backscatter mode using the concentric backscatter detector (CBS) in immersion mode at a working distance of 3.5-4 mm; 1536 × 1024 pixel resolution, 3 us dwell time, 4 line integrations. Stitched maps were acquired using FEI MAPS software using the default stitching profile and 5% tile overlap. Images of serial sections were imported into a single file using ImageJ/Fiji (Schindelin et al., 2012), and sections were aligned and reconstructed using the TrakEM2 plugin (Cardona et al., 2012). Reconstructions were further processed with 8 smoothing steps of Laplacian smooth function in the program MeshLab (http://meshlab.sourceforge.net). The rendering file generated is available in the underpinning data for the paper.

### Statistical analysis

After quantification, values were exported from .xlsx or .CSV to .txt files to be analysed with R Studio 1.3.1093, or GraphPad Prism 9 for statistical analysis and plotting. Figures were made using Adobe Illustrator CC 2017 or Affinity Designer 1.10.0. All R scripts were written by the authors and are available in the underpinning data for the paper.

A Shapiro-Wilk test was used to test for normality in the data distribution, and Levene’s test was used to test for differences between group variances. Normally distributed data were analyzed with mixed-effects model repeated-measures ANOVA or unpaired two-tailed Student’s T-tests. Data not normally distributed were analyzed using non-parametric Kruskal-Wallis or Mann-Whitney U tests. Post-hoc multiple comparisons were then applied where relevant. Tukey HSD test was used to compare each group with the other groups, while Dunnett’s test was used for planned comparisons between every group and a single control group.

## ACKNOWLEDGEMENTS

We thank Dion Dickman, Xun Huang, Andrea Daga, Guy Tear, Addgene, the Developmental Studies Hybridoma Bank, the Bloomington and Kyoto *Drosophila* Stock Centers, and the University of Cambridge Department of Genetics Fly Facility, for antibodies, constructs and stocks. We thank the University of Cambridge Department of Genetics Fly Facility for embryo injections, and the electron microscopy facility of the Cambridge Advanced Imaging Centre. We thank Beatriz Ibañez for helpful discussions, and the HSP Tom Wahlig Foundation for its support and dissemination of our work. The authors declare no competing financial interests.

## FUNDING

This work was supported by grants from the Medical Research Council (MRC) (MR/S011226/1), the Biotechnology and Biological Sciences Research Council (BBSRC) (BB/S001212/1), and the Spastic Paraplegia Foundation, Inc (SPF) to CJO’K. JJPM was supported by the SPF, a Marie-Sklodowska-Curie grant from the European Union (745007), and a Juan de la Cierva Incorporación grant (IJC2019-038819-I) from the Spanish State Research Agency (MCIN/AEI/10.13039/501100011033). MKO was supported by a Marie-Sklodowska-Curie grant from the European Union (660516), and SO by the SPF.

## AUTHOR CONTRIBUTIONS

JJPM: Conceptualization, Resources, Formal analysis, Funding acquisition, Investigation, Visualization, Writing - original draft, Writing - review and editing; generated and characterized new *Rtnl1* alleles, performed and analyzed most confocal microscopy, participated in the generation and imaging of EM samples. RCS: Conceptualization, Resources, Formal analysis, Investigation, Visualization, Writing - original draft, Writing - review and editing; performed and analysed most Ca^2+^ imaging, performed and analyzed confocal microscopy. MKO: Conceptualization, Resources, Formal analysis, Investigation, Visualization, Writing - review and editing; generated and characterized GCaMPs, performed Ib ER Ca^2+^ imaging. SO: Serial EM analysis. FG: generation of EM samples and serial EM sectioning, Writing - review and editing; KM: EM imaging, Writing - review and editing. CJO’K: Conceptualization, Resources, Formal analysis, Supervision, Funding acquisition, Writing - original draft, Project administration, Writing - review and editing.

## Online supplemental material

**Supp. Fig. S1** shows CRISPR-derived lesions in *Rtnl1* and their effects on the *Rtnl1* coding region. **Supp. Fig. S2** shows *Rtnl1* expression in *Rtnl1* mutant CRISPR alleles. **Supp. Fig. S3** shows high-magnification confocal planes of the presynaptic ER network organization in *Rtnl1* mutants. **Supp. Fig. S4** (tdTom::Sec61β) and **Supp. Fig. S5** (GFP::HDEL) show extended data on the effects of *Rtnl1* mutant alleles on presynaptic ER distribution. **Supp. Fig. S6** shows presynaptic PI(4,5)P_2_ levels in *Rtnl1* mutants. **Supp. Fig. S7** shows synaptic transmission in *Rtnl1* mutants. **Supp. Fig. S8** shows images of miniature neurotransmission in *Rtnl1* mutants. **Supp. Fig. S9** and **Supp. Fig. S10** show extended data on cytosolic Ca^2+^ handling in Is and Ib boutons, respectively, of *Rtnl1* mutants. **Supp. Fig. S11** shows extended data on ER Ca^2+^ handling in Is and Ib boutons of *Rtnl1* mutants. **Supp. Fig. S12** and **Supp. Fig. S13** show extended data on mitochondrial Ca^2+^ handling in Is and Ib boutons, respectively, of *Rtnl1* mutants. **Video 1** (*WT*) and **Video 2** (*Rtnl1*) show postsynaptic Ca^2+^ response to low-frequency stimulation. **Video 3** (*WT*) and **Video 4** (*Rtnl1*) show postsynaptic miniature Ca^2+^ events. **Video 5** (*WT*) and **Video 6** (*Rtnl1*) show cytosolic Ca^2+^ response to stimulation. **Video 7** (*WT*) and **Video 8** (*Rtnl1*) show ER Ca^2+^ response to stimulation in Is boutons. **Video 9** (*WT*) and **Video 10** and Video 11 (*Rtnl1*) show ER Ca^2+^ response to stimulation in Ib boutons. We provide two versions of the Rtnl1 video, the first (Video 10) with a brightness comparable to the *WT* **Video 9**, and the other (Video 11) with increased brightness for viewer clarity. **Video 12** (*WT*) and **Video 13** (*Rtnl1*) show mitochondria Ca^2+^ response to stimulation in Ib boutons. **Supp. Table 1** shows the *Drosophila* stocks used in this work. **Supp. Table 2** shows the crosses used in this work. **Supp. Table 3** shows the reagents used in this work.

### Data availability

The underpinning dataset for this article is openly available at the University of Cambridge Data Repository (https://www.repository.cam.ac.uk): https://doi.org/10.17863/CAM.93878.

## Notes

https://doi.org/10.17863/CAM.93878

### Competing Interest Statement

The authors have declared no competing interest.

### Summary of Updates

The whole manuscript text (including the title) has been revised to clarify the aim and the conclusions of the article, as well as to describe the new data included. In particular, Figures 3 and 5 include new data regarding ER ultrastructural analysis and STIM-mediated rescue experiments, respectively. New authors have been added, and the affiliation of one of the existing authors has been updated. Finally, all Figures and Supplemental files have been updated to improve quality and format.

