## Supplementary Table 1 for "*Drosophila* SPG12 ortholog, reticulon-like 1, governs presynaptic ER organization and Ca^2+^ dynamics"

**Supplementary Table 1. *Drosophila* stocks used or generated in this work.** References are shown where possible; unpublished stocks that we constructed are listed as "this work". The source of each stock is listed as an RRID identifier where possible. Any figure panels that present data from each stock are listed. Where necessary, the *GAL4* driver, Ca$^{2+}$ sensor, *UAS*, and recombinant stocks listed were constructed as two alternative versions not shown in the Table, one with a *WT* second chromosome, one with a *Rtnl1* mutant second chromosome, usually *Rtnl1*$^{18}$.

| ***Drosophila* stock genotype (Chr)** | **Reference** | **Source/RRID** | **Notes** | **Figs** |
|---|---|---|---|---|
| **Toolkit to generate new strains** | | | | |
| *SdhA$^{1110}$/CyO, P{GAL4-Kr.C}DC3, P{UAS-GFP.S65T}DC7* | (Casso et al., 1999) | BDSC_51659 | *2$^{nd}$ chr-balanced line* | ----- |
| *CyO, P{Tb$^{1}$}Cpr$^{CyO-A}$/sna$^{Sco}$ (II)* | (Lattao et al., 2011) | DGGR_36335 | *2$^{nd}$ chr-balanced line* | ----- |
| *w ; In(3LR)TM3, ri p$^{p}$ sep bx$^{34e}$ e Ser/ In(3LR)TM6B, Tb Hu* | ----- | Simon Collier Cambridge University fly facility | *3$^{rd}$ chr-balanced line* | ----- |
| *y$^{1}$ M{nos-Cas9.P}ZH-2A w* (X)* | (Port et al., 2014) | DGGR_54591 | *germline-expressed Cas9* | ----- |
| *U6-Rtnl1-gRNAs(attP2) (III)* | This work | ----- | *ubiquitously expressed gRNAs against Rtnl1* | ----- |
| *attP2$^{+}$ P{nos-phiC31\int.NLS}X #12;; P{CaryP}attP2) (X)* | ----- | Simon Collier Cambridge University fly facility | attP2 injection stock; attP2 *WT* background "*attP2$^{+}$*" | 2, 5, 8 |
| *P{nos-phiC31\int.NLS}X, y$^{1}$ sc$^{1}$ v$^{1}$ sev$^{21}$ ; ; P{CaryP}attP2 (X)* | (Bischof et al., 2007) | BDSC_25710 | *germline expressed phiC31 integrase and attP2 site (used for generation of U6-Rtnl1-gRNA line)* | ----- |
| ***Rtnl1* alleles** | | | | |
| *Rtnl1$^{1}$ (II)* | (Wakefield & Tear, 2006) | Guy Tear | *Rtnl1* mutant allele generated by imprecise *P*-element excision | 1, S2, S9, S12 |
| *Rtnl1$^{+}$ (II)* | This work | ----- | *Rtnl1* wildtype chromosome on which CRISPR mutants were made. Referred to as *wildtype* (*WT*) throughout text | ----- |
| *Rtnl1$^{4}$ (II)* | This work | ----- | *Rtnl1* CRISPR mutant allele | 1, S1 |
| *Rtnl1$^{18}$ (II)* | This work | ----- | *Rtnl1* CRISPR mutant allele | 1, 3, S1, S2 |
| *Rtnl1::YFP(PBac{681.P.FSVS-1}Rtnl1$^{CPTI001291}$ (II)* | (O'Sullivan et al., 2012) | DGRC_115146 | YFP-tagged Rtnl1 exon trap line | 1, S2 |
| *Rtnl1::YFP$^{3}$ (II)* | This work | ----- | *Rtnl1::YFP* CRISPR mutant allele | 1, S1, S2 |
| ***GAL4* drivers** | | | | |
| *Ib-GAL4(P{GMR94G06-GAL4}attP2) (III)* | (Pérez-Moreno & O'Kane, 2019) | BDSC_40701 | Type Ib MN driver | S5 |
| *Is-GAL4(P{GMR27E09-GAL4}attP2) (III)* | (Pérez-Moreno & O'Kane, 2019) | BDSC_49227 | Type Is MN driver | ----- |
| *P{GawB}D42 (III)* | *(Yeh et al., 1995)* | BDSC_8816 | MN driver | ----- |
| **Ca$^{2+}$ sensors** | | | | |
| *P{Mhc-SynapGCaMP6f}3-5 (III)* | (Newman et al., 2017) | BDSC_67739 | Postsynaptic Ca$^{2+}$ sensor | |
| *P{UAS-myr::GCaMP6s} (III)* | ----- | Dion Dickman, unpublished | Cytosolic Ca$^{2+}$ sensor | |
| *17xUASTattB-ER-GCaMP6-210(attP86Fb) (III)* | (Oliva et al., 2020) | BDSC_91397 | ER lumen Ca$^{2+}$ sensor | |

| | | | | |
|---|---|---|---|---|
| *17xUASTattB-CEPIA3mt::myc(attP86Fb) (III)* | This work | Construct from (Suzuki et al., 2014) | Mitochondria matrix $Ca^{2+}$ sensor | |
| *UAS-tdTom-p2a-GCaMP56* | (Daniels et al., 2014) | BDSC_80079 | Ratiometric $Ca^{2+}$ sensor | |
| ***UAS* constructs (non-$Ca^{2+}$ sensors)** | | | | |
| *PBac{UAS-CD4-tdGFP}VK00033 (III)* | (Han et al., 2011) | BDSC_35836 | PM marker | |
| *P{UAS-CD4-tdTom}7M1 (III)* | (Han et al., 2011) | BDSC_35841 | PM marker | |
| *P{UAS-Rtnl1::GFP}3 (III)* | (Rao et al., 2016) | BDSC_77908 | GFP-tagged *Rtnl1* | ----- |
| *UAS-Rtnl1::HA(attP2) (III)* | (Summerville et al., 2016) | Andrea Daga | HA-tagged *Rtnl1* | 2, 5, 8 |
| *PBac{20XUAS-tdTom-Sec61beta}VK00037 (II)* | (Summerville et al., 2016) | BDSC_64746 | ER membrane marker | S5 |
| *P{20XUAS-tdTom-Sec61beta}attP2 (III)* | (Summerville et al., 2016) | BDSC_64747 | ER membrane marker | 2, 5, 6, 7, 8, S3, S4, S7, S8, S9, S10, S11, S12, S13 |
| *UAS-Sturkopf::GFP (III)* *(formerly UAS-CG9186::GFP)* | (Thiel et al., 2013) | BDSC_91394 | ER membrane marker | ----- |
| *P{UAS-BiP-sfGFP-HDEL}attP2 (III)* | (Summerville et al., 2016) | BDSC_64749 | ER lumen marker | 3, S5 |
| *$L^1$/CyO ; P{UAS-PLCdelta-PH-EGFP}3/ TM6B, $Tb^1$* | (Verstreken et al., 2009) | BDSC_39693 | GFP-tagged $PI(4,5)P_2$ sensor | S6 |
| *UAS-STIM::mCherry(M{3xP3-RFP.attP}ZH-51D) / TM6c* | (Bi et al., 2014) | Xun Huang | mCherry-tagged STIM | 4 |
| **Recombinant chromosomes** | | | | |
| *Ib-GAL4(attP2), P{UAS-CD4-tdTom}7M1 (III)* | This work | ----- | Type Ib MN driver with PM marker | 3, S6 |
| *Ib-GAL4(attP2), PBac{UAS-CD4-tdGFP}VK00033 (III)* | This work | ----- | Type Ib MN driver with PM marker | 2, S3 |
| *Is-GAL4(attP2), PBac{UAS-CD4-tdGFP}VK00033 (III)* | This work | ----- | Type Is MN driver with PM marker | 2 |
| *Ib-GAL4(attP2), UAS-Sturkopf::GFP (III)* | This work | ----- | Type Ib MN driver with ER membrane marker | 2, 4, S3 |
| *Ib-GAL4(attP2), P{Mhc-SynapGCaMP6f}3-5 (III)* | This work | ----- | Type Ib MN driver with postsynaptic $Ca^{2+}$ sensor | ------- |
| *Is-GAL4(attP2), P{Mhc-SynapGCaMP6f}3-5 (III)* | This work | ----- | Type Is MN driver with postsynaptic $Ca^{2+}$ sensor | S7 |
| *Ib-GAL4(attP2), P{UAS-myr::GCaMP6s} (III)* | This work | ----- | Type Ib MN driver with cytosolic $Ca^{2+}$ sensor | 5, S10 |
| *Is-GAL4(attP2), P{UAS-myr::GCaMP6s} (III)* | This work | ----- | Type Is MN driver with cytosolic $Ca^{2+}$ sensor | 5, S9 |
| *Ib-GAL4(attP2), 17xUASTattB-ER-GCaMP6-210(attP86Fb) (III)* | (Oliva et al., 2020) | ----- | Type Ib MN driver with ER lumen $Ca^{2+}$ sensor | 7, S11 |
| *Is-GAL4(attP2), 17xUASTattB-ER-GCaMP6-210(attP86Fb) (III)* | (Oliva et al., 2020) | ----- | Type Is MN driver with ER lumen $Ca^{2+}$ sensor | 6, S11 |
| *Ib-GAL4(attP2), 17xUASTattB-CEPIA3mt::myc(attP86Fb) (III)* | This work | ----- | Type Ib MN driver with mitochondria $Ca^{2+}$ sensor | 8, S13 |
| *Is-GAL4(attP2), 17xUASTattB-CEPIA3mt::myc(attP86Fb) (III)* | This work | ----- | Type Is MN driver with mitochondria $Ca^{2+}$ sensor | 8, S12 |
| *P{GawB}D42, P{UAS-Rtnl1::GFP}3 (III)* | This work | ----- | MN driver with ectopic GFP-tagged copy of *Rtnl1* | S4 |
| *P{GawB}D42, UAS-tdTom::Sec61β(attP2) (III)* | This work | ----- | MN driver with PM marker | 2 |
| *P{GawB}D42, UAS-tdTom::Sec61β(attP2), UAS-CD4::tdGFP (III)* | This work | ----- | MN driver with ER and PM markers | S4 |
