## Supplementary Table 2 for "*Drosophila* SPG12 ortholog, reticulon-like 1, governs presynaptic ER organization and Ca^2+^ dynamics"

**Supplementary Table 2. Crosses used in this work.** The table shows simplified genotypes of the F0 parental stocks (from **Supp. Table 1**) used to generate the F1 larvae shown or analyzed in figures, as well as the F1 summary genotypes used. Floating balancers are shown in brackets. Where possible we selected non-balancer homozygous parents, otherwise we counterselected against the dominant balancer markers in F1 larvae: Tubby (on *CyO, Tb* or on *TM6B*) or the widely expressed GFP of *CyO::GFP*. Normally two alternative crosses were performed, one with both parents *WT (+)*, the other with both parents mutant for *Rtnl1*; the alternative alleles used are in brackets in F0 parental genotypes. In F1 larvae, the designation *Rtnl1*$^{x}$ represents any of the *Rtnl1* genotypes possible in the crosses shown.

| Fig | F0 parent stock 1 | F0 parent stock 2 | F1 Genotype |
|---|---|---|---|
| 2A, S3 | ( + OR Rtnl1$^{18}$); Ib-GAL4, UAS-CD4::tdGFP / (TM6B) | (+ OR Rtnl1$^{18}$) / (CyO::GFP); UAS-tdTom::Sec61β / (TM6B) | Rtnl1$^{x}$; Ib-GAL4, UAS-CD4::tdGFP / UAS-tdTom::Sec61β |
| 2B | ( + OR Rtnl1$^{18}$) ; Is-GAL4, UAS-CD4::tdGFP / (TM6B) | ( + OR Rtnl1$^{18}$)/(CyO::GFP); UAS-tdTom::Sec61β/(TM6B) | Rtnl1$^{x}$; Is-GAL4, UAS-CD4::tdGFP / UAS-tdTom::Sec61β |
| 2C | One of:<br>- ( + OR Rtnl1$^{18}$); + / (TM6B)<br>- ( + OR Rtnl1$^{18}$); UAS-Rtnl1::HA / (TM6B) | ( + OR Rtnl1$^{18}$) / (CyO::GFP); D42-GAL4, UAS-tdTom::Sec61β / (TM6B) | Rtnl1$^{x}$; D42-GAL4, UAS-tdTom::Sec61β / (+ OR UAS-Rtnl1::HA) |
| 2D-E, S3 | ( + OR Rtnl1$^{18}$) | ( + OR Rtnl1$^{18}$) / (CyO, Tb); Ib-GAL4, UAS-Sturkopf::GFP / (TM6B) | Rtnl1$^{x}$; Ib-GAL4, UAS-Sturkopf::GFP / + |
| 3B-E, S5C | ( + OR Rtnl1$^{18}$);<br>Ib-GAL4, UAS-CD4::tdTom / (TM6B) | ( + OR Rtnl1$^{18}$) / (CyO::GFP);<br>UAS-GFP::HDEL / (TM6B) | Rtnl1$^{x}$;<br>Ib-GAL4,<br>UAS-CD4::tdTom / UAS-GFP::HDEL |
| 4 | ( + OR Rtnl1$^{18}$) / (CyO, Tb);<br>Ib-GAL4, UAS-Sturkopf::GFP / (TM6B) | ( + OR Rtnl1$^{18}$) / (CyO, Tb);<br>Ib-GAL4,<br>UAS-STIM::mCherry / (TM6B) | Rtnl1$^{x}$;<br>Ib-GAL4, UAS-Sturkopf::GFP / UAS-STIM::mCherry |
| S4A | ( + OR Rtnl1$^{18}$) | ( + OR Rtnl1$^{18}$) / (CyO, Tb); D42-GAL4, UAS-tdTom::Sec61β, UAS-CD4::tdGFP / (TM6B) | Rtnl1$^{x}$; D42-GAL4, UAS-tdTom::Sec61β, UAS-CD4::tdGFP / + |
| S4B | ( + OR Rtnl1$^{18}$) | (Rtnl1$^{+}$ OR Rtnl1$^{1}$) / (CyO, Tb);<br>D42-GAL4, UAS-tdTom::Sec61β / (TM6B) | Rtnl1$^{x}$ ; D42-GAL4, UAS-tdTom::Sec61β / + |
| | (Rtnl1$^{+}$ OR Rtnl1$^{1}$) / (CyO, Tb); D42-GAL4, UAS-Rtnl1::GFP / (TM6B) | (Rtnl1$^{+}$ OR Rtnl1$^{18}$) / (CyO::GFP);<br>UAS-tdTom::Sec61β / (TM6B) | Rtnl1$^{x}$; D42-GAL4, UAS-Rtnl1::GFP / UAS-tdTom::Sec61β |
| S5A | Rtnl1$^{+}$ / (CyO::GFP); UAS-GFP::HDEL / (TM6B) | UAS-tdTom::Sec61β / (CyO, Tb);<br>Ib-GAL4 / (TM6B) | Rtnl1$^{+}$ / UAS-tdTom::Sec61β; UAS-GFP::HDEL / Ib-GAL4 |
| S5B | (Rtnl1$^{+}$) / (CyO::GFP) OR Rtnl1$^{1}$ / (CyO, Tb); UAS-GFP::HDEL / (TM6B) | (Rtnl1$^{+}$ OR Rtnl1$^{18}$) / (CyO, Tb); Ib-GAL4, UAS-CD4::tdGFP / (TM6B) | Rtnl1$^{x}$; UAS-GFP::HDEL / Ib-GAL4, UAS-CD4::tdGFP |
| S6 | (Rtnl1$^{+}$ OR Rtnl1$^{18}$);<br>Ib-GAL4, UAS-CD4::tdTom / (TM6B) | (Rtnl1$^{+}$ OR Rtnl1$^{18}$);<br>UAS-PLC□::PH::GFP / (TM6B) | Rtnl1$^{x}$;<br>Ib-GAL4, UAS-CD4::tdTom / UAS-PLC□::PH::GFP |
| S7, S8 | (Rtnl1$^{+}$ OR Rtnl1$^{18}$);<br>Is-GAL4, Mhc-SynapGCaMP6f / (TM6B) | (Rtnl1$^{+}$ OR Rtnl1$^{18}$) / (CyO::GFP);<br>UAS-tdTom::Sec61β / (TM6B) | Rtnl1$^{x}$; Is-GAL4,<br>Mhc-SynapGCaMP6f / UAS-tdTom::Sec61β |
| 6, S11 | (Rtnl1$^{+}$ OR Rtnl1$^{18}$) / (CyO::GFP); Is-GAL4, UAS-ER-GCaMP6-210 / (TM6B) | (Rtnl1$^{+}$ OR Rtnl1$^{18}$) / (CyO::GFP);<br>UAS-tdTom::Sec61β / (TM6B) | Rtnl1$^{x}$; Is-GAL4,<br>UAS-ER-GCaMP6-210 / UAS-tdTom::Sec61β |
| 7, S11 | (Rtnl1$^{+}$ OR Rtnl1$^{18}$) / (CyO::GFP); Ib-GAL4, UAS-ER-GCaMP6-210 / (TM6B) | (Rtnl1$^{+}$ OR Rtnl1$^{18}$) / (CyO::GFP);<br>UAS-tdTom::Sec61β / (TM6B) | Rtnl1$^{x}$; Ib-GAL4,<br>UAS-ER-GCaMP6-210 / UAS-tdTom::Sec61β |

| | | | |
|---|---|---|---|
| 5 | *(Rtnl1$^{+}$ OR Rtnl1$^{18}$) / (CyO,Tb); / UAS-tdTom-p2a-GCaMP56 / (TM6B)* | *(Rtnl1$^{+}$ OR Rtnl1$^{18}$)/(CyO,Tb); / Ib-Gal4 / (TM6B)* | *Rtnl1$^{x}$; UAS-tdTom-p2a-GCaMP56/ Ib-GAL4* |
| 5, S10 | *(Rtnl1$^{+}$ OR Rtnl1$^{18}$) ; Ib-GAL4, UAS-myr::GCaMP6s / (TM6B)* | *(Rtnl1$^{+}$ OR Rtnl1$^{18}$) / (CyO::GFP); UAS-tdTom::Sec61β / (TM6B)* | *Rtnl1$^{x}$; UAS-tdTom::Sec61β / Ib-GAL4, UAS-myr::GCaMP6s* |
| 5, S9 | *(Rtnl1$^{+}$ OR Rtnl1$^{18}$) / (CyO,Tb); Is-GAL4, UAS-myr::GCaMP6s / (TM6B)* | *(Rtnl1$^{+}$ OR Rtnl1$^{18}$) / (CyO::GFP); UAS-tdTom::Sec61β/(TM6B)* | *Rtnl1$^{x}$; UAS-tdTom::Sec61β / Is-GAL4, UAS-myr::GCaMP6s* |
| 5E | *(Rtnl1$^{+}$ OR Rtnl1$^{18}$) ; Ib-GAL4, UAS-myr::GCaMP6s / (TM6B)* | *(Rtnl1$^{+}$ OR Rtnl1$^{18}$) / (CyO,Tb); + / (TM6B)* | *Rtnl1$^{x}$; Ib-GAL4, UAS-myr::GCaMP6s / +* |
| 5E | *(Rtnl1$^{+}$ OR Rtnl1$^{18}$) ; Ib-GAL4, UAS-myr::GCaMP6s / (TM6B)* | *(Rtnl1$^{+}$ OR Rtnl1$^{18}$) / (CyO,Tb); UAS-Rtnl1::HA / (TM6B)* | *Rtnl1$^{x}$; Ib-GAL4, UAS-myr::GCaMP6s / UAS-Rtnl1::HA* |
| S9 | *(Rtnl1$^{+}$ OR Rtnl1$^{18}$) / (CyO,Tb); Is-GAL4, UAS-myr::GCaMP6s / (TM6B)* | *(Rtnl1$^{+}$ OR Rtnl1$^{18}$) / (CyO::GFP); UAS-tdTom::Sec61β / (TM6B)* | *Rtnl1$^{+}$ / Rtnl1$^{18}$; UAS-tdTom::Sec61β / Is-GAL4, UAS-myr::GCaMP6s* |
| S9 | *Rtnl1$^{18}$ / (CyO,Tb); Is-GAL4, UAS-myr::GCaMP6s / (TM6B)* | *Rtnl1$^{1}$ / (CyO::GFP); UAS-tdTom::Sec61β / (TM6B)* | *Rtnl1$^{1}$ / Rtnl1$^{18}$; UAS-tdTom::Sec61β / Is-GAL4, UAS-myr::GCaMP6s* |
| S13 | *(Rtnl1$^{+}$ OR Rtnl1$^{18}$) ; Ib-GAL4, UAS-CEPIA3mt::myc / (TM6B)* | *(Rtnl1$^{+}$ OR Rtnl1$^{18}$) ; Ib-GAL4, UAS-CEPIA3mt::myc / (TM6B)* | *Rtnl1$^{x}$; Ib-GAL4, UAS-CEPIA3mt::myc* |
| 8, S13 | *(Rtnl1$^{+}$ OR Rtnl1$^{18}$) ; Ib-GAL4, UASTattB-CEPIA3mt / (TM6B)* | *(Rtnl1$^{+}$ OR Rtnl1$^{18}$) / (CyO::GFP); UAS-tdTom::Sec61β / (TM6B)* | *Rtnl1$^{x}$; UAS-tdTom::Sec61β / Ib-GAL4, UAS-CEPIA3mt* |
| 8, S12 | *(Rtnl1$^{+}$ OR Rtnl1$^{18}$) / (CyO,Tb); Is-GAL4, UAS-CEPIA3mt/ (TM6B)* | *(Rtnl1$^{+}$ OR Rtnl1$^{18}$) / (CyO::GFP); UAS-tdTom::Sec61β/ (TM6B)* | *Rtnl1$^{x}$; UAS-tdTom::Sec61β / Is-GAL4, UAS-CEPIA3mt* |
| S12 | *(Rtnl1$^{+}$ OR Rtnl1$^{18}$) / (CyO,Tb); Is-GAL4, UAS-CEPIA3mt / (TM6B)* | *(Rtnl1$^{+}$ OR Rtnl1$^{18}$ ) / (CyO::GFP); UAS-tdTom::Sec61β/(TM6B)* | *Rtnl1$^{+}$/Rtnl1$^{18}$; UAS-tdTom::Sec61β / Is-GAL4, UAS-CEPIA3mt* |
| S12 | *Rtnl1$^{18}$ / (CyO,Tb); Is-GAL4, UAS-CEPIA3mt / (TM6B)* | *Rtnl1$^{1}$/(CyO::GFP); UAS-tdTom::Sec61β / (TM6B)* | *Rtnl1$^{1}$ / Rtnl1$^{18}$; UAS-tdTom::Sec61β / Is-GAL4, UAS-CEPIA3mt* |
| 8E | *(Rtnl1$^{+}$ OR Rtnl1$^{18}$) ; Ib-GAL4, UAS-CEPIA3mt / (TM6B)* | *(Rtnl1$^{+}$ OR Rtnl1$^{18}$) /(CyO,Tb); +/(TM6B)* | *Rtnl1$^{x}$; Ib-GAL4, UAS-CEPIA3mt::myc / +* |
| 8E | *(Rtnl1$^{+}$ OR Rtnl1$^{18}$) ; Ib-GAL4, UAS-CEPIA3mt / (TM6B)* | *(Rtnl1$^{+}$ OR Rtnl1$^{18}$) / (CyO,Tb); UAS-Rtnl1::HA / (TM6B)* | *Rtnl1$^{x}$; Ib-GAL4, UAS-CEPIA3mt::myc / UAS-Rtnl1::HA* |
