## Supplementary Table 3 for "*Drosophila* SPG12 ortholog, reticulon-like 1, governs presynaptic ER organization and Ca^2+^ dynamics"

**Supplementary Table 3. Reagents used in this work.**

| Reagent | Source/RRID | Notes |
|---|---|---|
| **Antibodies** | | |
| Mouse anti-Dlg 4F3 | DSHB Cat# 4F3 anti-discs large, RRID:AB_528203 (Parnas et al., 2001) | 1:100 dilution; Figure S5A |
| Chicken anti-GFP | Abcam Cat# ab13970, RRID:AB_300798 | 1:400 dilution; Figure S5A |
| Rat anti-tdTomato | ChromoTek Cat# 3f5-100, RRID:AB_2336063 | 1:400 dilution; Figure S5A |
| Rabbit anti-Myc | Cell Signaling Technology Cat# 2272, RRID:AB_10692100 | 1:400 dilution; Figure S15A |
| Goat anti-Mouse Secondary Antibody, Alexa Fluor 647 conjugate | Thermo Fisher Scientific Cat# A-21235, RRID:AB_2535804 | 1:100 dilution; Figure S5A |
| Goat anti-Chicken Secondary Antibody, Alexa Fluor 488 conjugate | Thermo Fisher Scientific Cat# A-11039, RRID:AB_2534096 | 1:100 dilution; Figure S5A |
| Goat anti-Rat Secondary Antibody, Alexa Fluor 488 conjugate | Thermo Fisher Scientific Cat# A-11006, RRID:AB_2534074 | 1:100 dilution; Figure S5A |
| Donkey anti-Rabbit IgG Secondary Antibody, Alexa Fluor 594 | Molecular Probes Cat# A-21207, RRID:AB_141637 | 1:100 dilution |
| **Dissection and $Ca^{2+}$ imaging** | | |
| Vectashield Antifade Mounting Medium | Vector Laboratories cat# H-1000-10 | ----- |
| PBS | Sigma Product number: P4417-100TAB<br>MDL: MFCD00131855 | for PBS 1x: 1P4417-100 tablet in 200 mL $H_2O$ |
| PBT | ----- | PBS + 0.1% triton-X100 |
| Fixation solution | ----- | PBS + 4% formaldehyde |
| BSA serum 1x | Sigma Product Number: A2153<br>CAS Number: 9048-46-8<br>MDL number: MFCD00130384 | 1 mL PBT + 0.61 mM BSA |
| Normal Goat Serum Blocking Solution | Vector Laboratories Cat# S-1000, RRID:AB_2336615 | PBT + 5% NGS |
| Schneider's *Drosophila* medium | Sigma MDL:<br>MFCD00242089 cat# S0146 | ----- |
| Prefixation HL3 ($Ca^{2+}$ free) | ----- | In mM: 70 NaCl, 5 KCl, 20 $MgCl_2$, 10 $NaHCO_3$, 115 sucrose, 5 Trehalose, 5 HEPES, 0.5 EGTA; pH 7.20±0.05 |
| HL3 (low $Mg^{2+}$) | (Oliva et al., 2020) | In mM: 7.0 L-Glutamic acid, 1.0 $Ca^{2+}$, 0.5 $Mg^{2+}$, 6.8 NaCl, 0.5 KCl, 9.5 $NaHCO_3$, 11.5 Sucrose, 5.5 Trehalose, 0.5 HEPES; pH 7.20±0.05 |
| HL3 (low $Mg^{2+}$; low L-Glutamic acid) | ----- | In mM: 1.0 L-Glutamic acid, 1.0 $Ca^{2+}$, 0.5 $Mg^{2+}$, 6.8 NaCl, 0.5 KCl, 9.5 $NaHCO_3$, 11.5 Sucrose, 5.5 Trehalose, 0.5 HEPES; pH 7.20±0.05 |

| | | |
|---|---|---|
| L-Glutamic acid | Sigma-Aldrich PubChem ID: 24895069 | in HL3 |
| KCl | Sigma-Aldrich PubChem ID: 329820207 | in HL3 |
| $CaCl_2$ solution | Sigma-Aldrich PubChem ID: 57647902 | in HL3 |
| $MgCl_2$ solution | Sigma-Aldrich PubChem ID: 329759772 | in HL3 |
| NaCl | Sigma-Aldrich PubChem ID: 329768085 | in HL3 |
| $NaHCO_3$ | Sigma-Aldrich PubChem ID: 329824613 | in HL3 |
| Sucrose | Sigma-Aldrich PubChem ID: 24899835 | in HL3 |
| Trehalose | Sigma-Aldrich PubChem ID: 57654693 | in HL3 |
| HEPES | Sigma-Aldrich PubChem ID: 24895572 | in HL3 |
| EGTA (Ethylene glycol-bis(2-aminoethylether)-*N,N,N′,N′*-tetraacetic acid) | Sigma-Aldrich PubChem ID: 24894530 | in $Ca^{2+}$-free HL3 |
| | **Molecular biology** | |
| Agarose | Sigma Aldrich cat# A9539, PubChem ID: 24891469 | ----- |
| Bbs1-HF | New England Biolabs cat# R0539 | ----- |
| BD Bacto Agar | Thermo Fisher Scientific cat# BD214010 | ----- |
| BD Difco LB Broth, Miller | Thermo Fisher Scientific cat# BD244620 | ----- |
| DreamTaq Green PCR Master Mix | Thermo Fisher Scientific cat# K1081 | ----- |
| GeneRuler 1 kb Plus DNA Ladder | Thermo Fisher Scientific cat# SM1333 | ----- |
| Gibson Assembly Master Mix | New England Biolabs cat# E2611L/S | ----- |
| High efficiency Transformation Protocol | New England Biolabs cat# C2987H | ----- |
| Phusion High-Fidelity Polymerase | New England Biolabs cat# M0531 | ----- |
| plasmid *pCFD5* | Addgene cat# 73914; RRID:Addgene_73914 | ----- |
| Q5 High-Fidelity DNA Polymerase | New England Biolabs cat# M0492 | ----- |
| QIAprep Spin Miniprep Kit | Qiagen cat# 27104 | ----- |
| Surveyor Mutation Detection Kit | Integrated DNA Technologies cat# 706020 | ----- |
| TBE | Thermo Fisher Scientific cat# 15581044 | ----- |
| TRIzol reagent | Invitrogen cat# 15596026 | ----- |

| | | |
|---|---|---|
| | **Electron Microscopy** | |
| Paraformaldehyde | Merck cat# 8.18715.1000 | ----- |
| Glutaraldehyde, 25 % solution | TAAB Laboratories cat# G017 | ----- |
| Uranyl acetate | AnalaR cat# 1426810 | ----- |
| Maleic acid | Merck cat# 8.00380.0500 | ----- |
| Sodium cacodylate trihydrate | Sigma-Aldrich cat# 20840 | ----- |
| Calcium chloride dihydrate | Sigma-Aldrich cat# C7902 | ----- |
| Thiocarbohydrazide | Sigma-Aldrich cat# 223220 | ----- |

| | | |
|---|---|---|
| Osmium tetroxide | TAAB Laboratories cat# O001 | ----- |
| Lead nitrate | Sigma-Aldrich cat# 10099-74-8 | ----- |
| L-Aspartic acid | Sigma-Aldrich cat# A8949 | ----- |
| Ethanol | Sigma-Aldrich cat# 32221 | ----- |
| Acetone | Sigma-Aldrich cat# 32201 | ----- |
| Acetonitrile | Macron Fine chemicals cat# 2856-25 | ----- |
| Potassium ferricyanide | AnalaR cat# 0153090 | ----- |
| Quetol 651 | TAAB Laboratories cat# Q001 | ----- |
| MNA, methyl nadic anhydride | TAAB Laboratories cat# M010 | ----- |
| NSA, nonenyl succinic anhydride | TAAB Laboratories cat# N007 | ----- |
| BDMA, benzyldimethylamine | TAAB Laboratories cat# B036 | ----- |
| Kapton tape | LabTech cat# RMCATUM308-15 | ----- |
| 4-inch silicon wafers | LabTech cat# 10-008140 | ----- |
