## Supplemental Figure S1 for "*Drosophila* SPG12 ortholog, reticulon-like 1, governs presynaptic ER organization and Ca^2+^ dynamics"

**A**

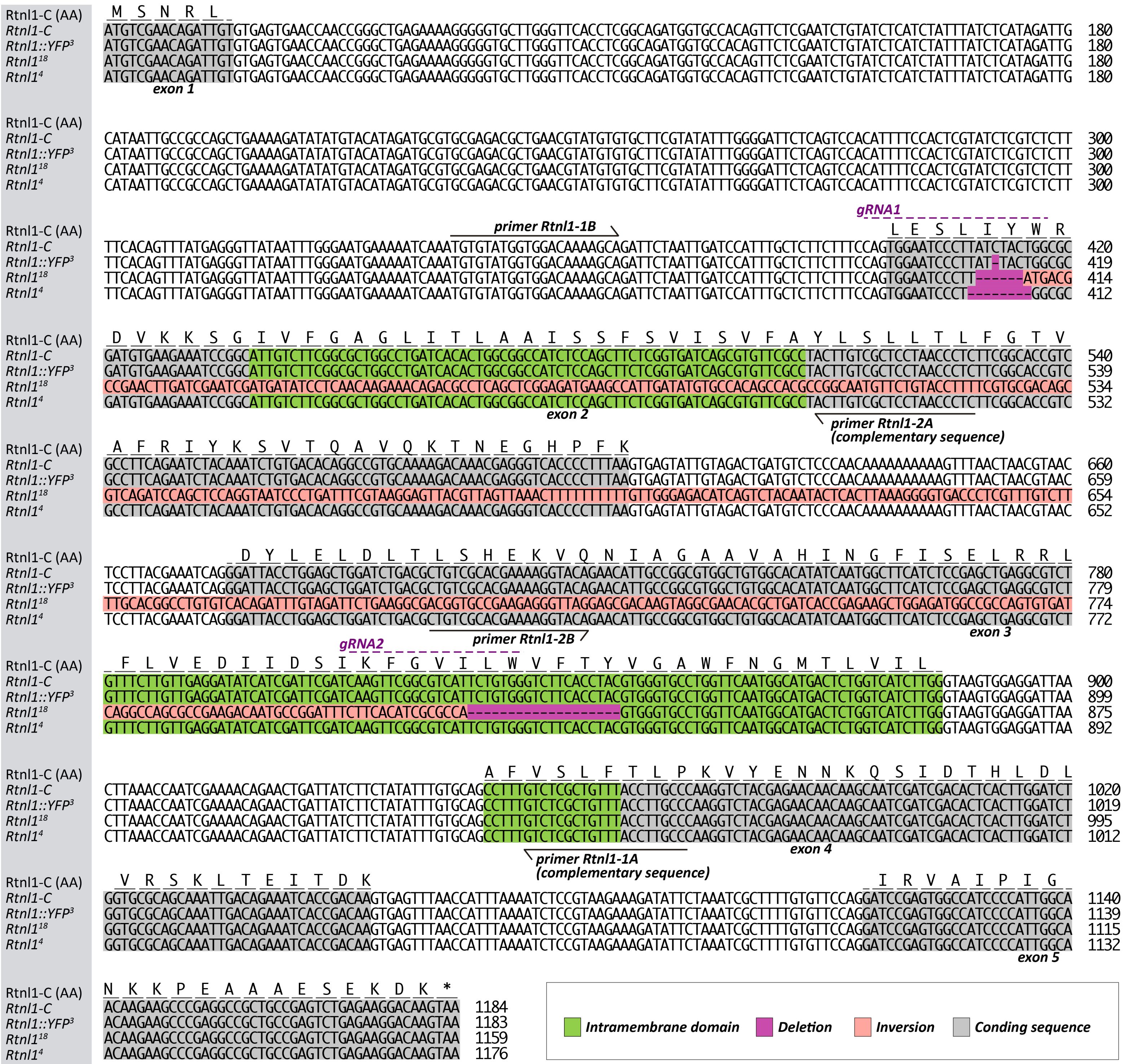


**B**

**Rtnl1 (Isoform C)**

MSNRLLESLIYWRDVKKSGIVFGAGLITLAAISSFSVISVFAYLSLLTLFGTVAFRIYKSVTQAVQKTNEGHPFKDYLELDLTLSHEKVQNIAGVAVAHINGFISELRRLFLV
EDIIDSIKFGVILWVFTYVGAWFNGMTLVILAFVSLFTLPKVYENNKQSIDTHLDLVRSKLTEITDKIRVAIPIGNKKPEAAAESEKDK*

**Rtnl1::YFP³ (Isoform C)**

MSNRLLESLITGAM*RNPALSSALA*SHWRPSPASR*SACSPTCRS*PSSAPSPSESTNL*HRPCKRQTRVTPLRITWSWI*RCRTKRYRTLPAWLWHISMASSPS*GV
CFLLRISSIRSSSASFCGSSPTWVPGSMA*LWSSWPLSRCLPCPRSTRTTSNRSTLTWIWCAAN*QKSPTRSEWPSPLATRSPRPLPSLRRTS

**Rtnl1¹⁸ (Isoform C)**

MSNRLLESLMTPNLIESMISSTRNRRLSSEMKPLICATATPAMFCTFSCDSVRSSSR*SLIS*GVTLVKLFFLLGDISLQYSLKGVTLVCLLHGLCHRFVDSEGDGAEEG*
ERQVGEHADHREAGDGRQCDQASAEDNAGFLHIAPVGAWFNGMTLVILAFVSLFTLPKVYENNKQSIDTHLDLVRSKLTEITDKIRVAIPIGNKKPEAAAESEKDK*

**Rtnl1⁴ (Isoform C)**

MSNRLLESLARCEEIRHCLRRWPDHTGGHLQLLGDQRVRLLVAPNPLRHRRLQNLQICDTGRAKDKRGSPL*GLPGAGSDAVARKGTEHCRRGCGTYQWLHLRAEASVSC*G
YHRFDQVRRHSVGLHLRGCLVQWHDSGHLGLCLAVYLAQGLREQQAIDRHSLGSGAQQIDRNHRQDPSGHPHWQQEARGRCRV*EGQV
