## Supplementary figures and images for "*Drosophila* SPG12 ortholog, reticulon-like 1, governs presynaptic ER organization and Ca^2+^ dynamics"

### Supplemental Figure S2

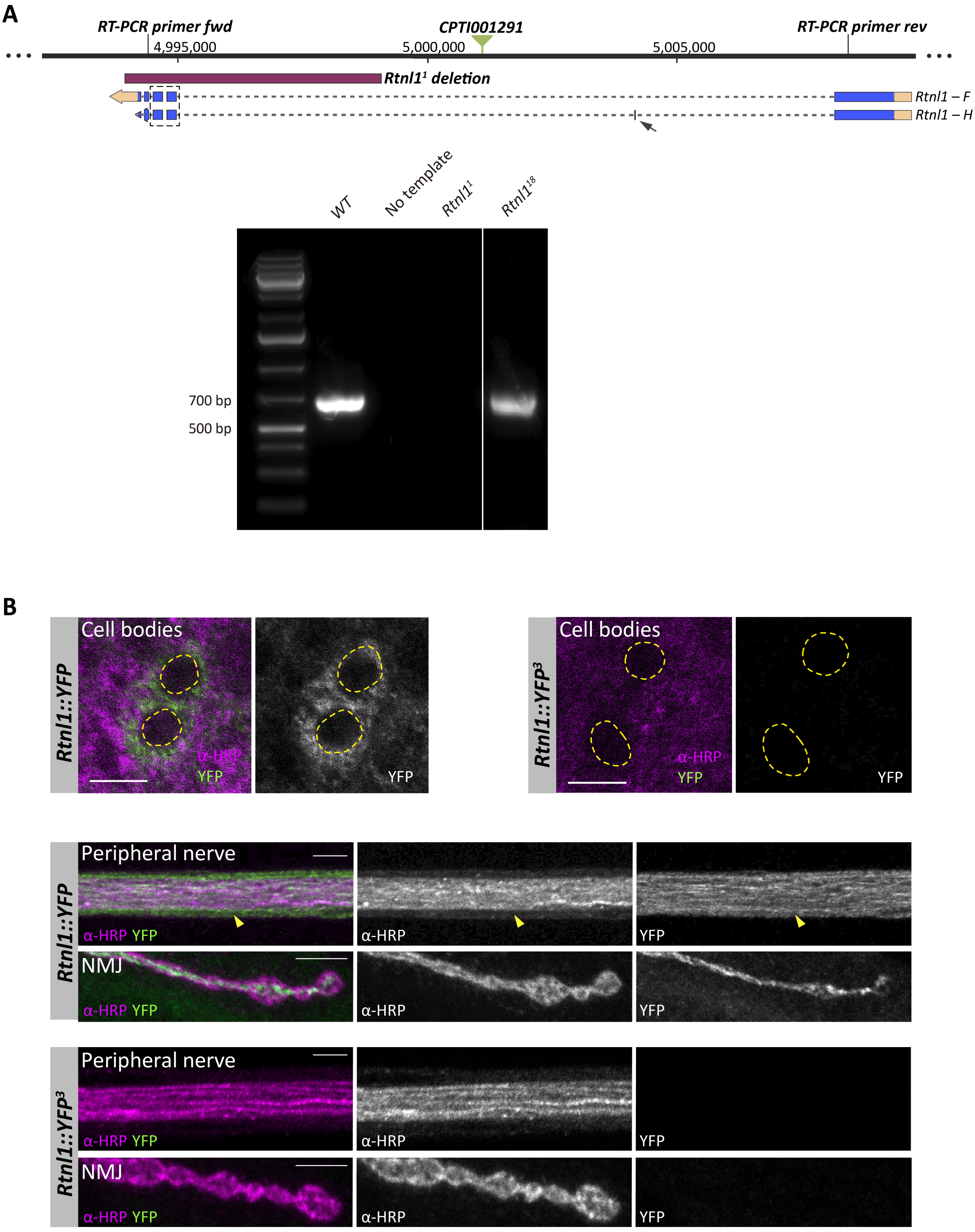

### Supplemental Figure S3

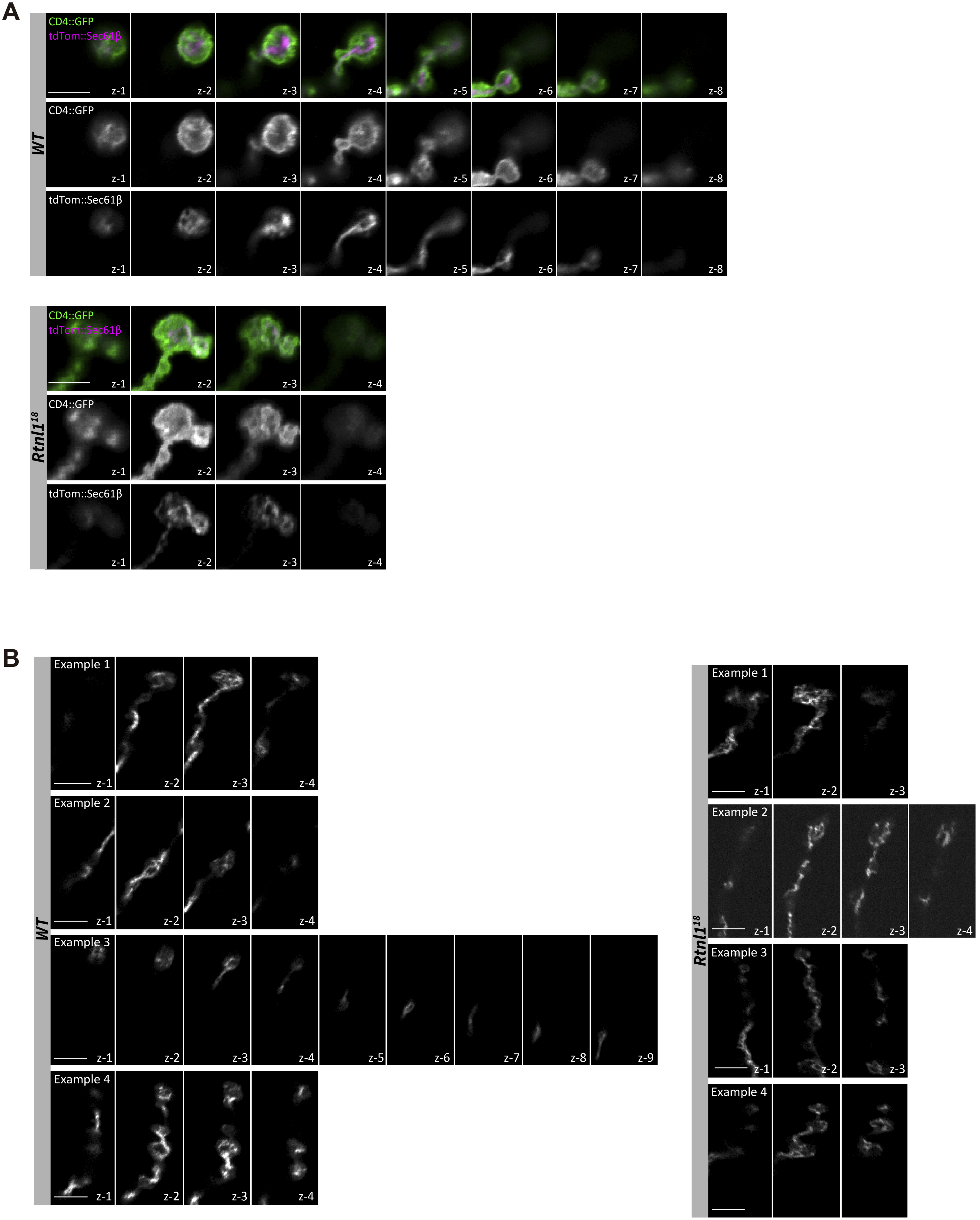

### Supplemental Figure S4

#### Figure S4

**A**

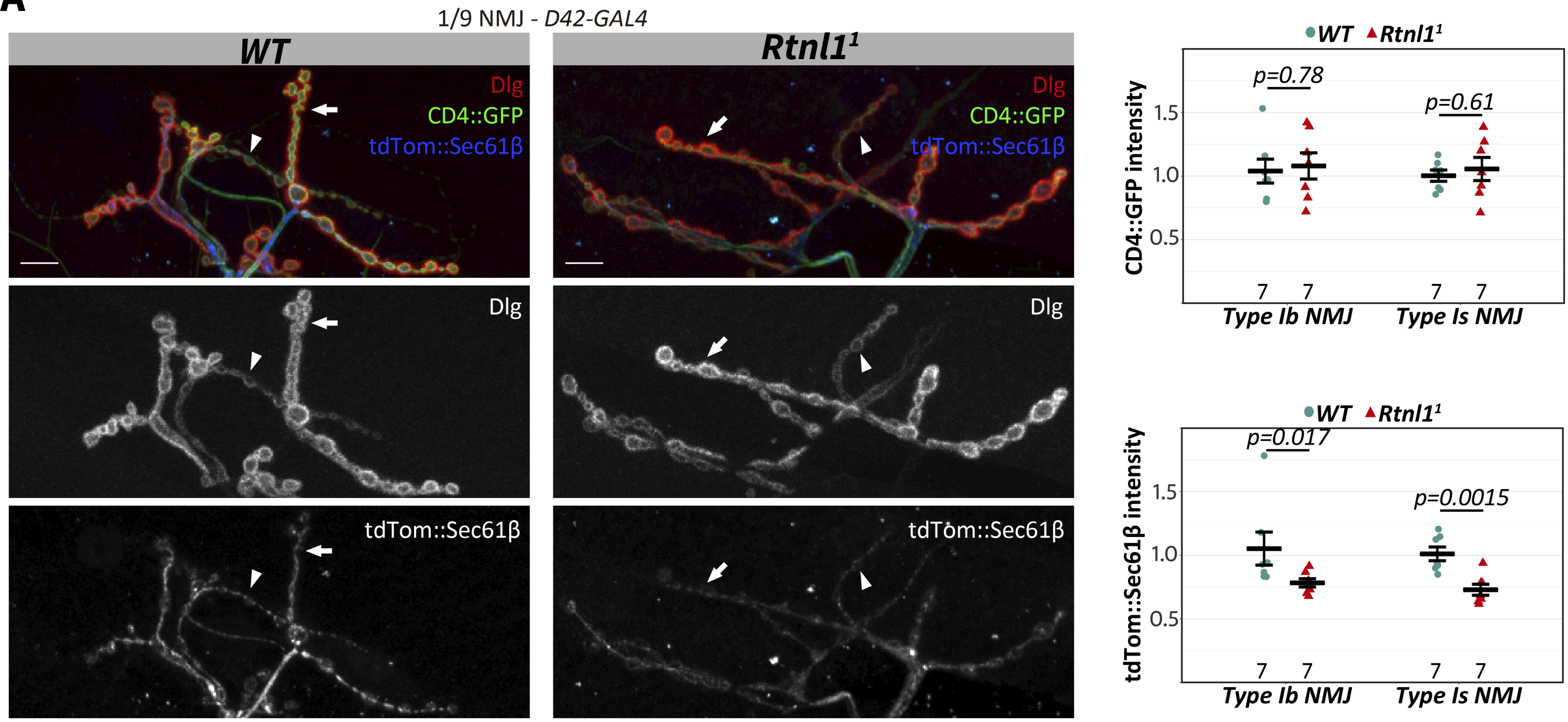


**B**

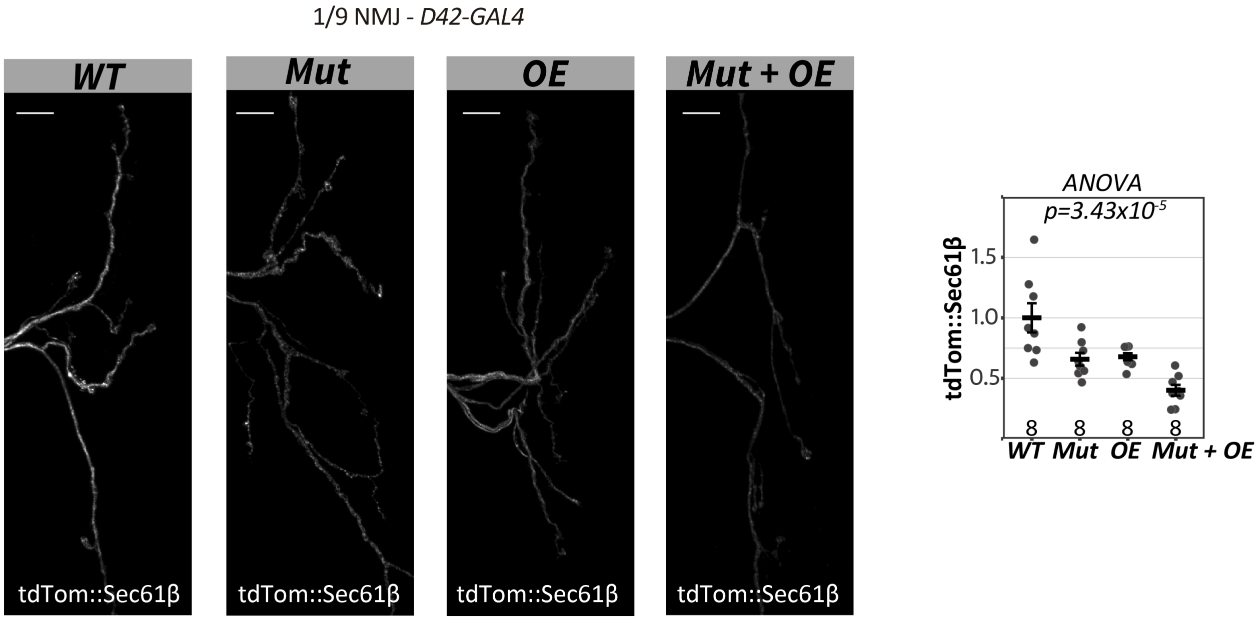


*Tukey Test*

| Genotype | p-value |
|---|---|
| Mut-WT | 0.0104 |
| Overexp-WT | 0.0173 |
| MutOE-WT | $12x10^{-5}$ |
| OE-Mut | 0.997 |
| MutOE-Mut | 0.0743 |
| MutOE-OE | 0.0475 |

### Supplemental Figure S5

#### Figure S5

**A**

Type Ib Motor Neuron

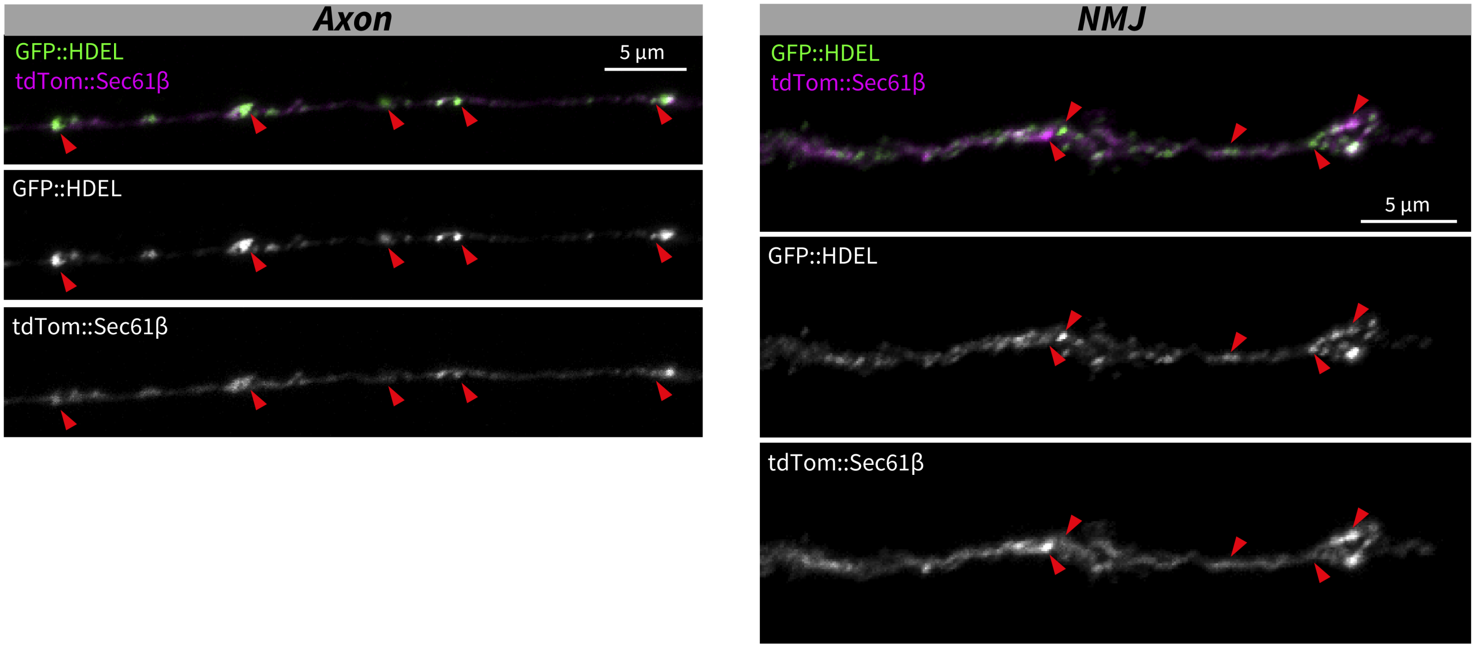


**B**

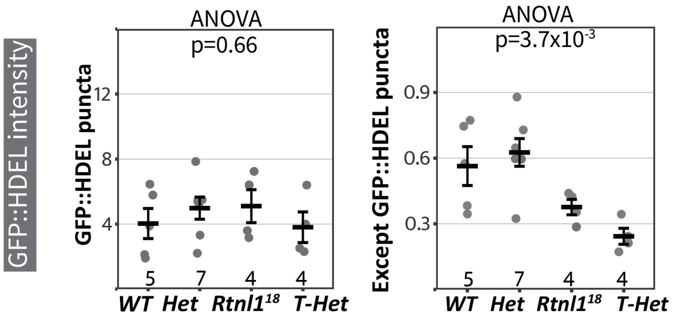


Tukey Test

| Genotype | p-value |
|---|---|
| WT-Het | 0.89 |
| Rtnl1[18]-THet | 0.596 |

**C**

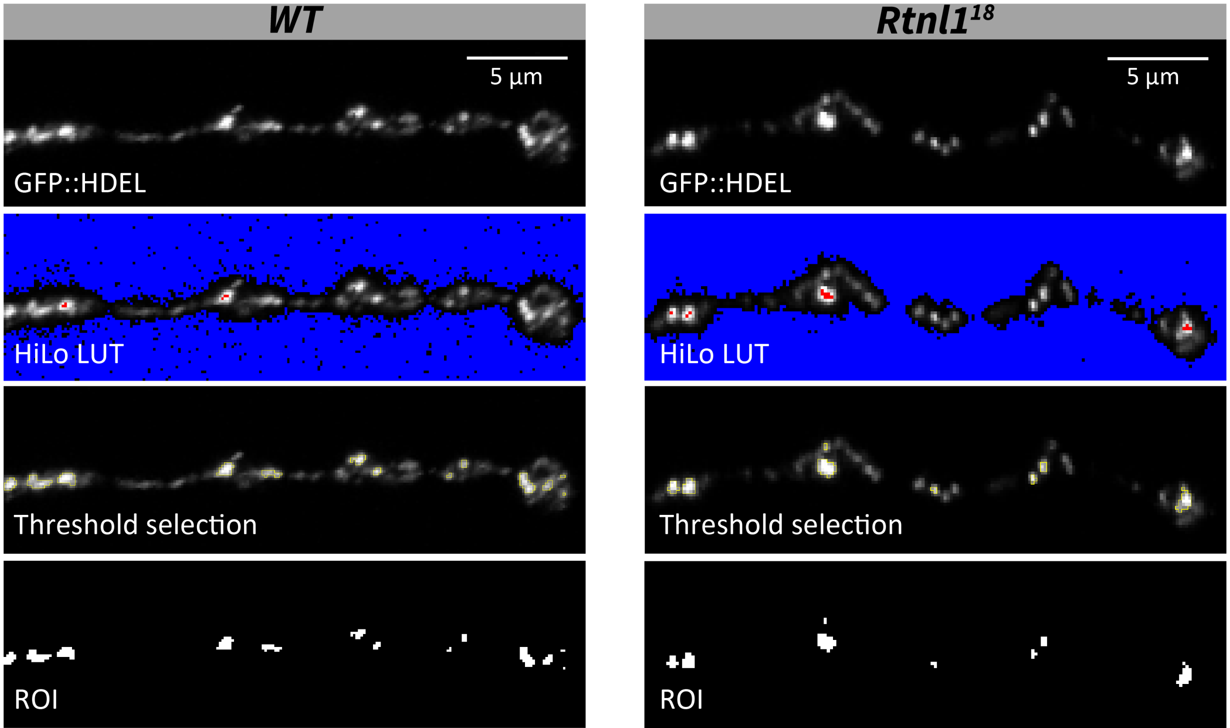

### Supplemental Figure S6

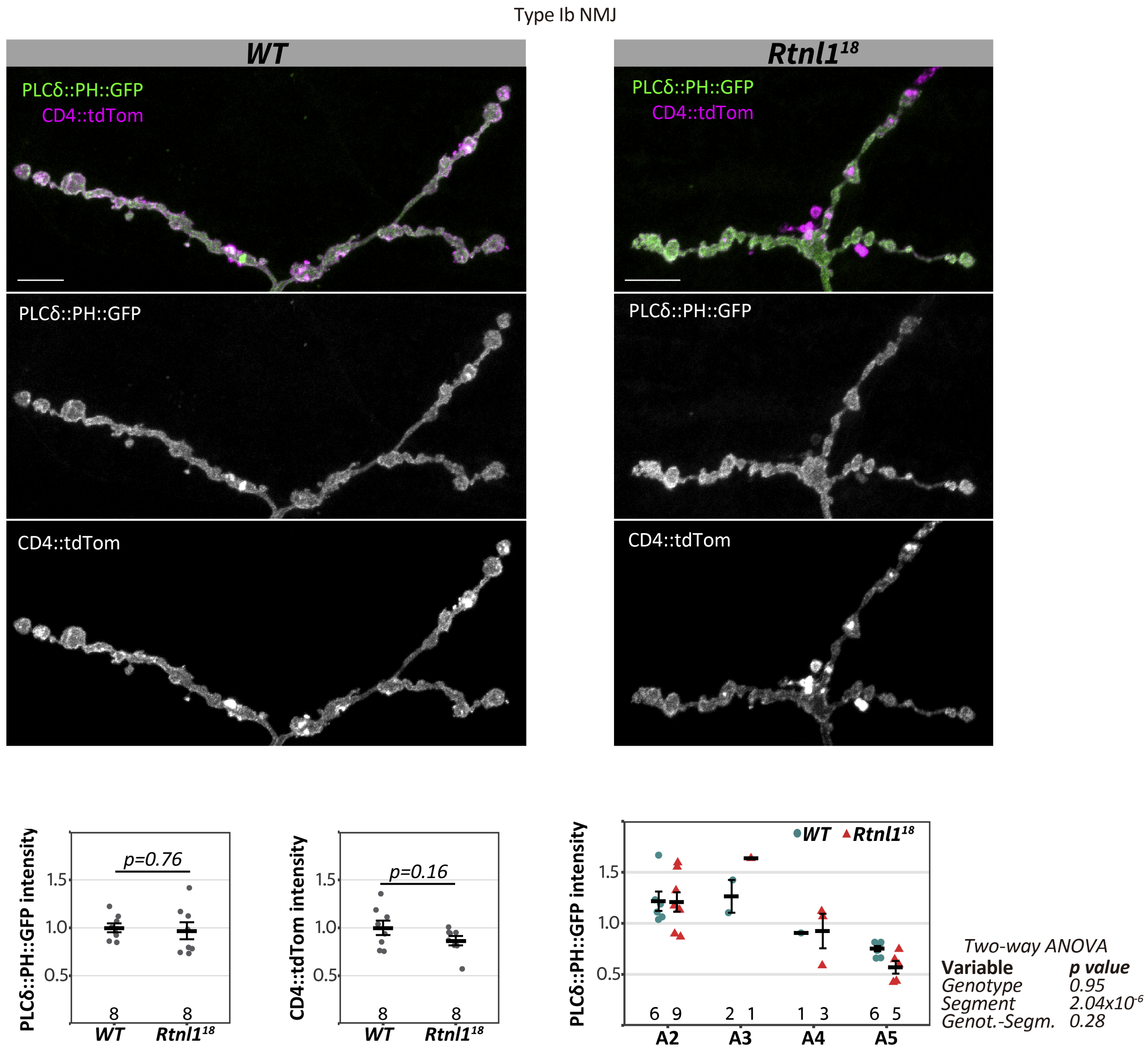

### Supplemental Figure S7

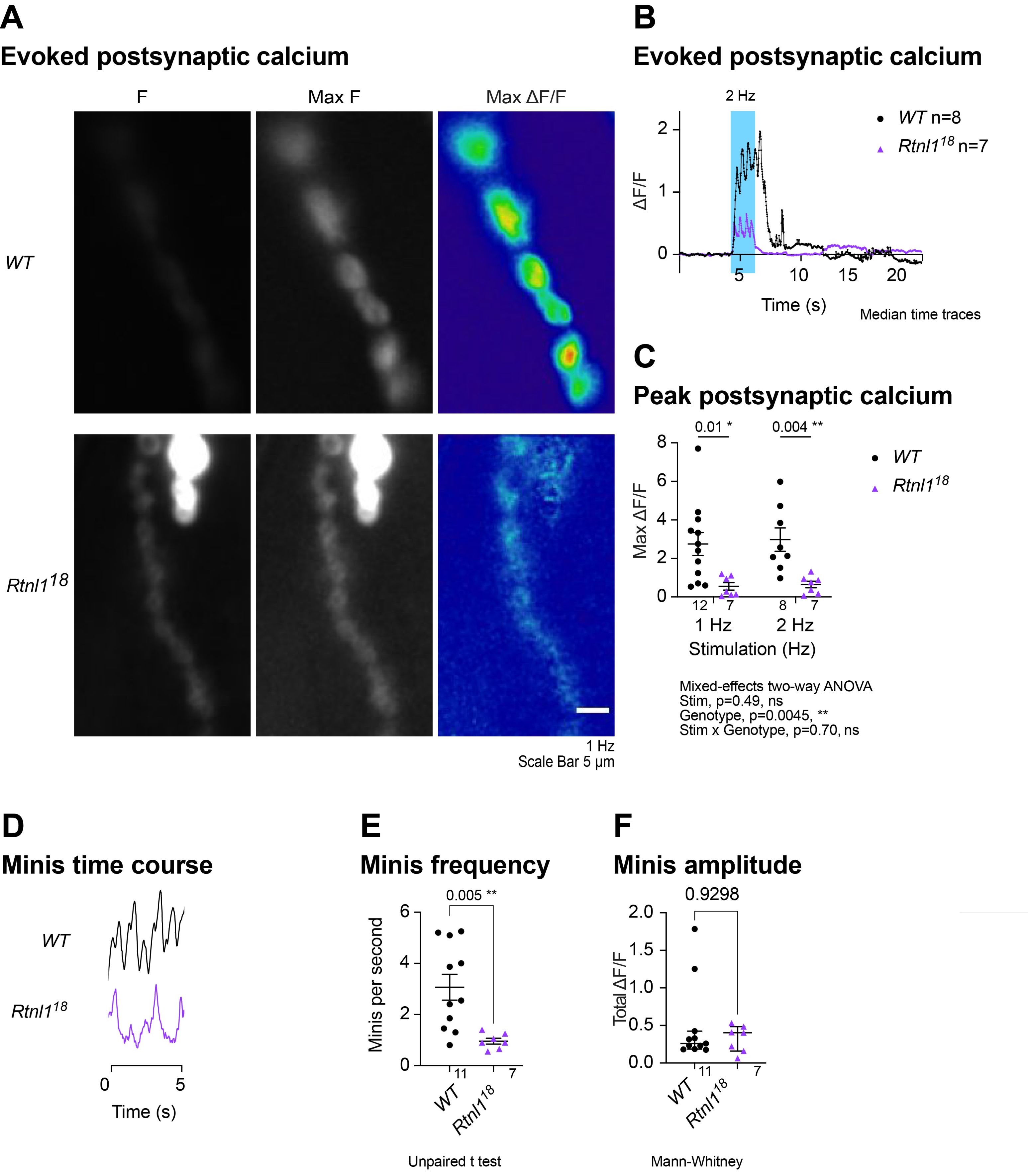

### Supplemental Figure S8

#### Figure S8

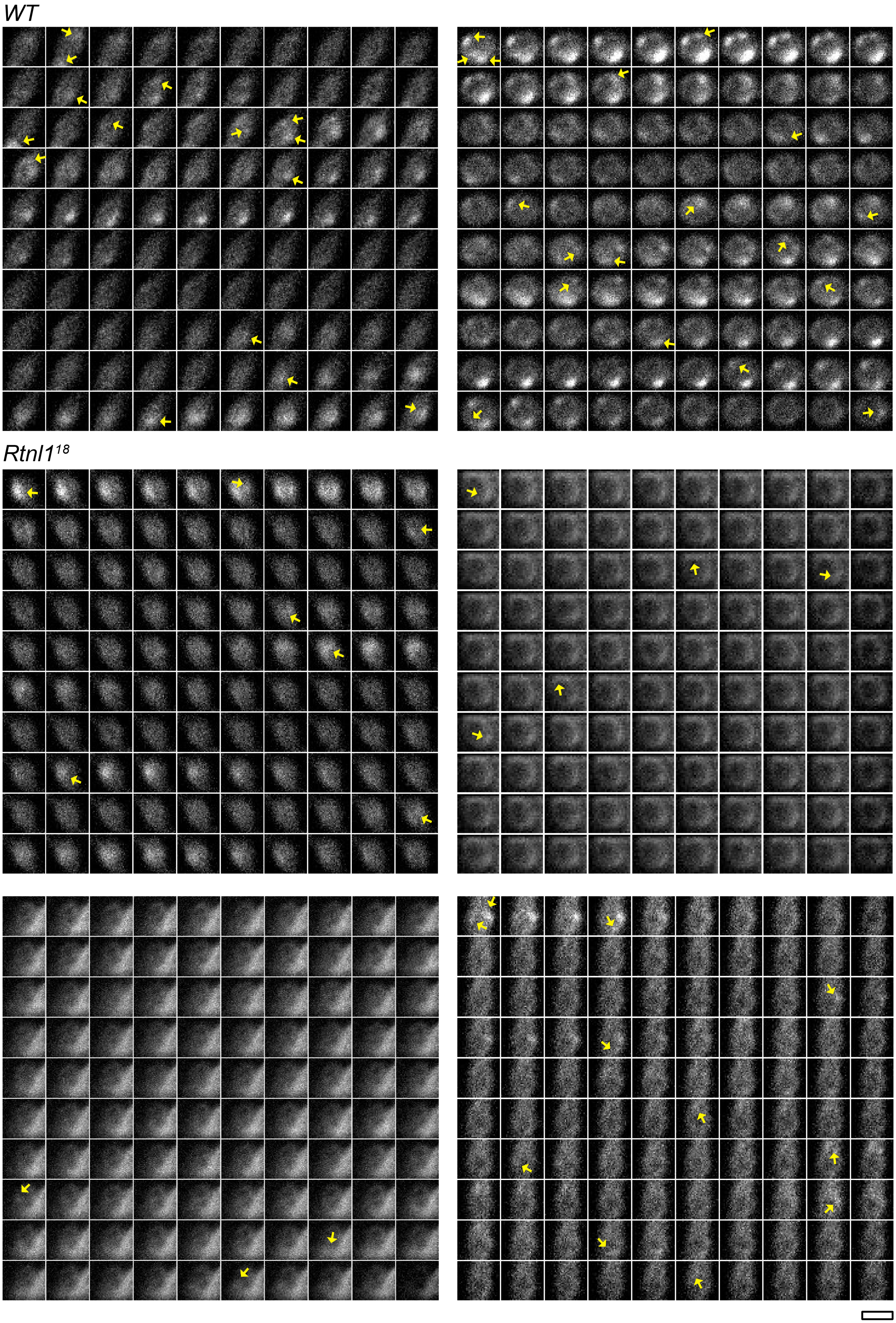


100 frames in 8 seconds, 12.5 frames/second

Scale bar 5 μm

### Supplemental Figure S12

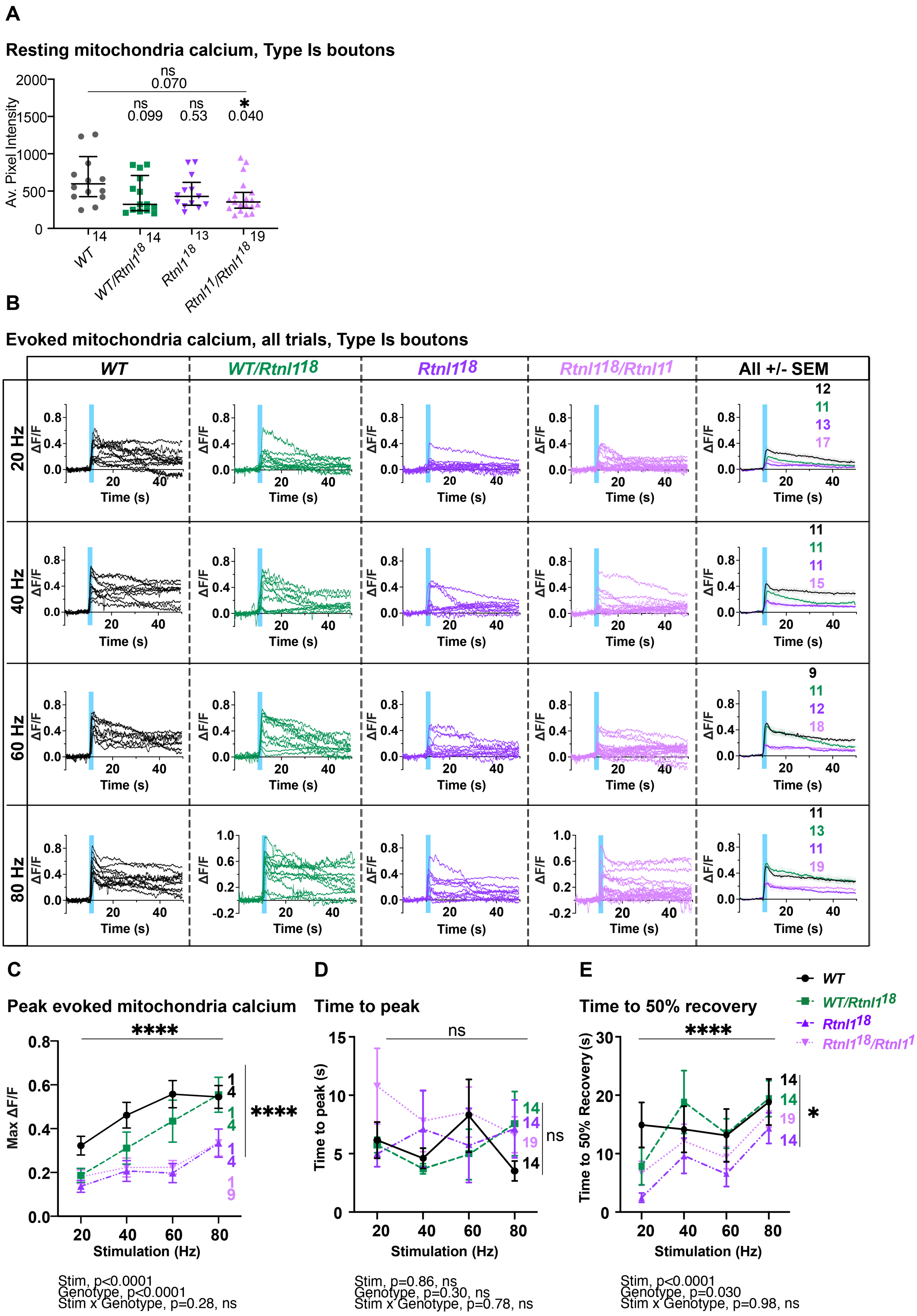

### Supplemental Figure S13

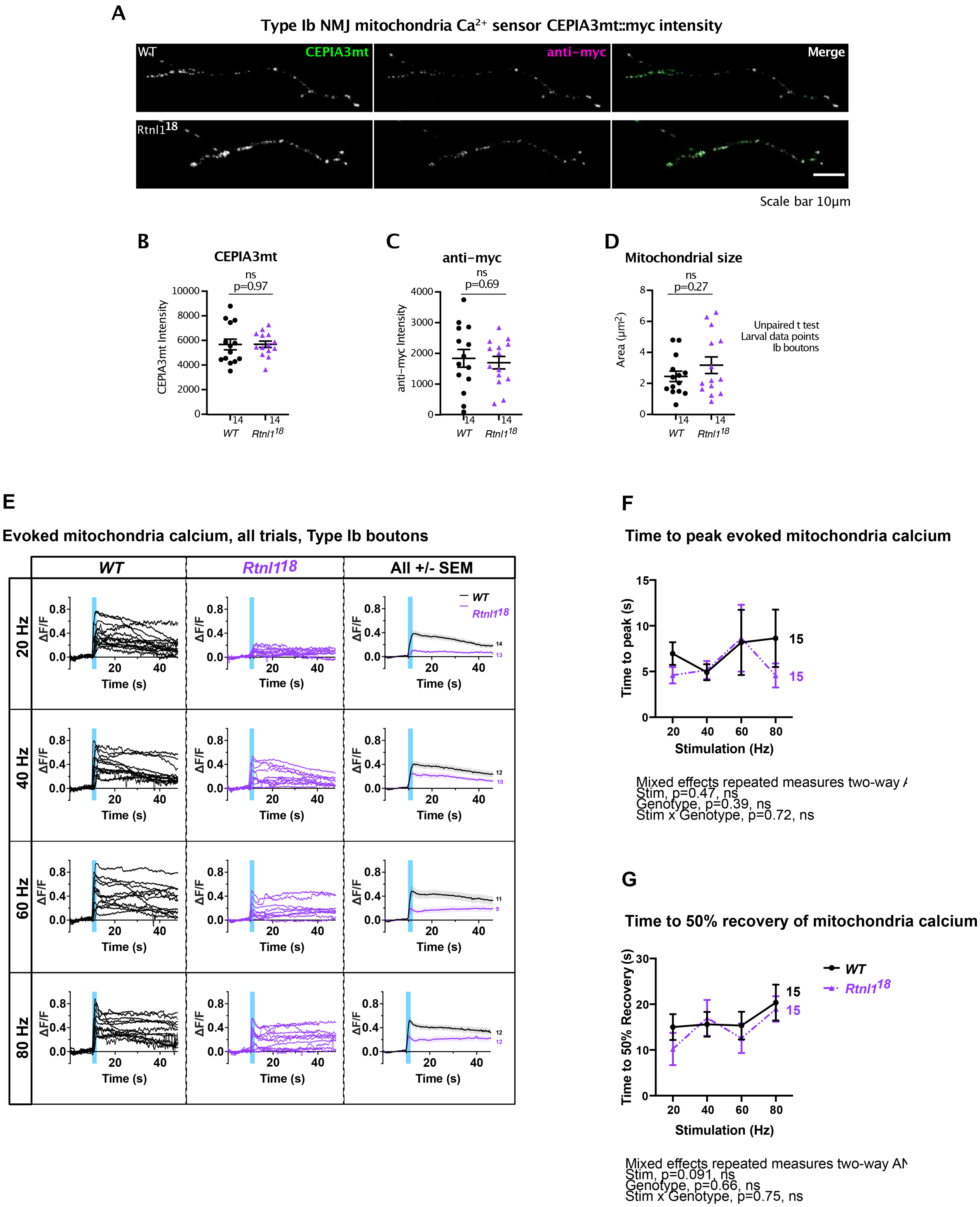
