## Supplemental Figure S9 for "*Drosophila* SPG12 ortholog, reticulon-like 1, governs presynaptic ER organization and Ca^2+^ dynamics"

**A**
**Resting cytosolic calcium, Type Is boutons**

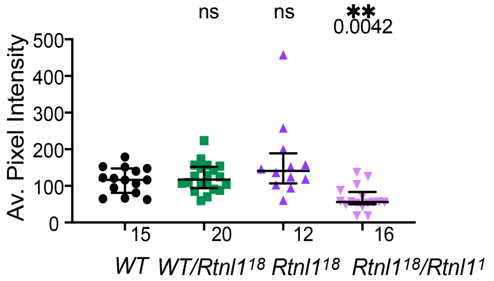


**B**
**Evoked cytosolic calcium, Type Is boutons**

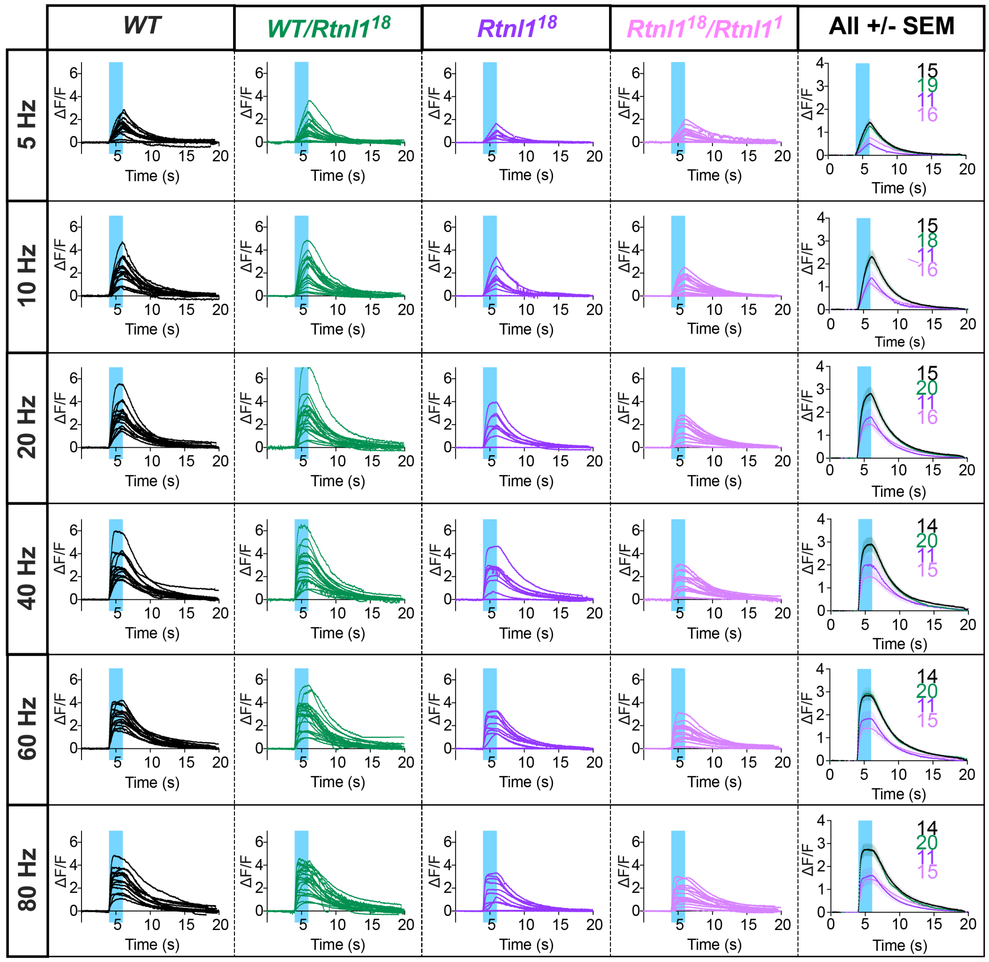


**C** **Maximum influx** **D** **Evoked time to peak influx** **E** **Time to 50% influx recovery** **F** **Time to full influx recovery**

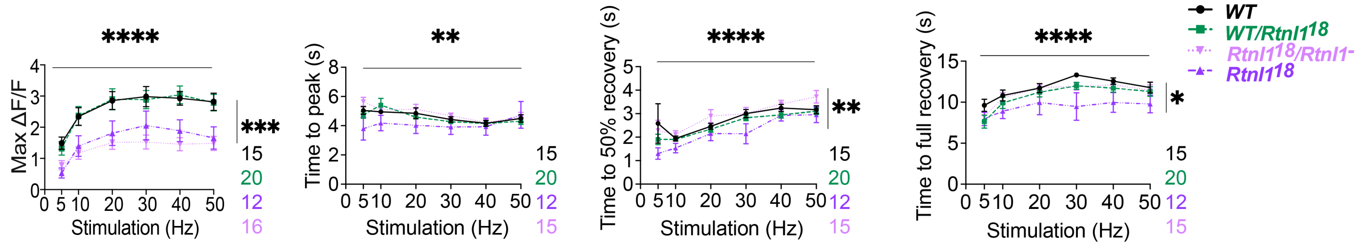


Stim, p<0.0001
Genotype, p=0.0001
Stim x Genotype, p=0.068, ns

**No *Rtnl1*^18**
Stim, p<0.0001
Genotype, p=0.0002 ***
Stim x Genotype, p=0.017 *

Stim, p=0.0061
Genotype, p=0.058, ns
Stim x Genotype, p=0.27, ns

**No *Rtnl1*^18**
Stim, p<0.0001
Genotype, p=0.27, ns
Stim x Genotype, p=0.50, ns

Stim, p<0.0001
Genotype, p=0.0021
Stim x Genotype, p=0.94, ns

**No *Rtnl1*^18**
Stim, p<0.0001
Genotype, p=0.056, ns
Stim x Genotype, p=0.93, ns

Stim, p<0.0001
Genotype, p=0.012
Stim x Genotype, p=0.70, ns

**No *Rtnl1*^18**
Stim, p<0.0001
Genotype, p=0.20, ns
Stim x Genotype, p=0.84, ns
