## Supplemental Figure S10 for "*Drosophila* SPG12 ortholog, reticulon-like 1, governs presynaptic ER organization and Ca^2+^ dynamics"

## A

### Evoked calcium influx, all trials, Type Ib boutons

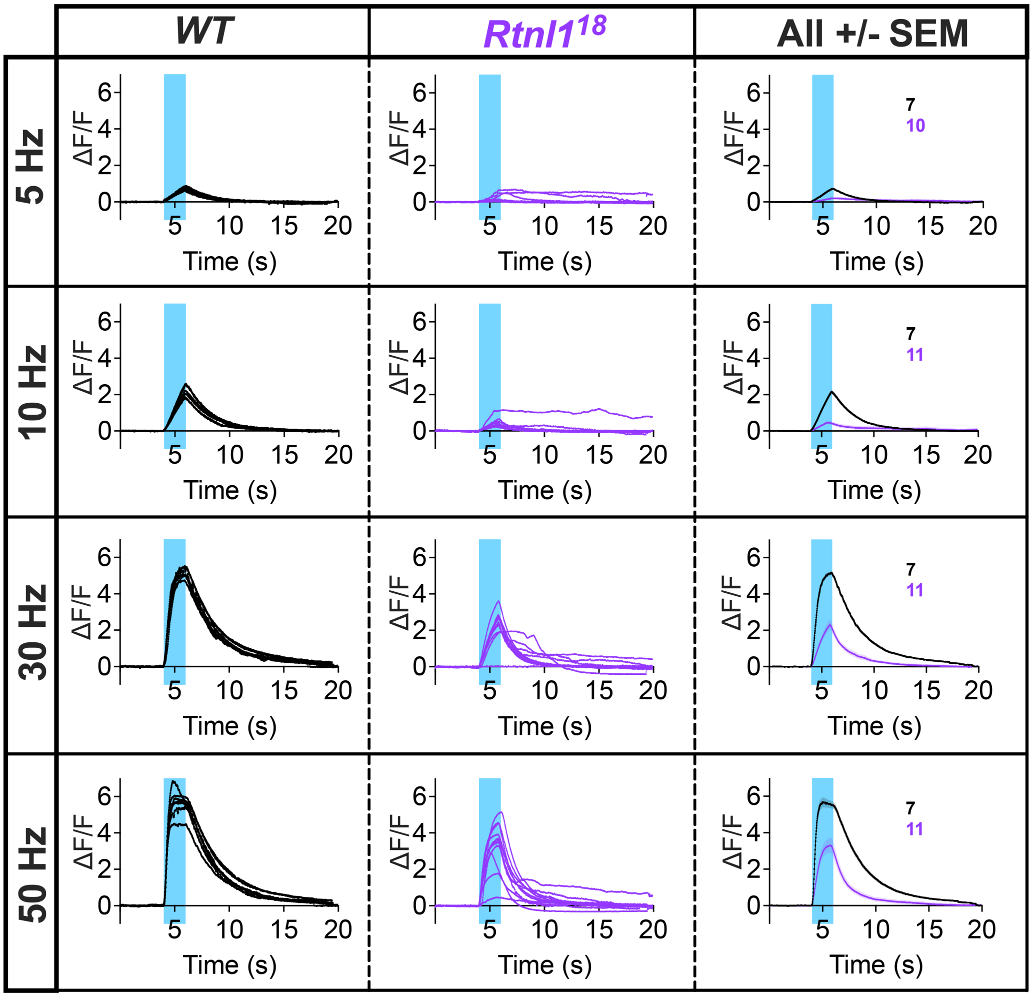


## B

### Time to peak

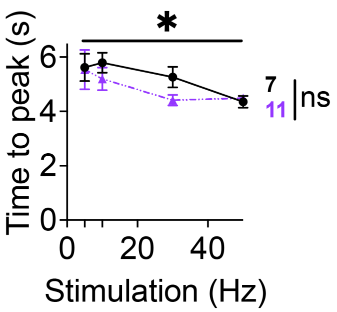


Stim, p=0.043
Genotype, p=0.25, ns
Stim x Genotype, p=0.63, ns

## C

### Time to 50% recovery

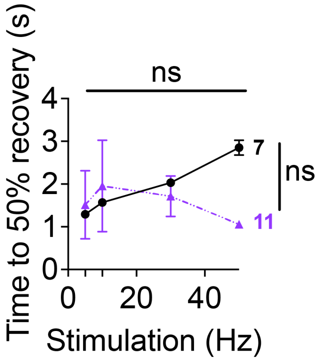


Stim, p=0.068, ns
Genotype, p=0.60, ns
Stim x Genotype, p=0.096, ns

## D

### Time to full recovery

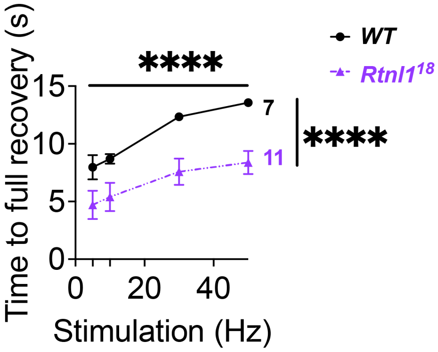


Stim, p<0.0001
Genotype, p<0.0001
Stim x Genotype, p=0.72, ns
