## Supplemental Figure S11 for "*Drosophila* SPG12 ortholog, reticulon-like 1, governs presynaptic ER organization and Ca^2+^ dynamics"

A

Evoked ER calcium response, all trials, Type Is boutons

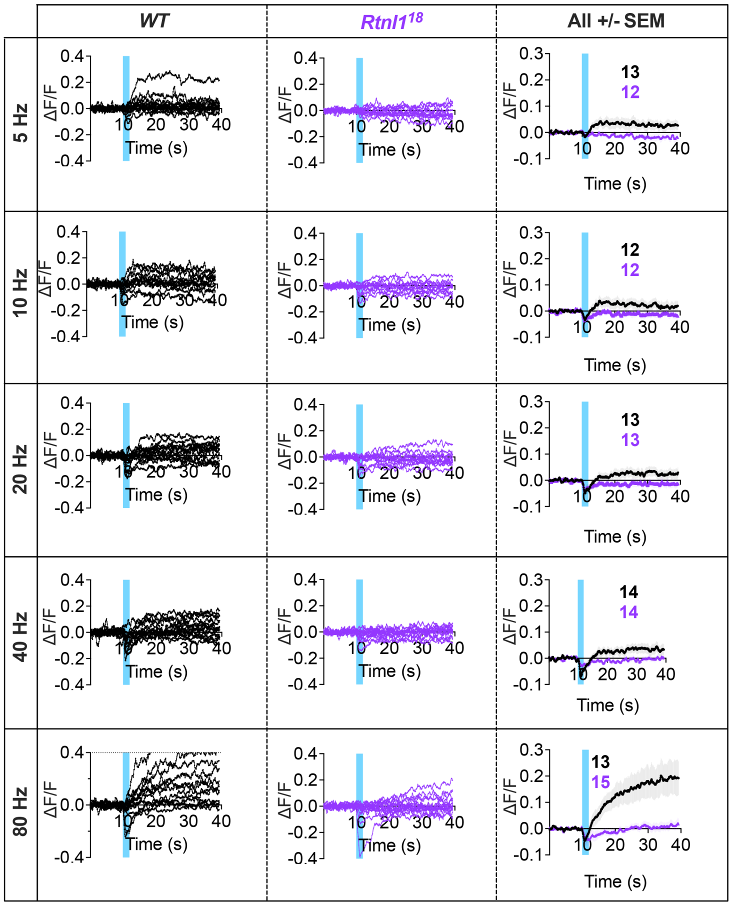


B

Evoked ER calcium response, all trials, Type Ib boutons

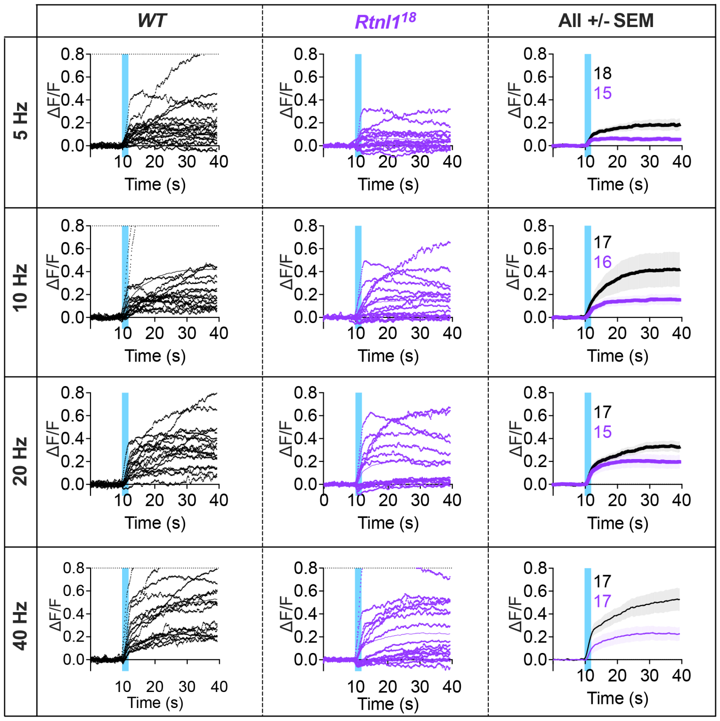
