## Supplementary Figures and Video Legends for "*Drosophila* SPG12 ortholog, reticulon-like 1, governs presynaptic ER organization and Ca^2+^ dynamics"

### SUPPLEMENTARY MATERIAL

**Supplementary Figure S1. CRISPR-derived lesions in *Rtnl1* and their effects on the *Rtnl1* coding region.** (**A**) Manual alignment of the *Rtnl1* transcript *C* for *WT* (*Rtnl1-C*), and *Rtnl1::YFP*$^{3}$, *Rtnl1*$^{18}$ and *Rtnl1*$^{4}$ mutant CRISPR alleles, showing the amino acids (AA) encoded by the *WT* (*Rtnl1-C*) allele, the location of the *gRNAs* used to generate the mutant alleles, and the primers used to genotype them (see Methods for details). Since the sequences encoding intramembrane domains are shared by all *Rtnl1* isoforms, the shortest one, isoform C, was chosen for convenience. Sequences read from 5' to 3'**.** The *Rtnl1::YFP*$^{3}$ allele was generated from *Rtnl1::YFP*$^{CPTI00291}$, and has a 1-bp deletion/frameshift at the position of *gRNA1* upstream of the first intramembrane domain, and lacks detectable YFP expression (**Supp. Fig. S2**); we did not use it further in this work, but it shows that frameshifts around this position can lead to loss of protein expression. The effects of *Rtnl1*$^{18}$ on the protein-coding sequence are described in the main paper. *Rtnl1*$^{4}$ has an 8-bp deletion/frameshift at the position of *gRNA1*, upstream of the first intramembrane domain. (**B**) Predicted Rtnl1-C protein sequences for the *Rtnl1* alleles shown in **A**. Intramembrane domains are decorated in green and stop codons shown with a red asterisk.

**Supplementary Figure S2**. **Effects of *Rtnl1* mutant CRISPR alleles on *Rtnl1* expression.** (**A**) RT-PCR strategy to test presence of *Rtnl1* transcripts (top). For the *Rtnl1* isoforms shown, blue boxes indicate exons, broken lines indicate introns, light orange boxes indicate 5' and 3' UTRs, and arrows indicate the direction of transcription. A green triangle shows the position of the *Rtnl1::YFP* exon trap insertion (*CPTI001291*). Genomic coordinates are based on Release 6.26 of the *Drosophila melanogaster* genome (http://flybase.org). Using the indicated primers, the expected amplicon from genomic DNA is around 14 kb. The expected amplicon for *Rtnl1* transcript *F* is 636 bp, and for transcript *H* is 672 bp, due to an extra small exon (arrow). The box surrounded by a broken line shows the region containing *Rtnl1*$^{18}$ lesions. Agarose gel electrophoresis (1%) of reverse-transcribed cDNA (bottom) shows that *Rtnl1*$^{18}$ does not disrupt *Rtnl1* mRNA transcription. For both *WT* and *Rtnl1*$^{18}$ alleles, only one amplicon is seen, which is slightly smaller in *Rtnl1*$^{18}$, due to the small deletions totaling 25bp within the amplified region (see **Fig. 1A** and **Supp. Fig. 1A** for details). (**B**) Confocal sections of neuronal cell bodies, peripheral nerves and NMJs (muscle 1) show that *Rtnl1::YFP*$^{3}$ mutation is enough to abolish Rtnl1::YFP expression. In cell bodies, nuclei are distinguished by the absence of α-HRP signal (dotted line regions). On the peripheral nerve, glia can be distinguished from neuronal axons due to the low levels of α-HRP signal (arrowhead). All larvae are also expressing *Ib-GAL4*, but this is not driving any reporter expression. Scale bars, 5 μm.

**Supplementary Figure S3. Rtnl1 loss does not noticeably disrupt presynaptic ER network organization.** (**A**) Confocal sections (z steps, 1 μm) of the projections presented in the magnified areas of **Fig. 2A**, showing the distribution of the ER marker tdTom::Sec61β in Type Ib muscle 1 NMJ of *WT* and *Rtnl1*$^{18}$ larvae. CD4::GFP labels the plasma membrane. Genotypes are *Ib-GAL4, UAS-CD4::tdGFP / UAS-tdTom::Sec61β*, in either a *WT or Rtnl1*$^{18}$ background. (**B**) Confocal sections (z steps, 1 μm) of the projections presented in the magnified areas of **Fig. 2E**, showing the distribution of the ER marker Sturkopf::GFP in Type Ib muscle 1 NMJ of *WT* and *Rtnl1*$^{18}$ larvae. Genotypes are *Ib-GAL4, UAS-Sturkopf::GFP / +*, in either a *WT* or *Rtnl1*$^{18}$ background. Scale bars, 5 μm.

**Supplementary Figure S4**. **Effects of *Rtnl1* mutant alleles on presynaptic tdTom::Sec61β levels.** (**A**) Representative examples of confocal projections and quantifications of *WT* and *Rtnl1*$^{1}$ larvae showing the distribution of the ER marker tdTom::Sec61β in Type I NMJs (muscles 1/9 NMJ). Samples were immunostained for tdTom, GFP, and Dlg to distinguish between Type Ib (arrows) and Type Is (arrowheads) NMJs. Genotypes: *D42-GAL4, UAS-tdTom::Sec61β, UAS-CD4::tdGFP /* +, in either *WT* or *Rtnl1*$^{1}$ background. (**B**) Representative examples of confocal projections and quantification of *WT*, *Mut*, *OE*, and *Mut* + *OE* larvae showing the distribution of the ER marker tdTom::Sec61β in Type I NMJs (muscles 1/9 NMJ). Genotypes:

*WT - Rtnl1*$^{+}$ *; D42-GAL4, UAS-tdTom::Sec61β / +*,

*Mut - Rtnl1*$^{18}$ */ Rtnl1*$^{1}$ *; D42-GAL4, UAS-tdTom::Sec61β / +*

*OE - Rtnl1*$^{+}$ ; *D42-GAL4, UAS-Rtnl1::GFP / UAS-tdTom::Sec61β*

*Mut + OE - Rtnl1*$^{18}$ */ Rtnl1*$^{1}$ ; *D42-GAL4, UAS-Rtnl1::GFP / UAS-tdTom::Sec61β*.

For **A** and **B**, Plots show individual larval datapoints and mean ± SEM; y-axis indicates arbitrary units (au) after normalization to control (*WT*); sample size (number of larvae) is indicated within the plots for each genotype. For each larva, several NMJs between A2-A6 segments were analysed, and the mean value is shown as a larval datapoint. Student's t-tests were performed for pairwise comparisons, except for tdTom::Sec61*β* intensity in Type Ib NMJ comparison (**A**), where a Mann-Whitney U test was performed. Scale bars, 10 μm.

**Supplementary Figure S5**. **GFP::HDEL distribution.** (**A**) Representative examples of confocal projections of *WT* larvae showing the distribution of the ER markers tdTom::Sec61β and GFP::HDEL in a Type Ib motor neuron. Genotype: *Rtnl1$^{+}$ / UAS-tdTom::Sec61β ; Ib-GAL4 / UAS-GFP::HDEL*. (**B**) Quantification of the ER marker GFP::HDEL in Type I NMJs (muscles 1/9 NMJ) of *WT*, *Het*, *Rtnl1$^{18}$*, and *T-Het* larvae. Genotypes**:**

*WT* - *Rtnl1$^{+}$ ; Ib-GAL4, UAS-CD4::tdTom / UAS-GFP::HDEL*

*Het* - *Rtnl1$^{+}$* / *Rtnl1$^{18}$* ; *Ib-GAL4, UAS-CD4::tdTom / UAS-GFP::HDEL*

*Rtnl1$^{18}$* - *Rtnl1$^{18}$* ; *Ib-GAL4, UAS-CD4::tdTom / UAS-GFP::HDEL*

*T-Het* - *Rtnl1$^{18}$* / *Rtnl1$^{1}$* ; *Ib-GAL4, UAS-CD4::tdTom / UAS-GFP::HDEL*

Plots show individual larval datapoints and mean ± SEM; y-axis indicates arbitrary units (au) after normalization to control (*WT*); intensity values are relative to CD4::tdTom signal; sample size (number of larvae) is indicated within the plots for each genotype. For each larva, several NMJs between A2-A6 segments were analysed, and the mean value is shown as a larval datapoint. (**C**) Representative examples showing the selection of GFP::HDEL puncta in confocal projections of Type Ib muscle 1 NMJ (magnified areas from **Fig. 3C)**. HiLo lookup table (LUT) shows saturated pixels in red and pixels with no detectable signal in blue. Intermodes thresholding of GFP::HDEL intensity was used to select only those regions with high GFP::HDEL levels (region of interest, ROI). Genotypes are *Ib-GAL4, UAS-CD4::tdTom / UAS-BiP::sfGFP::HDEL*, in either a *Rtnl1$^{+}$* (*WT*) *or Rtnl1$^{18}$* background.

**Supplementary Figure S6**. **Rtnl1 loss does not affect presynaptic PI(4,5)$P_2$ levels.** Representative examples of confocal projections and quantifications of *WT and Rtnl1$^{18}$* larvae showing the distribution of the PI(4,5)$P_2$ marker PLC$\delta$::PH::GFP in Type Ib muscle 1 NMJ. Scale bars, 10 µm. Plots show individual larval datapoints and mean ± SEM; y-axis indicates arbitrary units (au) after normalization to control (*WT*); Sample size (larvae) is indicated within the plot for each genotype. For each larva, several NMJs between A2-A6 segments were analyzed, and the mean larval value is shown as a datapoint. Student's t-tests were performed for pairwise comparisons. Genotypes are *Ib-GAL4, UAS-CD4::tdTom / UAS-PLC$\delta$::PH::GFP*, in either a *Rtnl1$^{+}$* (*WT*) or *Rtnl1$^{18}$* background.

**Supplementary Figure S7. Rtnl1 loss impacts synaptic transmission.** (**A**) GCaMP fluorescence at rest (F), maximum fluorescence (Max F) in response to 1 Hz stimulation, and maximum change in fluorescence (Max ΔF) in examples of *WT* and *Rtnl1$^{18}$* at muscle 1, Type Is postsynaptic terminals. (**B**) Impact of Rtnl1 loss-of-function on peak evoked postsynaptic $Ca^{2+}$. Plot shows the median responses to a burst of 2 Hz stimulation of larvae from each genotype. (**C**) Impact of Rtnl1 loss-of-function on peak evoked postsynaptic $Ca^{2+}$ responses. Plot shows individual larval datapoints and mean ± SEM; Datapoints represent the largest ΔF/F reached after either a 1 Hz or 2 Hz stimulation during the recording. Comparisons were analyzed with a mixed-effects two-way ANOVA. (**D**) Impact of Rtnl1 loss-of-function on postsynaptic resting miniature responses (minis). Plots show representative time traces (close to the mean stimulation frequency) over 5 seconds from the distal bouton during the recording collected. (**E**) Impact of Rtnl1 loss-of-function on minis frequency. Plot shows individual larval datapoints and mean ± SEM. Frequency datapoints represent the number of minis per second over a five-second recording. Pairwise comparison was performed using Student's t-test. (**F**) Rtnl1 loss-of-function did not affect minis amplitude. Plot shows individual larval datapoints and median ± interquartile ranges. For each larva in **E** and **F**, minis from a 20-second recording from the distal bouton of one NMJ between segments A4-A6. Amplitude datapoints represent the largest ΔF/F over the recording. Pairwise comparison was performed using Mann-Whitney U-test (maximum mini amplitude). (**B**-**F**) Sample size (larvae) is indicated within each plot for each genotype. Each recording was from one muscle 1, Type Is NMJ between segments A4-A6. Genotypes are *Is-GAL4, mhc-SynapGCaMP6f / UAS-tdTom::Sec61β*, in either a *WT* or *Rtnl1*$^{18}$ background.

**Supplementary Figure S8. Rtnl1 loss decreases miniature neurotransmission frequency.** (Extended data from **Supp. Fig. S7 D**-**E**). Representative examples (close to the median frequency) of miniature events in 100 frames over 8 seconds (12.5 frames/second). Panels show time-lapse GCaMP fluorescence at rest in distal boutons of muscle 1, Type Is postsynaptic terminals, in *WT* (two boutons) and *Rtnl1*$^{18}$ (four boutons). Arrows indicate miniature events counted. Genotypes are *Is-GAL4, mhc-SynapGCaMP6f/ UAS-tdTom::Sec61β*, in either a *WT* or *Rtnl1*$^{18}$ background.

**Supplementary Figure S9. Rtnl1 loss decreases cytosolic $Ca^{2+}$ handling in Type Is boutons.** (Extended data from **Fig 5**). (**A**) Loss of Rtnl1 does not affect resting myrGCaMP6s fluorescence. *Rtnl1*$^{18}$/*Rtnl1*$^{1}$ flies show a decrease in myrGCaMP6s fluorescence, but this effect is not replicated in a homozygous *Rtnl1*$^{18}$ background, indicating that loss of Rtnl1 does not affect resting cytosolic $Ca^{2+}$. The plot shows individual larval datapoints and median ± interquartile ranges; sample size (larvae) is indicated within the plot for each genotype. The pairwise comparison performed was a Kruskal-Wallis test. (**B**) Impact of Rtnl1 loss-of-function on peak evoked cytosolic $Ca^{2+}$

responses. Plots show all single time traces, as well as mean ± SEM time traces in every genotype for 6 stimulation frequencies tested. Sample size is indicated within the plot for each genotype. (**C**) Rtnl1 loss of function decreases peak evoked cytosolic $Ca^{2+}$. Datapoints represent mean ± SEM of the largest ΔF/F reached after stimulation during the recording, shown in **B**. (**D**) Rtnl1 loss did not affect time to peak cytosolic $Ca^{2+}$. Datapoints represent the time between stimulation and peak ΔF/F. Comparisons were analyzed as in **C**. (**E-F**) Rtnl1 loss did not affect time recovery from evoked cytosolic $Ca^{2+}$ influx. Although the mixed-effects two-way ANOVA records a significant genotypic effect in both time to 50% recovery (**E**) and 100% recovery (**F**), two-way ANOVAs without the $Rtnl1^{18}$ group erase this significance, indicating that $Rtnl^{18}/Rtnl^{1}$ transheterozygotes are not significantly different from controls in these assays and that the effect is likely due to the homozygosity of the chromosome carrying $Rtnl1^{18}$, and not to the loss of Rtnl1. Datapoints represent the time between peak ΔF/F and 50% or 100% recovery. Comparisons were analyzed as in **C**. In **C-F**, plots show mean ± SEM of every genotype for each stimulation frequency tested; sample size (larvae) is indicated within the plot for each genotype. For each larva, responses from a 20 second recording from one muscle 1 NMJ between segments A4-A6 were analysed. Comparisons were analyzed with a mixed-effects two-way ANOVA. (**A-F**) Genotypes are *Is-GAL4, UAS-myrGCaMP6s / UAS-tdTom::Sec61β*, in either a *WT,* or $Rtnl1^{18}$, $Rtnl1^{18}/Rtnl1^{1}$, or *WT /* $Rtnl1^{18}$ background.

**Supplementary Figure S10. Rtnl1 loss decreases cytosolic $Ca^{2+}$ handling in Type Ib boutons.** (Extended data from **Fig 5**). (**A**) Impact of Rtnl1 loss of function on peak evoked cytosolic $Ca^{2+}$. Plots show all single time traces, and mean ± SEM of time traces in both genotypes for the 4 stimulation frequencies tested. Sample size (larvae) is indicated within the plot for each genotype. (**B**) Rtnl1 loss of function does not affect time to peak cytosolic $Ca^{2+}$, or (**C**) time to 50% recovery. (**D**) Rtnl1 loss of function decreases time to 100% recovery in every stimulation frequency tested. In **B-D** plots show mean ± SEM of every genotype for each stimulation frequency tested; sample size (larvae) is indicated within the plot for each genotype. For each larva, we analyzed responses from a 20-second recording from one muscle 1 NMJ between segments A4-A6. Datapoints represent the time to peak ΔF/F, and 50% or 100% recovery. Comparisons were analyzed with a mixed-effects two-way ANOVA. Genotypes are *Ib-GAL4, UAS-myrGCaMP6s / UAS-tdTom::Sec61β*, in either a *WT* or $Rtnl1^{18}$ background.

**Supplementary Figure S11. Rtnl1 loss decreases ER $Ca^{2+}$ handling in Type Is and Ib boutons.** (Extended data from **Fig 6 and 7**). (**A**) Impact of Rtnl1 loss-of-function on evoked ER $Ca^{2+}$ in Is boutons. Plots show all single time traces, and mean ± SEM time traces in both genotypes for the 5 stimulation frequencies tested. Sample size (larvae) is indicated within the plot for each genotype. Genotypes are *Is-GAL4, UAS-ER-GCaMP6-210/UAS-tdTom::Sec61β*, in either a *WT,* or *$Rtnl1^{18}$* background. (**B**) Impact of Rtnl1 loss of function on evoked ER $Ca^{2+}$ in Ib boutons, plotted as in **A**. Genotypes are *Ib-GAL4, UAS-ER-GCaMP6-210 / UAS-tdTom::Sec61β*, in either a *WT* or *$Rtnl1^{18}$* background.

**Supplementary Figure S12. Rtnl1 loss decreases mitochondrial $Ca^{2+}$ handling in Type Is boutons.** (Extended data from **Fig 8**). (**A**) Loss of Rtnl1 does not affect resting CEPIA3mt fluorescence. Although *$Rtnl1^{18}$/$Rtnl1^{1}$* flies show a marginally significant decrease in CEPIA3mt fluorescence, this effect is not found in a homozygous *$Rtnl1^{18}$* background, indicating that loss of Rtnl1 does not affect resting mitochondrial $Ca^{2+}$. Plot shows individual larval datapoints and median ± interquartile ranges; sample size (larvae) is indicated within the plot for each genotype. Comparisons were performed using Kruskal-Wallis tests. (**B**) Impact of Rtnl1 loss-of-function on peak evoked mitochondrial $Ca^{2+}$. Plots show all single time traces, and mean ± SEM time traces in every genotype for 4 stimulation frequencies tested. Sample size (larvae) is indicated within the plot for each genotype. (**C**) Impact of Rtnl1 loss-of-function on peak evoked mitochondrial $Ca^{2+}$. Plots show mean ± SEM of every genotype for each stimulation frequency tested; sample size (larvae) is indicated within the plot for each genotype. For each larva, responses from a 50-second recording from one muscle 1 NMJ between A4-A6 segments were analyzed. Comparisons were analyzed with a mixed-effects two-way ANOVA. (**D**) Rtnl1 loss did not affect time to peak mitochondria $Ca^{2+}$. Data analyzed as in **C**. (**E**) Impact of Rtnl1 loss on time to 50% recovery of mitochondria $Ca^{2+}$. Rtnl1 loss in both *$Rtnl1^{18}$* and *$Rtnl1^{18}$ / $Rtnl1^{1}$* backgrounds slightly decreases time to 50% recovery when compared to *WT* and *WT / $Rtnl1^{18}$* backgrounds, indicating that mitochondria in Rtnl1 mutants lose their $Ca^{2+}$ slightly faster than in *WT*. Data analyzed as in **C**. In **D-E,** datapoints represent the time to peak ΔF/F, and 50% or 100% recovery. Genotypes are *Is-GAL4, UAS-CEPIA3mt / UAS-tdTom::Sec61β*, in either a *WT,* or *$Rtnl1^{18}$*, *$Rtnl1^{18}$ / $Rtnl1^{1}$*, or *WT/$Rtnl1^{18}$* background.

**Supplementary Figure S13. Effect of Rtnl1 loss on mitochondrial $Ca^{2+}$ handling in Type Ib boutons.** (Extended data from **Fig 8**). (**A**) Panels show mitochondrial CEPIA3mt fluorescence and anti-myc signal of CEPIA3mt::myc in typical examples of

*WT* and *Rtnl1$^{18}$* muscle 1, Type Ib postsynaptic terminals. (**B-D**) Rtnl1 loss-of-function does not impact CEPIA3mt fluorescence intensity (**B**), anti-myc signal intensity (**C**) of CEPIA3mt::myc, or mitochondrial size (**D**). Plots shows individual larval datapoints and mean ± SEM; sample size (larvae) is indicated within the plot for each genotype. For each larva, all mitochondria from several muscle 1 NMJs between A2-A6 segments were analysed, and each mean larval value is shown as a datapoint. Pairwise comparisons were performed using Student's t-tests. Genotypes are *Ib-GAL4, CEPIA3mt::myc*, in either a *WT* or *Rtnl1$^{18}$* background. (**E**) Impact of Rtnl1 loss-of-function on peak evoked mitochondria $Ca^{2+}$. Plots show all single time traces, and mean ± SEM time traces in both genotypes for the 4 stimulation frequencies tested. Sample size (larvae) is indicated within the plot for each genotype. (**F**) Rtnl1 loss of function does not affect time to peak mitochondria $Ca^{2+}$, or (**G**) time to 50% recovery of mitochondria $Ca^{2+}$. (**F-G**) Plots show mean ± SEM of every genotype for each stimulation frequency tested; sample size (larvae) is indicated within the plot for each genotype. For each larva, responses from a 50 second recording from one muscle 1 NMJ between A4-A6 segments were analyzed. Datapoints represent the time between stimulation and peak ΔF/F, or peak ΔF/F and half recovery. Comparisons were analyzed with a mixed-effects two-way ANOVA. Genotypes are *Ib-GAL4, UAS-CEPIA3mt::myc* / *UAS-tdTom::Sec61β*, in either a *WT,* or *Rtnl1$^{18}$* background.

**Video 1. *WT* postsynaptic $Ca^{2+}$ response to low-frequency stimulation.** Evoked postsynaptic $Ca^{2+}$ response to a 1 Hz stimulation. Genotype is *WT; Is-GAL4, Mhc-SynapGCaMP6f / UAS-tdTom::Sec61β.* In this and in all subsequent videos, $Ca^{2+}$ sensor GCaMP is in green, ER marker tdTom::Sec61β in magenta; time is shown in seconds, and stimulation period (when present) as 'STIM"; scale bar 10 µm.

**Video 2. *Rtnl1* mutant postsynaptic $Ca^{2+}$ response to low-frequency stimulation.** Evoked postsynaptic $Ca^{2+}$ response to a 1 Hz stimulation. Genotype is *Rtnl1$^{18}$; Is-GAL4, Mhc-SynapGCaMP6f / UAS-tdTom::Sec61β.*

**Video 3. *WT* postsynaptic miniature $Ca^{2+}$ events.** Spontaneous postsynaptic $Ca^{2+}$ at rest. Genotype is *WT; Is-GAL4, Mhc-SynapGCaMP6f / UAS-tdTom::Sec61β.*

**Video 4. *Rtnl1* mutant postsynaptic miniature $Ca^{2+}$ events.** Spontaneous postsynaptic $Ca^{2+}$ at rest. Genotype is *Rtnl1$^{18}$; Is-GAL4, Mhc-SynapGCaMP6f / UAS-tdTom::Sec61β.*

**Video 5. *WT* cytosolic $Ca^{2+}$ response to stimulation.** Evoked cytosolic $Ca^{2+}$ response to a 30 Hz stimulation. Genotype is *WT; Ib-GAL4, UAS-myr::GCaMP6s / UAS-tdTom::Sec61β.*

**Video 6. *Rtnl1* mutant cytosolic $Ca^{2+}$ response to stimulation.** Evoked cytosolic $Ca^{2+}$ response to a 30 Hz stimulation. Genotype is *Rtnl1$^{18}$; Ib-GAL4, UAS-myr::GCaMP6s / UAS-tdTom::Sec61β.*

**Video 7. *WT* ER $Ca^{2+}$ response to stimulation in Is boutons.** Evoked ER $Ca^{2+}$ response to a 40 Hz stimulation. Genotype is *WT; Is-GAL4, UAS-ER-GCaMP6-210 / UAS-tdTom::Sec61β.*

**Video 8. *Rtnl1* mutant ER $Ca^{2+}$ response to stimulation in Is boutons.** Evoked ER $Ca^{2+}$ response to a 40 Hz stimulation. Genotype is *Rtnl1$^{18}$; Is-GAL4, UAS-ER-GCaMP6-210 / UAS-tdTom::Sec61β.*

**Video 9. *WT* ER $Ca^{2+}$ response to stimulation in Ib boutons.** Evoked ER $Ca^{2+}$ response to a 40 Hz stimulation. Genotype is *WT; Ib-GAL4, UAS-ER-GCaMP6-210 / UAS-tdTom::Sec61β.*

**Video 10. *Rtnl1* mutant ER $Ca^{2+}$ response to stimulation in Ib boutons.** Evoked ER $Ca^{2+}$ response to a 40 Hz stimulation. We provide two video versions, the first with the brightness comparable to the WT Video 9, Genotype is *Rtnl1$^{18}$; Ib-GAL4, UAS-ER-GCaMP6-210 / UAS-tdTom::Sec61β.*

**Video 11. *Rtnl1* mutant ER $Ca^{2+}$ response to stimulation in Ib boutons, brighter image.** A brighter version of Video 10.

**Video 12. *WT* mitochondria $Ca^{2+}$ response to stimulation in Ib boutons.** Evoked mitochondria $Ca^{2+}$ response to a 40 Hz stimulation. Genotype is *WT; Ib-GAL4, UAS-CEPIA3mt / UAS-tdTom::Sec61β.*

**Video 13. *Rtnl1* mutant mitochondria $Ca^{2+}$ response to stimulation in Ib boutons.** Evoked mitochondria $Ca^{2+}$ response to a 40 Hz stimulation. Genotype is *Rtnl1$^{18}$; Ib-GAL4, UAS-CEPIA3mt / UAS-tdTom::Sec61β.*
